# Distinct innate immune states associate with divergent serological outcomes after syphilis treatment

**DOI:** 10.64898/2026.11.15.766608

**Authors:** Qin Wei, Yifan Jin, Yue Zhang, Xinyuan Li, Qihang Zheng, Lu Gan, Wenjing Le, Anlan Hong, Yuanyuan Zhao, Sai Li, Lixia Huang, Futing Liu, Qi Wang, Shaochun Chen, Xiaohong Su, Qianqiu Wang, Xiangsheng Chen, Suying Feng, Wujian Ke, Wenqi Xu, Ning Yan, Wei Chen, Zihai Li, Xiaoyun Yu, Tong Lin, Yetao Wang

## Abstract

The global resurgence of syphilis poses a major public health challenge, even in the era of effective antibiotic therapy. Following treatment, some patients achieve seroreversion, whereas others remain serofast with persistent nontreponemal antibody reactivity. Although nontreponemal antibody titers guide clinical follow-up, it remains unclear how closely they reflect the resolution of inflammation and recovery of immune function. We integrated single-cell transcriptomics, immunophenotyping in clinical samples, and functional assays to characterize circulating monocytes and natural killer (NK) cells across seroreversion and low- and high-titer serofast states, and further examined the impact of HIV coinfection. CD14^+^ monocytes retained inflammatory and CD163-associated scavenging programs after seroreversion, whereas serofast patients showed additional IFN-STAT1 activation in this subset. NK cells from serofast patients exhibited reduced JUN expression and impaired degranulation. JNK inhibition reduced the magnitude and altered the kinetics of NK cell degranulation, IFN-γ production, and amphiregulin (AREG) induction. Among low-titer serofast patients, HIV coinfection was associated with higher mitochondrial complex I gene expression and partially restored NK cell degranulation compared with patients with syphilis alone. Complex I inhibition also reduced NK cell degranulation. These findings suggest that seroreversion does not necessarily indicate immune normalization and identify post-treatment immune alterations not fully captured by conventional serological monitoring, providing a rationale for investigating cellular immune markers as complementary measures of recovery after syphilis treatment.

**GRAPHICAL ABSTRACT:** 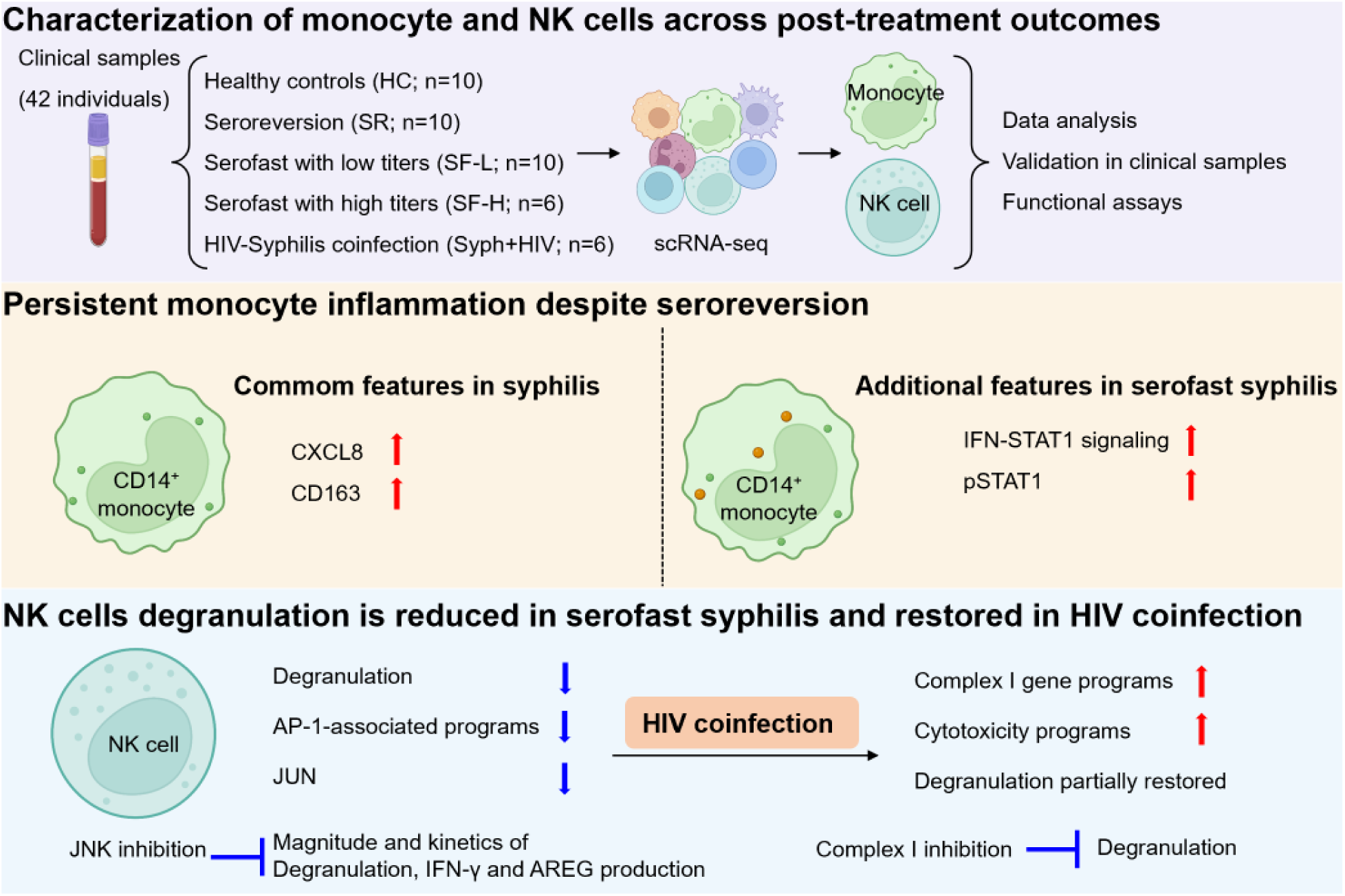

## INTRODUCTION

Syphilis, caused by *Treponema pallidum*, has resurged worldwide despite the availability of effective penicillin therapy. This sexually transmitted infection can persist for years and cause severe neurological and cardiovascular complications if untreated. For 2022, the World Health Organization (WHO) estimated 8.0 million new infections among adults aged 15-49 years (up from 7.1 million in 2020) and approximately 230,000 syphilis-related deaths worldwide^1,2^. Congenital syphilis cases in the United States increased for a 12th consecutive year in 2024^3^, and China reported more than 500,000 syphilis cases in 2025^4^. Rising transmission and associated clinical consequences make syphilis a pressing global public health and social challenge.

A central challenge in syphilis care is determining which patients with persistent seroreactivity after adequate treatment require further intervention^5^. Treatment monitoring relies on nontreponemal tests such as the toluidine red unheated serum test (TRUST) and rapid plasma reagin (RPR) test, which detect antibodies against lipid antigens shared by *T. pallidum* and damaged host cells^6,7^. Some patients achieve seroreversion, whereas others remain serofast, a state not consistently reversed by additional antibiotics^8,9^. Such persistent reactivity may reflect ongoing *T. pallidum* infection, residual antigenic stimulation, unresolved inflammatory injury or sustained antibody production after infection has resolved^5,10^. These possibilities have different implications for retreatment and follow-up but cannot be distinguished by antibody titres alone. Syphilis and human immunodeficiency virus (HIV) share transmission routes and frequently coexist. Although serological responses to syphilis treatment are generally similar in patients with and without HIV, coinfection may be associated with slower declines in nontreponemal antibody titers in some patients and an increased risk of neurological complications, including neurosyphilis^8,11,12^. The cellular immune states associated with these divergent serological outcomes, and their modification by HIV, remain poorly defined. Defining these states could therefore provide a biological basis for interpreting persistent seroreactivity and improving post-treatment risk stratification.

Innate immunity plays a central role in *T. pallidum* clearance and shapes the inflammatory signals that regulate adaptive responses^10,13,14^. Altered innate immune signalling has also been associated with persistent nontreponemal seroreactivity after syphilis treatment^15^. Monocytes and macrophages are key effectors of antibody-mediated clearance, internalizing and degrading opsonized *T. pallidum*^13,14^. NK cells provide an early innate source of interferon-γ (IFN-γ), which enhances this clearance by increasing macrophage CD64 expression and phagocytic uptake^13,14^. *T. pallidum*, in turn, interferes with these innate defenses. For example, its protein Tp47 suppresses macrophage uptake of the bacterium by inducing prostaglandin E2 (PGE2) production16. The pathogen also acquires platelet-derived major histocompatibility complex class I (MHC-I) molecules that trigger KIR2DL3-mediated inhibitory signalling in NK cells^17^. Despite these reciprocal interactions, how the transcriptional states and effector functions of these cell types vary across post-treatment serological outcomes remains poorly defined.

Here, we investigated the innate immune landscape across distinct post-treatment outcomes of syphilis, focusing on circulating monocytes and NK cells. Our analyses revealed distinct monocyte and NK-cell states associated with these outcomes and identified HIV coinfection-associated alterations in NK cells. These findings highlight distinct post-treatment innate immune states that are not fully reflected by conventional serological measures, providing new insight into immune recovery after syphilis treatment.

## RESULTS

### Persistent monocyte inflammation with subset-specific scavenging and STAT1 activation across syphilis treatment outcomes

To investigate whether monocyte and NK-cell states were associated with distinct post-treatment serological outcomes, we performed single-cell RNA sequencing (scRNA-Seq) of peripheral blood mononuclear cells (PBMCs) from 36 individuals, including healthy controls (HC; n=10), patients with syphilis who achieved seroreversion (SR; n=10), and patients who remained serofast with low (SF-L; TRUST <1:8; n=10) or high (SF-H; TRUST ≥1:8; n=6) titers. Seroreversion was defined as conversion of the TRUST result from reactive to nonreactive after adequate treatment, and serofast status was defined as persistent TRUST reactivity for more than 1 year after treatment. All patients with syphilis had nonreactive cerebrospinal fluid (CSF) TRUST results and no clinical evidence of neurosyphilis. Key findings from the single-cell analysis were validated in additional clinical samples, and selected alterations were further examined for their functional consequences. The study design and participant characteristics are summarized in Figure S1A and Table S1.

We first re-clustered the monocyte and dendritic-cell compartments identified by unsupervised graph-based clustering (Figures S1B and S1C), in which CD14^+^ and CD16^+^ monocytes together accounted for approximately 90% of total myeloid cells, with the remainder comprising cDC1, cDC2, and plasmacytoid dendritic cells (pDCs) (Figures 1A–1C). Among these populations, SF-L patients showed an increased proportion of CD14^+^ monocytes and a decreased proportion of CD16^+^ monocytes compared with HC (Figure 1C). Principal component analysis (PCA) showed that transcriptional variation was primarily driven by cell identity rather than post-treatment serological outcome (Figure 1D), prompting us to examine outcome-associated changes within each cell type. Gene-level comparison revealed distinct patterns of differentially expressed genes (DEGs) across myeloid subsets (Figure 1E). CD14⁺ monocytes showed relatively high numbers of DEGs in pairwise comparisons among HC, SR, and SF-H, whereas CD16⁺ monocytes showed a markedly higher number of DEGs between HC and SF-H than in other comparisons. Both monocyte subsets showed relatively few DEGs between SF-L and SF-H, suggesting limited transcriptional divergence between the two serofast states (Figure 1E).

**Figure 1.**
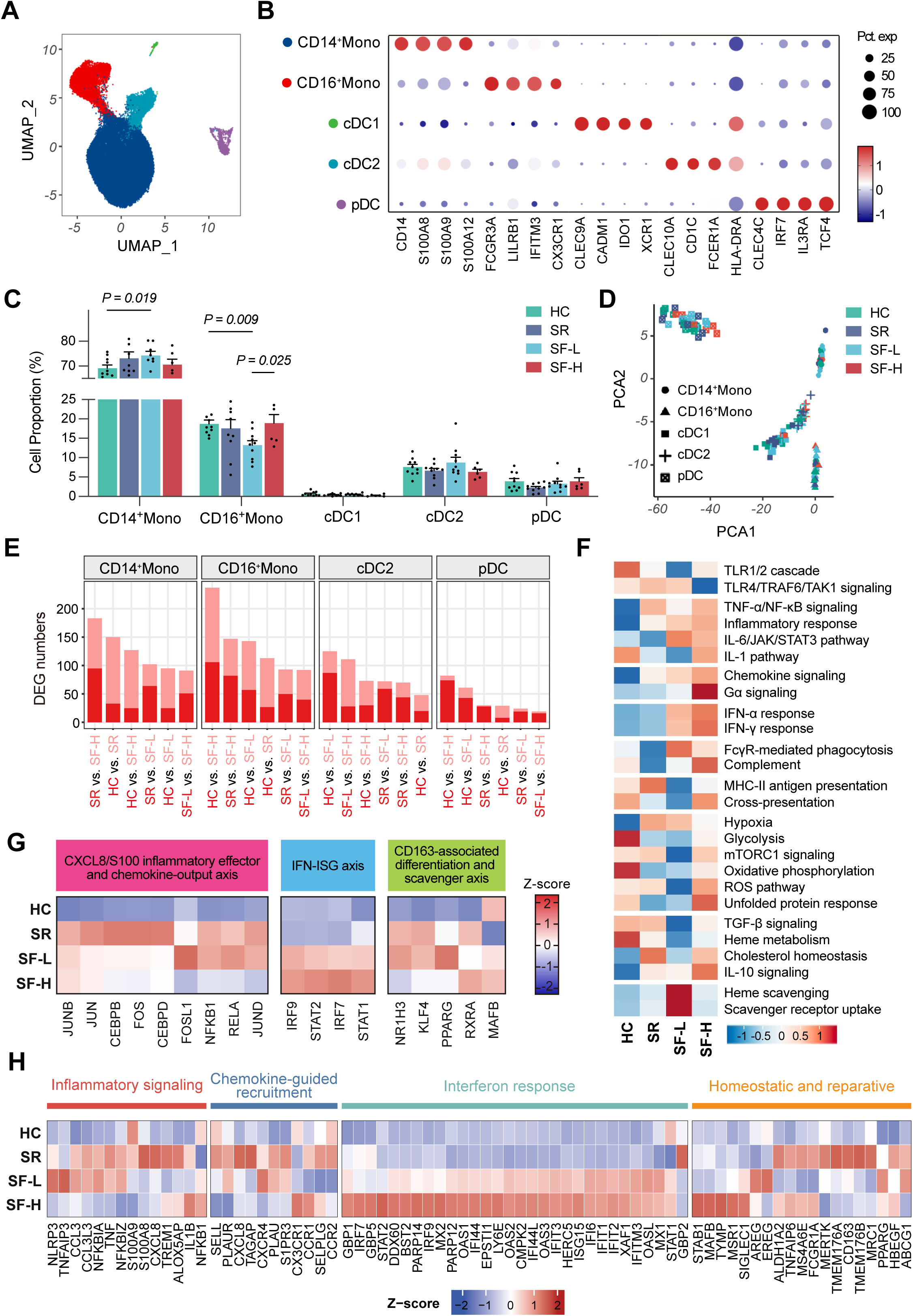
Shared inflammatory and scavenging programs and serofast-associated IFN-STAT1 responses in CD14⁺ monocytes. (A) UMAP of myeloid cells, colored by five annotated subsets. (B) Dot plot of representative myeloid subset markers. (C) Proportions of the indicated subsets among total myeloid cells in HC (n=10), SR (n=10), SF-L (n=10), or SF-H (n=6). Each dot represents one donor; Mean ± standard error of the mean (SEM). (D) PCA of myeloid transcriptional profiles, colored by clinical group, with symbols indicating cell subsets. (E) Numbers of DEGs across pairwise group comparisons in the indicated myeloid subsets. (F) Heatmap of Z-scored enrichment scores for the indicated pathways in CD14^+^ monocytes across the four clinical groups. (G) Heatmaps of module-related TF activities in CD14⁺ monocytes across the four clinical groups. (H) Heatmaps of Z-scored expression of clinical group-specific marker genes in CD14⁺ monocytes across the four clinical groups, annotated by functional module.

To further define monocyte transcriptional programs associated with distinct post-treatment serological outcomes, we first performed pathway enrichment analysis in CD14^+^ monocytes across the HC, SR, SF-L, and SF-H groups (Figure 1F). Several pathways with established roles in anti-treponemal immunity showed outcome-dependent enrichment, including Fcγ receptor-mediated phagocytosis in SF-L, complement activation in SF-H, MHC class II antigen presentation in HC and SR, and cross-presentation in HC and SF-H^13,18–20^. More importantly, several broader transcriptional patterns were associated with post-treatment serological outcomes. TNF-NF-κB signaling and inflammatory-response programs were broadly increased across the syphilis groups relative to HC, suggesting a shared inflammatory state after treatment. In contrast, type I and type II interferon-response programs were selectively enriched in both serofast groups, whereas heme scavenging and scavenger-receptor-mediated ligand uptake were most prominently enriched in SF-L monocytes (Figure 1F).

The transcriptional programs underlying these patterns were further characterized by assessing transcription factor activity and group-enriched gene expression (Figures 1G and 1H). The two analyses showed concordant patterns across clinical groups. At the gene level, inflammatory genes, including CXCL8, CCL3, IL1B, NLRP3, and TNF, together with scavenging-, homeostasis-, and repair-associated genes, including CD163, MRC1, MAFB, FCGR1A, AREG, EREG, and HBEGF, were preferentially expressed across the SR, SF-L, and SF-H groups. In contrast, IFN-STAT1-associated genes, including IRF7, STAT1, MX1, MX2, IFIT1, IFIT2, IFIT3, and IFI44, were preferentially expressed in the serofast groups (Figure 1H).

Although scavenger-related pathways were also enriched in SF-L CD16^+^ monocytes, CD163-associated scavenging genes and homeostatic/repair-associated genes, including CD163, AREG, EREG, and HBEGF, were not preferentially expressed in this subset (Figures S2A and S2B). Thus, CD163-associated scavenging and homeostatic/repair transcriptional features were more prominent in CD14^+^ than in CD16^+^ monocytes.

To determine whether the outcome-associated monocyte transcriptional programs identified above were reflected at the protein level, we performed flow-cytometric validation in additional clinical samples, focusing on markers of the scavenging, inflammatory, and IFN-STAT1 programs (Figures 2A and 2B). In CD14⁺ monocytes, CD163 the mean fluorescence intensity (MFI) for CD163 and CXCL8 were increased in SR, SF-L, and SF-H compared with HC (Figures 2C and 2D), whereas pSTAT1 expression was selectively elevated in SF-L and SF-H (Figure 2E). Plasma EREG concentrations were also increased in SF-L (Figure 2F).

**Figure 2.**
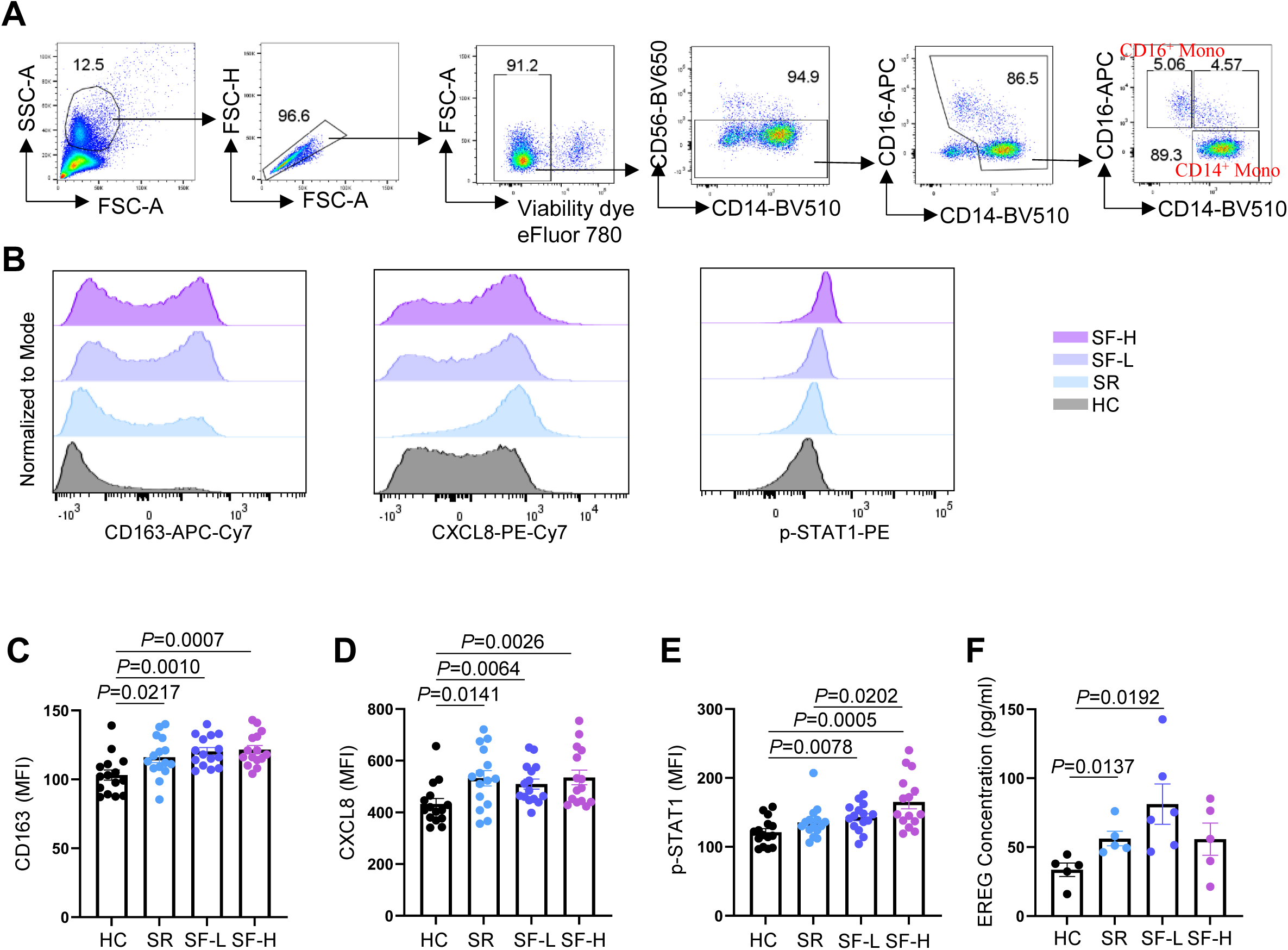
Persistent inflammatory and scavenging features after seroreversion and serofast-associated STAT1 activation in CD14⁺ monocytes. (A) Flow-cytometric gating for identifying CD14⁺ and CD16⁺ monocytes in PBMCs. (B) CD14⁺ monocytes identified in (A) were stained with antibodies against CD163, CXCL8, and pSTAT1. (C-E) MFI of CD163 (C), CXCL8 (D), and p-STAT1 (E) in CD14⁺ monocytes (n=15 per group). (F) Plasma EREG concentrations measured by ELISA (n=5 per group for HC, SR, and SF-H, and n=6 for SF-L). For (C, and F), two-tailed unpaired t-tests. For (D, and E), two-tailed Mann-Whitney U test. Each dot represents an individual donor. Data are mean ± SEM. ns, not significant.

In CD16⁺ monocytes, CXCL8 expression was similarly elevated across all three post-treatment syphilis groups (Figure S2C). Unlike in CD14^+^ monocytes, CD163 expression remained unchanged across groups, whereas pSTAT1 was elevated in all three post-treatment groups, including SR (Figures S2D and S2E). Together, these findings support persistent inflammatory features after seroreversion in both monocyte subsets, with distinct patterns of CD163 expression and STAT1 activation.

### Reduced AP-1-associated transcription in serofast NK cells and distinct receptor profiles across clinical groups

To characterize NK cell transcriptional states associated with post-treatment serological outcomes, we reclustered NK cells using unsupervised graph-based clustering. Consistent with previous studies^21–23^, NK cells were mainly grouped into CD56^dim^ and CD56^hi^ subsets (Figures 3A and 3B). The relative proportions of these populations did not differ significantly across clinical groups (Figure 3C). PCA showed that NK cell transcriptional profiles segregated primarily by subset identity rather than post-treatment serological outcome (Figure 3D), and the CD56^dim^ and CD56^hi^NK cells were therefore analyzed separately. Differential expression analysis revealed substantially more DEGs in CD56^dim^ than in CD56^hi^NK cells across clinical group comparisons, indicating greater outcome-associated transcriptional variation (Figure 3E). Within CD56ᵈⁱᵐNK cells, the largest numbers of DEGs were observed when HC was compared with either SF-H or SF-L, with fewer DEGs detected between the two serofast groups (Figure 3E).

**Figure 3.**
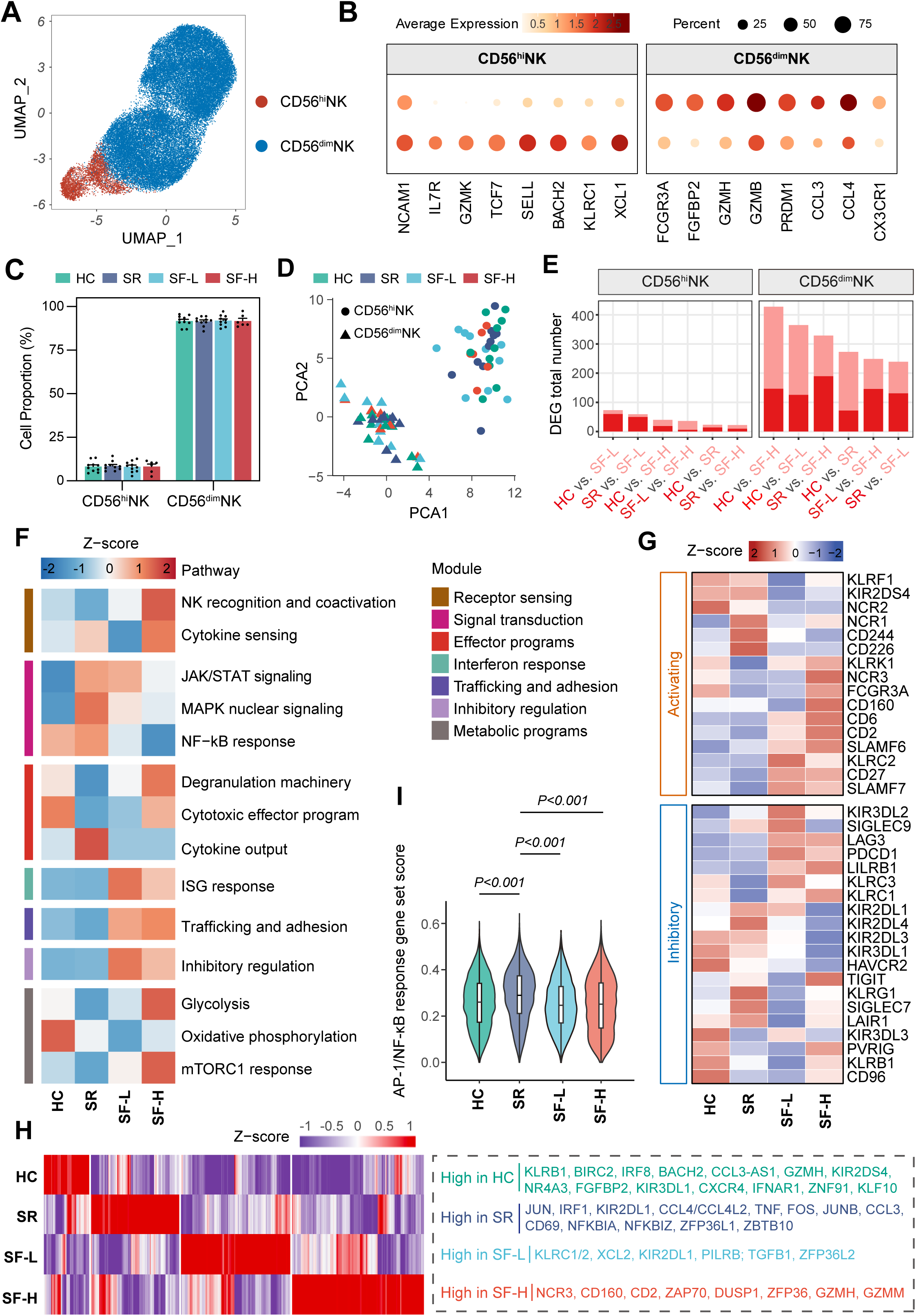
Reduced AP-1-associated transcription and distinct NK cell receptor profiles across clinical groups. (A) UMAP of NK subsets, colored by two annotated subsets. (B) Dot plot of representative NK cell subset markers. (C) Proportions of the indicated subsets among total NK cells in HC (n=10), SR (n=10), SF-L (n=10), or SF-H (n=6). Each dot represents one donor; Mean ± SEM. (D) PCA of transcriptional profiles for NK cells, colored by clinical group, with symbols indicating cell subsets. (E) Numbers of DEGs across pairwise group comparisons in the indicated NK subsets. (F) Heatmap of Z-scored enrichment scores for the indicated pathways in CD56^dim^NK cells across the four clinical groups. (G) Heatmaps of Z-scored expression of NK receptors in CD56^dim^ NK cells across the four clinical groups. (H) Heatmaps of Z-scored expression of group-specific marker genes in CD56^dim^NK cells across the four clinical groups, annotated by functional module. (I) AP-1/NF-κB response gene set score in CD56^dim^NK cells across the four clinical groups.

We first examined pathway-level differences in CD56^dim^NK cells across the four clinical groups. (Figure 3F). Notably, MAPK nuclear signaling, a major upstream regulator of AP-1 activity^24,25^, progressively decreased from SR to SF-L and SF-H, accompanied by a similar reduction in NF-κB responses. Although degranulation, cytotoxic-effector, and cytokine-output programs showed heterogeneous patterns across groups, ISG responses, trafficking and adhesion, and inhibitory regulation were more strongly enriched in both serofast states (Figure 3F). NK cell receptor-associated pathways also varied across groups (Figure 3G). HC and SR showed relatively higher expression of receptors such as KLRF1 and KIR2DS4, with NCR1, CD244, and CD226 additionally enriched in SR. In contrast, serofast groups showed increased expression of several activating and inhibitory receptors, including CD160, CD6, CD2, SLAMF6, LAG3, PDCD1, and LILRB1, while TIGIT was particularly enriched in SF-H (Figure 3G). These analyses revealed distinct patterns of AP-1-associated signaling and NK cell receptor expression across serological outcomes.

Analysis of group-enriched genes further showed that SR NK cells preferentially expressed the AP-1 factors JUN, FOS, and JUNB, together with NF-κB-associated and inflammatory genes including NFKBIA, NFKBIZ, TNF, CCL3, and CCL4 (Figure 3H), consistent with higher AP-1/NF-κB response scores in SR than in both serofast groups (Figure 3I). SF-L NK cells preferentially expressed TGFB1, XCL2, and ZFP36L2, while SF-H NK cells were characterized by higher expression of signaling- and effector-related genes, including ZAP70, DUSP1, ZFP36, GZMH, and GZMM (Figure 3H).

In CD56^hi^NK cells, ISG responses were enriched in both serofast groups, whereas NF-κB responses were highest in SR and reduced in SF-L and SF-H (Figure S3A), consistent with the patterns observed in CD56ᵈⁱᵐNK cells. CD56^hi^NK cells also showed marked remodeling of the receptor repertoire, with SR enriched for KLRF1, NCR2, and CD226, SF-L for KLRC2, SLAMF6, and CD160, and SF-H for NCR1, NCR3, FCGR3A, SLAMF7, TIGIT, KIR3DL2, LILRB1, and SIGLEC9 (Figure S3B). Thus, receptor remodeling was shared across NK cell subsets, although the specific receptor profiles were subset dependent.

### Reduced JUN expression and degranulation in serofast NK cells with subset-specific receptor changes

To determine whether the AP-1-associated transcriptional changes and receptor remodeling identified by scRNA-seq were reflected at the protein level, we examined JUN and surface receptor expression in CD56^dim^ and CD56^hi^NK cells by flow cytometry (Figures 4A and 4B). JUN expression was reduced in SF-H compared with HC and SR in both NK cell subsets, with an additional reduction in SF-L compared with SR in CD56^hi^NK cells (Figures 4C and 4D), consistent with the reduced AP-1-associated transcriptional activity observed in serofast NK cells (Figures 3F, 3H and 3I).

**Figure 4.**
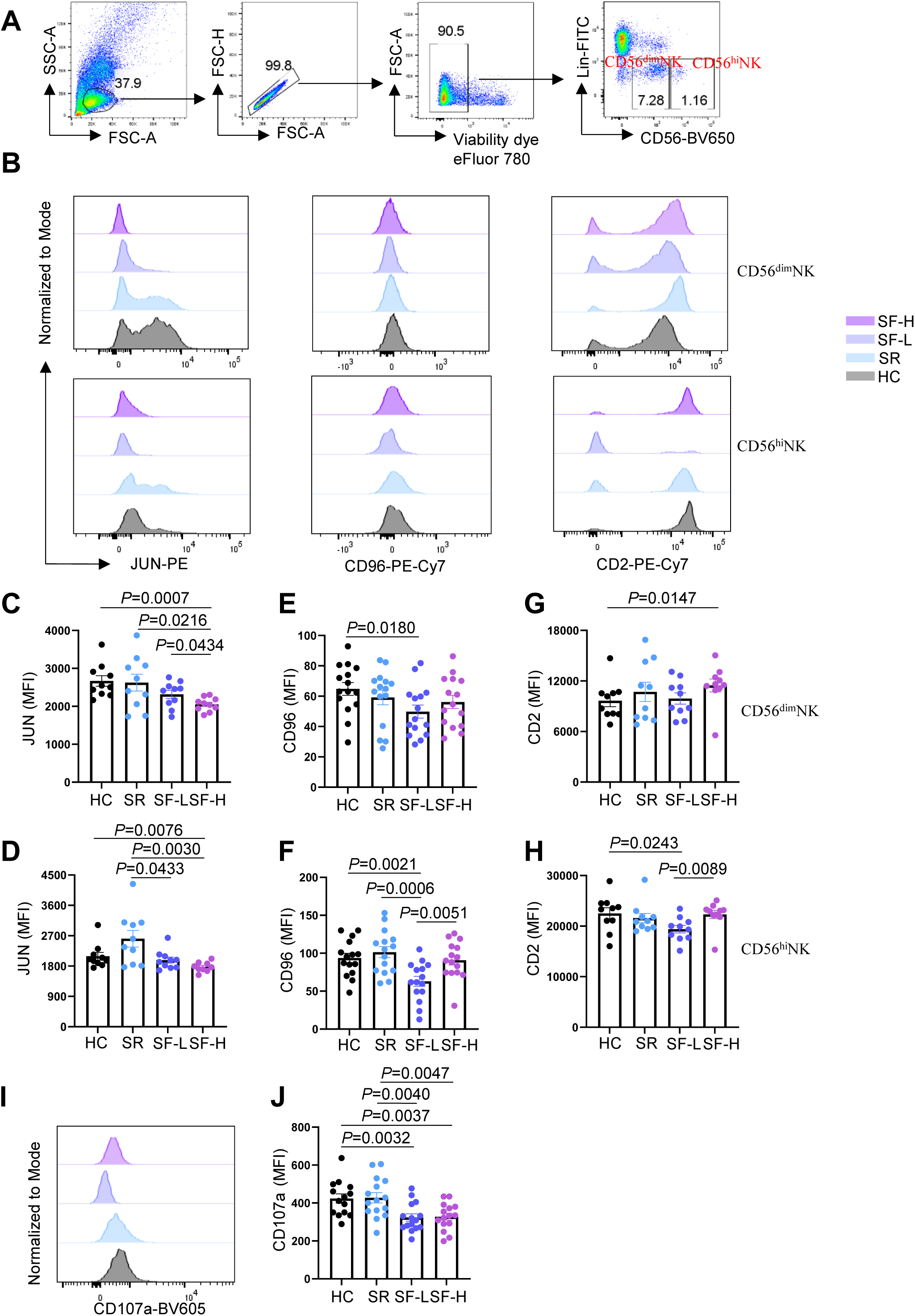
Reduced JUN expression and degranulation in serofast NK cells and differential CD96 and CD2 expression across clinical groups. (A) Flow-cytometric gating for CD56^dim^ and CD56^hi^NK cells in PBMCs. The lineage (Lin) antibody cocktail included antibodies against CD3, CD4, TCRαβ, TCRγδ, CD19, CD20, CD22, CD34, FcεRIα, CD11c, CD303, CD123, CD1a, and CD14. (B) CD56^dim^ and CD56^hi^NK cells identified in (A) were stained with antibodies against JUN, CD96, and CD2. (C-H) MFI for JUN (C, D), CD96 (E, F), and CD2 (G, H) in CD56^dim^ and CD56^hi^NK cells (For C, G, and H, n=10 per group; for E and F, n=15 per group; for D, n=10 each for HC, SR and SF-L, and n=9 for SF-H). (I) Total NK cells (Lin⁻CD56⁺) were surface stained with antibody against CD107a following K562 stimulation. (J) CD107a MFI in total NK cells (n=14 for HC, n=15 each for SR, SF-L and SF-H). For (C, F, and J), two-tailed unpaired t-tests, for (D, E, G, and H), two-tailed Mann-Whitney U test. Each dot represents an individual donor. Data are mean ± SEM; ns, not significant.

NK cell surface receptor expression also varied across clinical groups. CD96, an immunoregulatory NK cell receptor^26^, showed reduced expression in SF-L compared with HC in CD56^dim^NK cells and with HC, SR, and SF-H in CD56^hi^NK cells (Figures 4E and 4F). CD2, an adhesion and costimulatory receptor^27^, was expressed at higher levels in CD56^dim^NK cells from SF-H than HC, but at lower levels in CD56^hi^NK cells from SF-L than HC and SF-H (Figures 4G and 4H). We further assessed NK cell degranulation following K562 stimulation and observed reduced surface CD107a expression in both SF-L and SF-H compared with HC and SR (Figures 4I and 4J).

Together, flow-cytometric analyses revealed reduced JUN expression and impaired NK cell degranulation in serofast syphilis, alongside subset specific changes in surface receptor expression across serological outcomes.

### JNK signaling regulates the magnitude and kinetics of NK cell effector responses

Given the reduced JUN expression in serofast NK cells, we investigated its potential functional relevance by inhibiting JNK, an upstream regulator of JUN activity. JNK inhibition reduced JUN protein level and attenuated NK cell degranulation and IFN-γ production (Figures 5A–5D, 5F and 5G). It also reduced the expression of AREG (Figures 5I and 5J), a tissue repair-associated immunoregulatory factor expressed by NK cells^28,29^. Consistently, JUN⁺NK cells exhibited higher CD107a, IFN-γ, and AREG expression than their JUN⁻ counterparts (Figures 5E, 5H, and 5K). Furthermore, the proportion of JUN⁺NK cells was positively correlated with the proportions of NKp30⁺, AREG⁺, and IFN-γ⁺NK cells (Figures 5L–5N).

**Figure 5.**
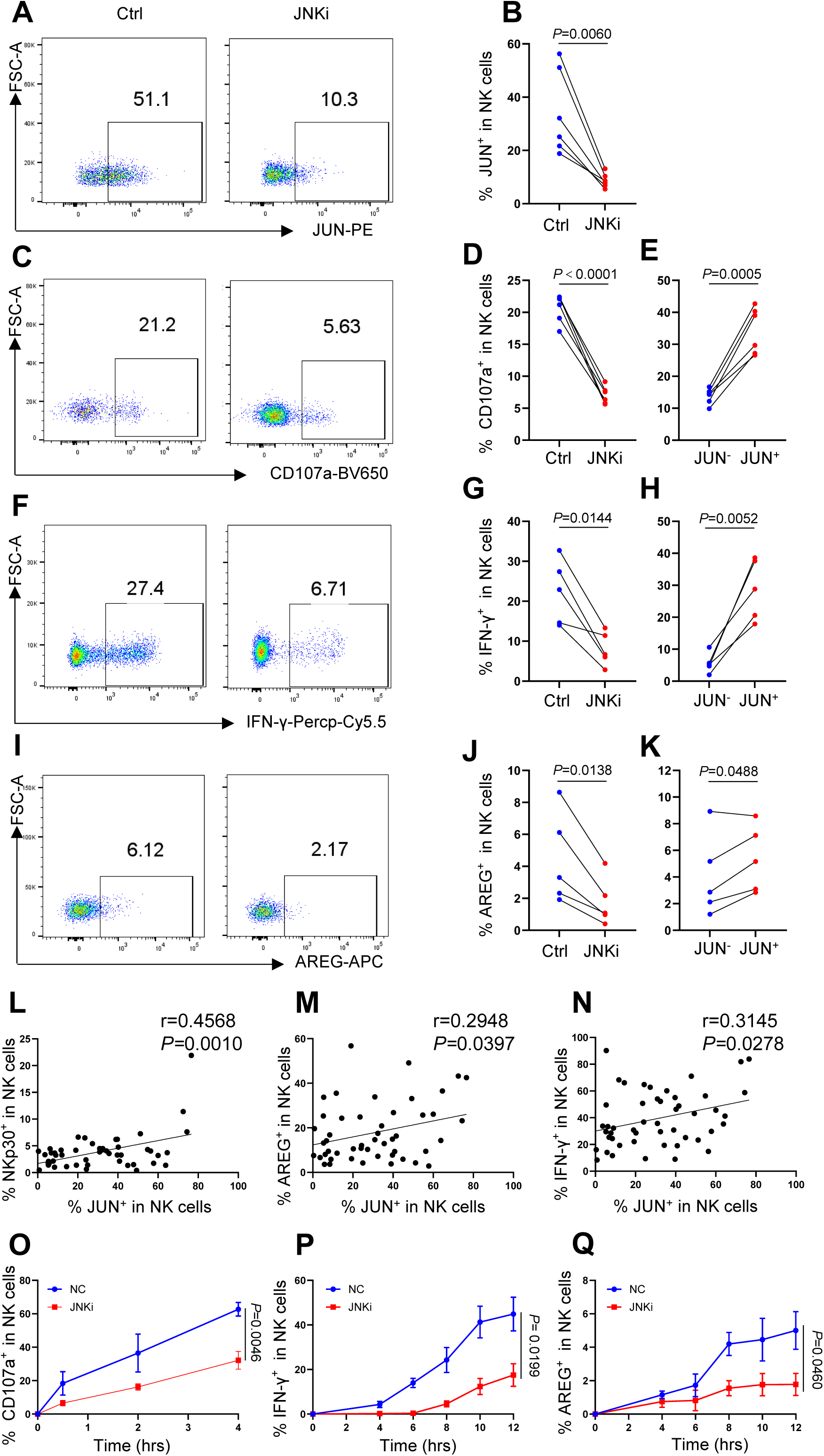
JNK inhibition reduces the magnitude and alters the kinetics of NK cell effector responses. (A, B) PBMCs were untreated or treated with JNK-IN-8 (JNKi; 10 μM) for 16 hrs. JUN expression in NK cells was assessed by flow cytometry (A), and the percentage of JUN⁺ NK cells was quantified (B, n=6). (C-E) PBMCs treated as in (A) were cocultured with K562 cells. NK cell surface CD107a expression was assessed by flow cytometry (C), and the percentage of CD107a^+^NK cells was quantified (D, n=6); CD107a^+^ cell percentages were compared between JUN^−^ and JUN^+^NK cells from the untreated group (E, n=6). (F-H) PBMCs treated as in (A) were stimulated with IL-12 (10 ng/mL), IL-15 (50 ng/mL), and IL-18 (50 ng/mL), NK cell IFN-γ expression was assessed by flow cytometry (F), and the percentage of IFN-γ^+^NK cells was quantified (G, n=5); IFN-γ^+^ cell percentages were also compared between JUN^−^ and JUN^+^NK cells from the untreated group (H, n=5). (I-K) PBMCs treated as in (A) were stimulated with dexamethasone (100 nM), NK cell AREG expression was assessed by flow cytometry (I), and the percentage of AREG^+^NK cells was quantified (J, n=5); AREG^+^ cell percentages were also compared between JUN^−^ and JUN^+^NK cells from the untreated group (K, n=5). (L-N) Pearson correlations between JUN^+^ NK cell percentages and NKp30^+^ (L), AREG^+^ (M), and IFN-γ^+^ (N) NK cell percentages. (O-Q) Percentages of CD107a^+^ (O, n=4), IFN-γ^+^ (P, n=4), and AREG^+^ (Q, n=4) NK cells at the indicated time points under the corresponding stimulation conditions described in (C), (F), and (I), respectively. For (B, D, E, G, H, J, and K), two-tailed paired t-tests. For (L-N), Pearson’s correlation. For (O-Q), two-way ANOVA. Data are mean ± SEM; ns, not significant.

Time-course analysis showed that JNK inhibition delayed NK cell degranulation and the induction of IFN-γ and AREG following stimulation (Figures 5O–5Q), suggesting that JNK-JUN-associated signaling also influences the kinetics of NK cell effector responses. These data support a potential link between reduced JUN expression and impaired NK cell degranulation in patients with serofast syphilis.

### HIV coinfection is associated with selective mitochondrial complex I gene upregulation and enhanced cytotoxic programs in serofast NK cells

Given that HIV coinfection has been associated with altered serological responses to syphilis^30,31^, we investigated whether HIV further reshapes NK cell states. To minimize potential confounding by differences in post-treatment syphilis serology and given the availability of clinical samples, we restricted both syphilis groups to individuals with low-titer serofast status (SF-L; TRUST <1:8). We compared CD56^hi^ and CD56^dim^NK cells among healthy controls (HC; n=10), patients with syphilis alone (Syph; n=10), and ART-treated patients with HIV-syphilis coinfection (Syph+HIV; n=6) (Figure 6A). PCA again showed that samples segregated primarily by NK cell subset rather than clinical group (Figure 6B). Differential expression analysis revealed substantially more DEGs in CD56^dim^ than in CD56^hi^NK cells across clinical-group comparisons (Figure 6C).

**Figure 6.**
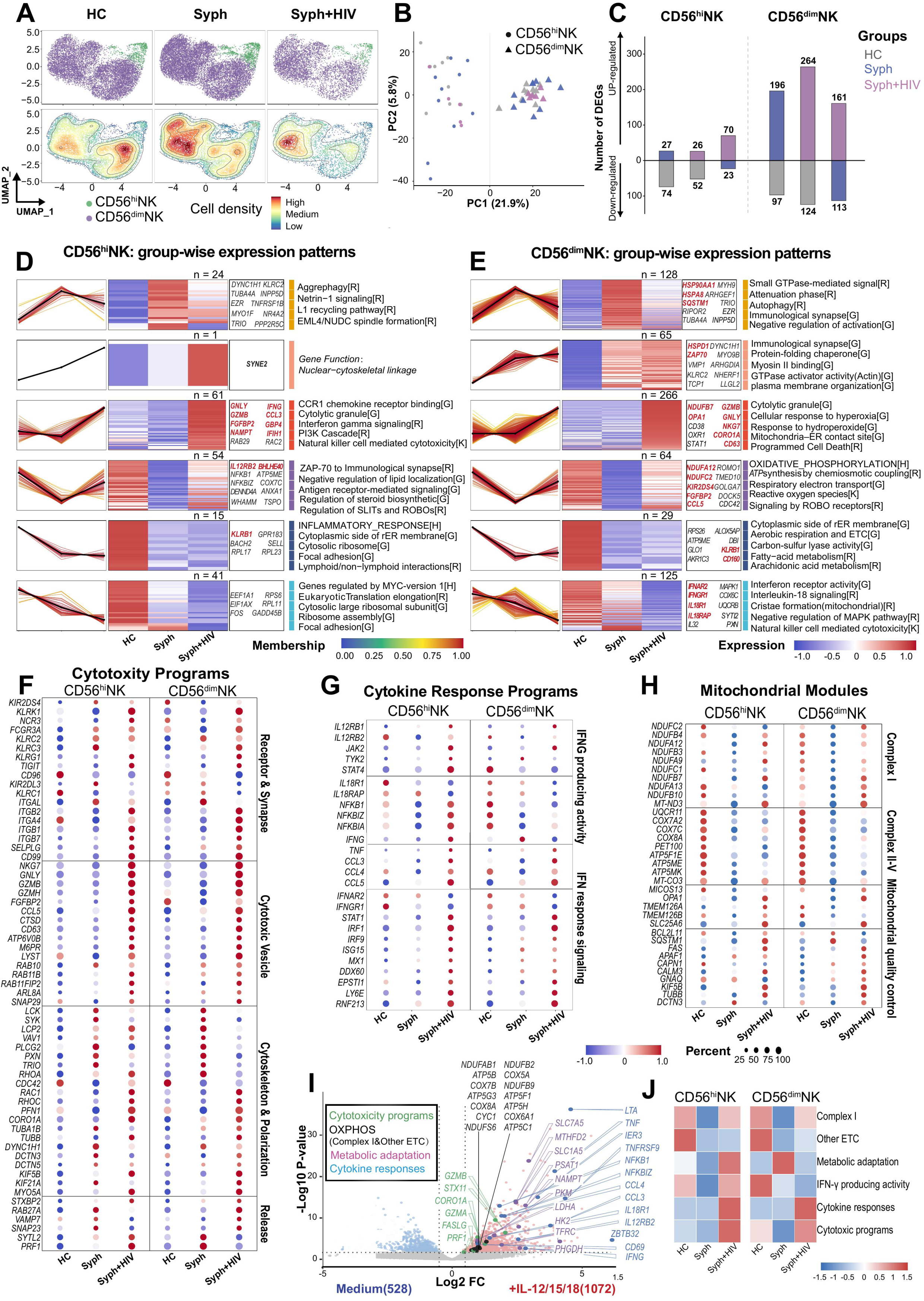
HIV coinfection-associated mitochondrial and effector programs in NK cells from serofast patients. (A) UMAP of CD56^hi^ and CD56^dim^NK cells across HC, Syph, and HIV + Syph, colored by NK cell subset and cell density (HC, n=10; Syph, n=10; Syph+HIV, n=6). (B) PCA of NK cell expression profiles. (C) Numbers of upregulated and downregulated genes in indicated clinical-group comparisons within each NK cell subset (adjusted *P*<0.05, |log₂FC|≥0.25, and pct.1 or pct.2≥0.25). (D, E) Six DEG expression patterns across clinical groups in CD56^hi^ (D) and CD56^dim^NK cells (E). Line plots show Z-scored gene expression patterns (colored by membership score; black lines, the mean standardized expression of genes within each cluster), and heatmaps show relative expression. Selected DEGs and enriched pathways are indicated (R, Reactome; G, Gene Ontology; K, KEGG; H, Hallmark). (F-H) Dot plots of genes associated with mitochondrial modules (F), cytokine response programs (G), and cytotoxicity programs (H). (I) Volcano plot highlighting genes upregulated in IL-12, IL-15 and IL-18-stimulated NK cells relative to unstimulated counterparts (log_2_(fold change, FC)>0.6, *P*<0.01, determined by DESeq2, n=3). (J) Heatmap of AUCell scores for selected gene sets.

To characterize transcriptional changes associated with HIV coinfection, we classified differentially expressed genes across HC, Syph, and Syph+HIV into six expression patterns within each NK cell subset (Figures 6D, 6E, S4A, and S4B). Compared with Syph, Syph+HIV showed increased expression of cytotoxicity- and granule-associated genes (GNLY, GZMB, FGFBP2, CD63, NKG7 and CORO1A)^32–35^ and IFN-response genes (RNF213, GBP4, IFIH1, CCL3, CCL4, CCL5, STAT1, IRF1, ISG15, EIF2AK2, LY6E, DDX60, UBE2L6, and SAMD9L)^36–39^, across the two NK cell subsets (Figures 6D–6G). Notably, mitochondrial complex I genes (NDUFA12, NDUFB7, and NDUFC2) showed reduced expression in Syph relative to HC but were partially restored in Syph+HIV. In contrast, genes encoding subunits of OXPHOS complexes III (UQCR11), IV (COX7A2, COX7C, and COX8A), and V (ATP5F1E, ATP5MC3, and ATP5MK) remained lower than in HC (Figure 6H)^40,41^.

Our previous work linked HIV-induced increases in inflammatory cytokines to altered NK cell activation^23^. We therefore examined whether the transcriptional changes observed in Syph+HIV NK cells resembled those induced by inflammatory cytokines, using RNA-seq profiles of NK cells stimulated with IL-12, IL-15, and IL-18^28^. Cytokine stimulation increased the expression of genes associated with cytokine responses (IFNG, TNF, CCL3, and CCL4), cytotoxicity programs (GZMA, GZMB, and PRF1), metabolic adaptation (SLC7A5, NAMPT, and HK2)^42–44^, and oxidative phosphorylation (NDUFAB1, NDUFS6, NDUFB9, COX7B, and ATP5B) (Figure 6I), features that partially overlapped with those observed in Syph+HIV NK cells (Figures 6D–6H). Gene set scoring further showed that metabolic adaptation and IFN-γ-producing activity scores were higher in Syph+HIV than in Syph only in CD56^hi^NK cells, whereas complex I, IFN-response signaling, and cytotoxicity program scores were higher across both NK cell subsets (Figure 6J).

### Mitochondrial complex I is linked to NK cell degranulation in syphilis and HIV coinfection

We next examined correlations among NK cell metabolic and effector program scores (Figures 7A and 7B).To determine whether the metabolic transcriptional changes observed in Syph+HIV NK cells were associated with effector programs, we examined correlations between metabolic and effector gene-set scores in CD56^hi^ and CD56^dim^NK cells (Figures 7A and 7B). IFN-γ producing activity was positively correlated with metabolic adaptation scores in both NK cell subsets, whereas its association with complex I was not consistently significant (Figures 7A and 7B) across subsets. At the protein level, IFN-γ production did not differ among clinical groups (Figure S4C). In contrast, complex I and cytotoxicity gene-set scores were positively correlated in both NK cell subsets, suggesting a consistent association between complex I-related transcription and cytotoxicity-associated programs.

**Figure 7.**
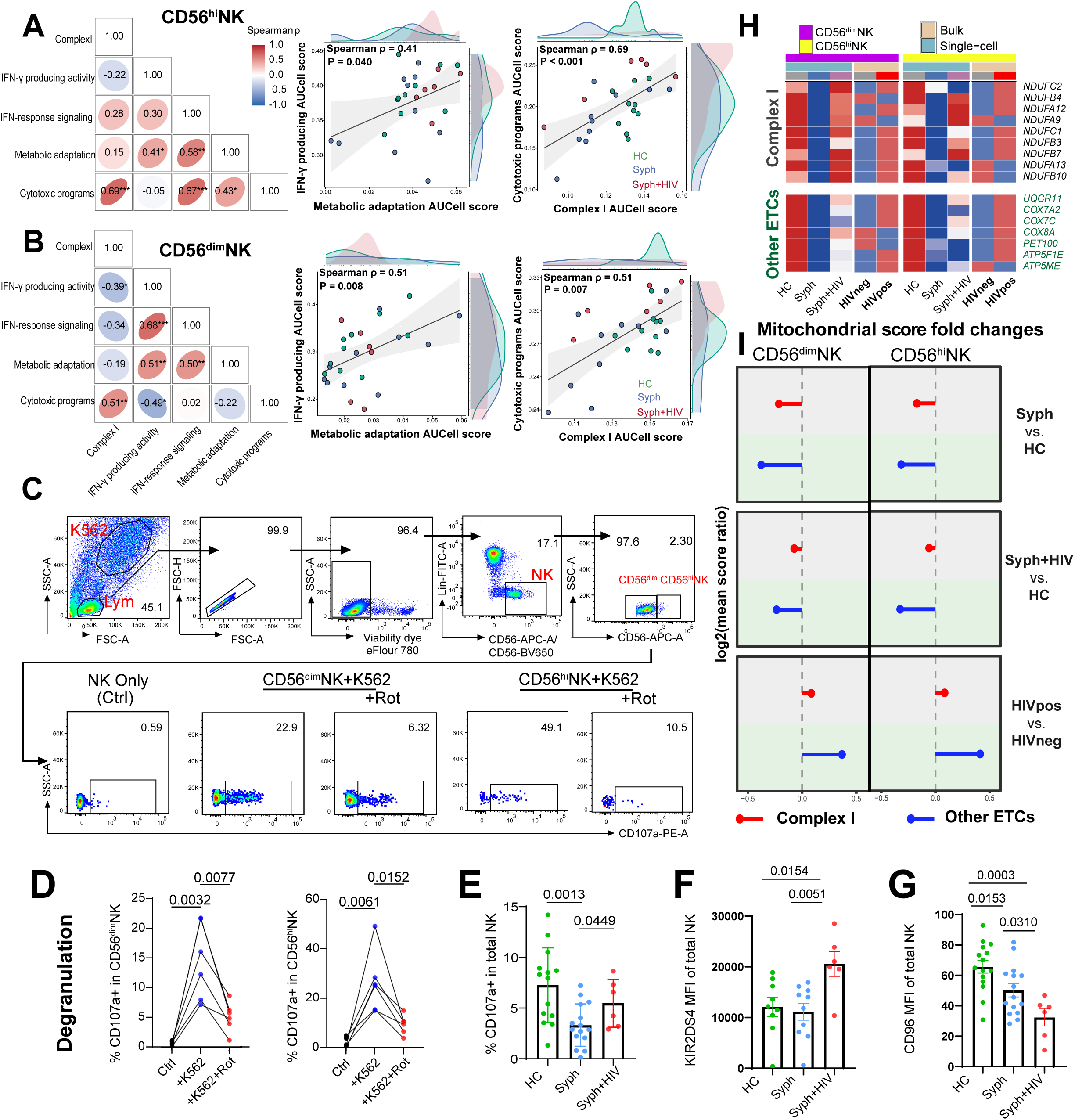
Mitochondrial complex I-associated transcription and NK cell degranulation in syphilis and HIV coinfection. (A, B) Spearman correlations among AUCell scores for selected metabolic and effector gene sets in CD56^hi^ (A) and CD56^dim^NK cells (B). Scatter plots show correlations between metabolic adaptation and IFN-γ production scores and between complex I and cytotoxicity scores. Each dot represents one donor. ρ, Spearman’s rank correlation coefficient; *P* values were calculated using two-sided Spearman’s rank correlation tests. (C) PBMCs were left untreated or treated with rotenone (Rot, 300nM) for 16 h, followed by 4 h of K562 coculture. K562-induced surface CD107a expression in NK cells was assessed by flow cytometry. (D) Percentages of CD107a^+^ cells in indicated NK subsets in (C) (n=6). (E-G) Percentage of CD107a⁺NK cells following K562 stimulation (HC, n=14; Syph, n=15; Syph+HIV, n=6) (E) and MFI of KIR2DS4 (HC, n=9; Syph, n=10; Syph+HIV, n=6) (F) and CD96 (HC, n=15; Syph, n=15; Syph+HIV, n=6) (G) in total NK cells. (H) Heatmaps of complex I and other ETC gene expression in NK cell subsets from HC, Syph, and Syph+HIV, and from HIV-positive and HIV-negative individuals without syphilis. (I) log_2_ ratios of mean mitochondrial gene set scores in NK cell subset in the indicated comparisons. For (D), two-tailed paired t-test, for (E), two-tailed Mann-Whitney U test, data are mean ± SEM, for (F-G), two-tailed unpaired t-test. Each dot represents an individual donor.

Inhibition of mitochondrial complex I with rotenone (Rot) reduced K562-induced degranulation in both CD56^dim^ and CD56^hi^NK cells (Figures 7C and 7D). In clinical samples, the proportion of CD107a^+^NK cells was lower in Syph than HC but partially restored in Syph+HIV (Figure 7E), consistent with the corresponding changes in complex I-associated transcriptional scores (Figure 6J). HIV coinfection was also associated with increased expression of the activating receptor KIR2DS4^45^ and reduced CD96 expression (Figures 7F and 7G), indicating additional receptor remodeling accompanying these functional changes. Together, these findings support a role for mitochondrial complex I in regulating NK cell degranulation and suggest that increased complex I expression may contribute to the partial recovery of this function in HIV-coinfected patients.

To assess whether HIV infection alone was associated with similar changes in complex I, we reanalyzed our previously published bulk RNA-seq data from sorted CD56^dim^ and CD56^hi^NK cells from HIV-negative and ART-treated HIV-positive individuals^22^. HIV infection alone was associated with higher complex I gene expression and corresponding module scores in both NK cell subsets, paralleling the changes observed in Syph+HIV relative to Syph (Figures 7H and 7I). In contrast, genes encoding subunits of other mitochondrial oxidative phosphorylation (OXPHOS) complexes did not show similarly consistent changes across the two comparisons (Figures 7H and 7I). These parallel patterns suggest that HIV infection itself may contribute to the increased complex I expression observed in Syph+HIV, although additional effects of coinfection cannot be excluded.

## DISCUSSION

Divergent serological outcomes following syphilis treatment raise important questions about the extent of immune recovery and the mechanisms underlying persistent seroreactivity. By integrating single-cell transcriptomics, immunophenotyping, and functional assays, we identified distinct monocyte and NK cell states associated with these outcomes. Two main findings emerged: first, monocytes retained inflammatory and CD163-associated scavenging programs even after seroreversion, whereas serofast patients exhibited additional IFN-STAT1 activation; second, serofast status was associated with reduced NK cell JUN expression and degranulation, while HIV coinfection partially restored degranulation alongside increased mitochondrial complex I gene expression. Functional experiments further implicated JNK signaling and complex I in NK cell effector regulation. Together, these findings reveal incomplete innate immune recovery after seroreversion and distinct monocyte and NK cell alterations associated with serofast status, with NK cell function further shaped by HIV coinfection.

Monocyte alterations remained evident even after seroreversion. Compared with HC, CD14⁺ monocytes exhibited elevated inflammatory programs involving CXCL8 and TNF-NF-κB signaling, alongside CD163-associated scavenging and tissue homeostasis features. CD163 mediates the clearance of hemoglobin-haptoglobin complexes and participates in inflammatory regulation, suggesting that these concurrent programs may reflect sustained responses to tissue injury rather than a simple shift toward an anti-inflammatory phenotype. Nevertheless, their presence after seroreversion suggests that these programs alone are insufficient to explain serofast status, pointing to additional immune alterations that distinguish these outcomes.

Beyond these shared changes, serofast patients showed additional IFN-STAT1 activation, supported by increased pSTAT1 in CD14⁺ monocytes. This suggests that serofast status involves a distinct interferon-responsive state rather than simply greater activation of the inflammatory programs observed after seroreversion. Interferon signaling, which can enhance antimicrobial defenses, may also contribute to inflammation and tissue injury. These observations raise the possibility that serofast-associated immune activation does not necessarily translate into more effective pathogen clearance. Whether this state reflects residual microbial stimulation or longer-lasting immune responses following pathogen elimination remains unresolved.

NK cells exhibited a distinct pattern of post-treatment immune remodeling, characterized by reduced JUN expression and degranulation in serofast patients. Previous studies demonstrated that JNK signaling contributes to NK cell cytotoxicity through cytotoxic granule polarization^46^. Our findings extend this role by implicating JNK signaling in the magnitude and kinetics of cytotoxic, inflammatory, and tissue homeostasis-associated NK cell responses, raising the possibility that reduced JUN expression in serofast patients reflects broader alterations in NK cell functional competence. These observations suggest that reduced JUN-associated signaling may have broader consequences for NK cell functional competence in serofast patients. For example, reduced NK cell degranulation could compromise cytotoxic surveillance against virus-infected and transformed cells. Moreover, IFN-γ enhances macrophage CD64 expression and phagocytosis of opsonized *T. pallidum*^13^, suggesting that altered NK cell cytokine responses may also affect macrophage-mediated clearance of the pathogen. Whether reduced JUN expression directly contributes to NK cell dysfunction in serofast patients and whether restoring JUN expression could improve NK cell effector responses remain to be determined.

HIV coinfection adds a clinically relevant dimension to NK cell remodeling, as HIV-associated immunosuppression has been linked to impaired serological responses following syphilis treatment, whereas ART may mitigate this risk^47,48^. In this context, the partial recovery of NK cell degranulation alongside selective mitochondrial complex I remodeling in our ART-treated coinfected patients suggests that HIV does not uniformly exacerbate NK cell dysfunction. The similar complex I signature observed in HIV-only individuals further suggests an HIV-associated adaptation. Notably, increased degranulation was observed despite persistent serofast status. Given the importance of macrophage-mediated opsonophagocytosis in *T. pallidum* clearance^13^, enhanced NK cell cytotoxic responses may not necessarily translate into improved antitreponemal immunity. Whether HIV-associated NK cell remodeling influences subsequent serological outcomes remains to be determined.

Collectively, our findings indicate that nontreponemal antibody titers do not fully reflect innate immune recovery after syphilis treatment. Persistent monocyte and NK cell alterations, including monocyte IFN–STAT1 activation and HIV-associated changes in mitochondrial regulation, identify candidate immune pathways associated with post-treatment immune dysregulation. These findings provide a basis for evaluating cellular immune profiling as a complement to conventional serological monitoring and for exploring strategies to improve immune recovery after treatment.

### Limitations

This study has several limitations. The modest size of the clinical cohorts, particularly the HIV-coinfected group, limits statistical power and the assessment of potential confounders. The cross-sectional design also prevents us from determining whether the observed immune abnormalities contribute to persistent seroreactivity or arise from ongoing antigenic or inflammatory stimulation. Moreover, the absence of a prospective longitudinal cohort with immune profiling before treatment and serial follow-up precludes determining whether these immune features precede and predict subsequent serological outcomes, including seroreversion or serofast status. Peripheral blood transcriptional profiles do not directly establish tissue-level immune function, microbial persistence, or metabolic activity. Although pharmacological experiments support the involvement of JNK signaling and complex I in NK cell responses, they do not establish molecular specificity or directly demonstrate their roles in patient-associated immune alterations. Furthermore, the HIV-only comparison involved independently generated transcriptomic datasets, and the K562-based assays cannot determine antimicrobial protection in vivo. Prospective cohorts with longitudinal immune profiling and serological measurements from before treatment through follow-up will be needed to determine whether these immune alterations precede and predict the development of serofast status.

## RESOURCE AVAILABILITY

## Lead contact

Further information and requests for resources should be directed to the lead contact, Yetao Wang.

## Materials availability

This study did not generate new unique reagents.

## Data availability

The scRNA-seq data generated in this study have been deposited in the Genome Sequence Archive for Humans (GSA-Human) at the National Genomics Data Center under accession number HRA021795.

## Acknowledgments

This study was supported by National Natural Science Foundation of China (82471876), CAMS Innovation Fund for Medical Sciences (CIFMS) (2024-I2M-3-005) and Hospital for Skin Diseases, Institute of Dermatology, Chinese Academy of Medical Sciences and Peking Union Medical College grant (3301030103119) to Yetao Wang. The graphical abstract and schematic illustrations were created using BioRender and BioGDP.

## Author contributions

Yetao Wang: conceptualization; experiment design; data analysis; resources; supervision; validation; investigation; visualization; methodology; data curation; software; writing – original draft, review and editing; project administration; funding acquisition. Qin Wei, Yifan Jin, Yue Zhang: experiment design; performing experiments; data analysis; validation; visualization; methodology; data curation; writing – original draft, review and editing. Xinyuan Li, Qihang Zheng: data analysis. Lu Gan, Wenjing Le, Anlan Hong, Yuanyuan Zhao, Sai Li, Lixia Huang, Futing Liu, Qi Wang, Shaochun Chen, Xiaohong Su, Qianqiu Wang, Xiangsheng Chen, Suying Feng, Wujian Ke, Wenqi Xu, Ning Yan, Wei Chen, Zihai Li: resource. Xiaoyun Yu, Tong Lin: resource, review manuscript.

## FIGURE TITLES AND LEGENDS

**Figure S1. Study design and single-cell characterization of PBMCs**

(A) Schematic overview of the study design and experimental workflow. PBMCs from HC (n=10), SR (n=10), SF-L (n=10), and SF-H (n=6) were subjected to scRNA-seq, followed by marker validation and functional analyses. (B) UMAP of PBMCs colored by annotated cell types. (C) Dot plot showing representative marker expression across annotated cell types.

**Figure S2. Transcriptional profiles and marker validation in CD16^+^ monocytes across serological outcomes**

(A) Heatmap of standardized pathway scores in the HC, SR, SF-L, and SF-H groups.

(B) Heatmap of relative expression levels of selected genes, grouped into innate signaling, inflammatory response, adhesion and remodeling, antigen processing, and stress-adaptive programs. (C-E) MFI of CXCL8 (C), CD163 (D) and pSTAT1 (E) in CD16⁺ monocytes (n=15 per group). For (C, and E), two-tailed Mann-Whitney U test. For (D), two-tailed unpaired t-tests. Each dot represents an individual donor. Data are mean ± SEM. ns, not significant.

**Figure S3. Functional programs and receptor expression in CD56^hi^NK cells across serological outcomes**

(A) Heatmap of Z-scored enrichment scores for the indicated pathways in CD56^hi^NK cells across the four clinical groups. (B) Heatmaps of Z-scored expression of activating and inhibitory receptors in CD56^hi^NK cells across the four clinical groups.

**Figure S4. Differential expression patterns and IFN-γ expression in NK cells across syphilis and HIV coinfection**

(A) Differential expression patterns in CD56^hi^ and CD56^dim^NK cells across three pairwise comparisons (Syph vs. HC, Syph+HIV vs. Syph, and Syph+HIV vs. HC). Colors indicate the direction and significance of differential expression, and bar plots show gene numbers for each pattern. (B) Venn diagrams showing overlapping and comparison-specific differentially expressed genes across six modules in both NK cell subsets. (C) Percentages of IFN-γ⁺ cells in total NK cells across groups (HC, n=10; Syph, n=9; Syph+HIV, n=7). Each dot represents an individual donor, two-tailed unpaired t-test, data are mean ± SEM. Comparisons without annotated *P* values were considered nonsignificant.

## METHODS

### Clinical samples

PBMCs from HC, SR, SF-L, SF-H, and Syph+HIV groups were collected at The Second Hospital of Nanjing. PBMCs from additional healthy donors used for *in vitro* functional experiments were obtained from the Jiangsu Province Blood Center. All participants provided written informed consent. The study was approved by the Ethics Committees of The Second Hospital of Nanjing (2025-LS-ky-018) and The Institute of Dermatology, Chinese Academy of Medical Sciences and Peking Union Medical College (2024-KY-042). Clinical characteristics of the participants are summarized in Table S1.

### PBMCs isolation

PBMCs were isolated from human peripheral blood by density gradient centrifugation using Lymphoprep. Briefly, blood samples were diluted with PBS and carefully layered onto Lymphoprep, followed by centrifugation at 500 × g for 30 min at room temperature. The PBMC layer was collected and washed three times with PBS. Isolated PBMCs were either used immediately for subsequent experiments or cryopreserved in FBS supplemented with 10% DMSO.

### scRNA-Seq library preparation

PBMC-derived single-cell suspensions were adjusted to a concentration of 700-1,200 cells/μL. Single-cell 5′ gene expression libraries were constructed using the MobiCube High-Throughput Single-Cell 5′ Transcriptome Kit V2.1 (MobiDrop, Cat. No. S060200501) according to the manufacturer’s instructions.

### scRNA-Seq data processing and quality control

Raw sequencing data were processed using MobiVision. The syphilis cohort (HC, SR, SF-L, and SF-H) comprised 343,997 cells from 36 PBMC samples, with a mean sequencing depth of 36384 reads per cell. Quality control was performed using Seurat v5. Cells with more than 300 and fewer than 6,500 detected genes, fewer than 50,000 total UMIs, and mitochondrial and hemoglobin UMI percentages below 20% and 1%, respectively, were retained, yielding 339,549 cells. For the HIV coinfection analysis, 255,561 cells from 26 donors (HC, Syph (SF-L), and Syph+HIV) were independently processed using the same criteria, retaining 220,098 cells. Both datasets underwent normalization, variable feature selection, scaling, principal component analysis, and Harmony integration using sample identity as the batch variable.

### Cell clustering and annotation

Cluster-enriched genes were identified using the FindAllMarkers function in Seurat with the Wilcoxon rank-sum test. Candidate markers were restricted to positively enriched genes detected in at least 25% of cells in the target cluster, with a log_2_FC≥0.5 and an adjusted *P*<0.05. Based on established marker expression, cells were annotated into 10 major populations: T cells (CD3D, CD3G, and CD3E), NK cells (NKG7, FCGR3A, and NCAM1), cycling cells (MKI67, PCNA, and CKS1B), B cells (MS4A1, CD79A, and CD79B), plasmablasts (IGHA1, IGHA2, and IGHG1), monocytes (CD14, CSF3R, and CD68), plasmacytoid dendritic cells (pDCs; CLEC4C, IL3RA, and LILRA4), type 2 conventional dendritic cells (cDC2s; FCER1A, CLEC10A, and CD1C), platelets (ITGA2B, ITGB3, and GP9), and hematopoietic stem and progenitor cells (HSPCs; CD34, KIT, and HOXA9). NK cells and myeloid cells were subsequently extracted and independently re-clustered using a similar workflow, with subcluster markers identified using the same approach and thresholds. NK cells were classified into CD56^hi^NK cells (NCAM1, IL7R, GZMK, TCF7, SELL, BACH2, KLRC1, and XCL1) and CD56^dim^NK cells (FCGR3A, FGFBP2, GZMH, GZMB, PRDM1, CCL3, CCL4, and CX3CR1) based on their relative marker expression. The monocyte and dendritic-cell compartment was further resolved into CD14+ monocytes (CD14, S100A8, S100A9, and S100A12), CD16+ monocytes (FCGR3A, LILRB1, IFITM3, and CX3CR1), cDC1s (CLEC9A, CADM1, IDO1, and XCR1), cDC2s (CLEC10A, CD1C, and FCER1A), and pDCs (CLEC4C, IRF7, IL3RA, and TCF4).

### Quantification of differentially expressed genes across clinical groups

Pairwise differential expression analyses were performed across the four clinical groups within each myeloid and NK cell subset using the Wilcoxon rank-sum test implemented in Seurat’s FindMarkers function. DEGs were identified using the following criteria: |log₂FC|>0.5 for myeloid cell subsets, |log₂FC|>0.25 for NK cell subsets, pct.1>0.2, and adjusted *P*<0.05.

### Functional gene set analysis

Gene set variation analysis (GSVA; version 2.0.4) was performed using donor-level mean log-normalized expression profiles. Gene sets included selected Hallmark and Reactome gene sets from the Molecular Signatures Database (MSigDB) which were retrieved using msigdbr, and custom-defined functional signatures. Sources of database-derived gene sets and gene members of custom-defined signatures are provided in Table S2. GSVA scores were calculated using a Gaussian kernel, averaged across donors within each study group, and standardized to Z-scores across groups for heatmap visualization.

### Flow cytometry

PBMCs were initially stained with Fixable Viability Dye eFluor 780 (Invitrogen, Cat. No. 65-0865-18) to exclude dead cells. For surface marker staining, cells were incubated with fluorochrome-conjugated antibodies in MACS buffer at 4°C for 30 min in the dark. For intracellular staining, cells were subsequently fixed and permeabilized using the Foxp3/Transcription Factor Staining Buffer Set (eBioscience, Cat. No. 00-5523-00), followed by incubation with antibodies against intracellular targets in permeabilization buffer at 4°C for 30 min in the dark. After washing, cells were resuspended in MACS buffer for flow cytometric analysis. Samples with insufficient cell numbers for reliable analysis were excluded.

### Transcription factor activity inference

Transcription factor (TF) activities were inferred using the univariate linear model (ULM) method in decoupleR (version 2.12.0) with DoRothEA interactions of confidence levels A-C, requiring ≥5 matched target genes per TF. Analyses were performed on donor-level mean normalized expression profiles for each cell population. Activity scores were averaged within each study group and standardized to Z-scores across groups for heatmap visualization.

### Identification of group enriched marker genes and functional modules

Within each cell subset, group-enriched marker genes were identified by comparing each clinical group with all other groups combined using Seurat’s FindMarkers function (Wilcoxon rank-sum test), with thresholds of log_2_FC≥0.25, adjusted *P*<0.05, and expression in ≥20% of cells in the target group. Mean log-normalized expression of the identified genes was calculated for each clinical group. Genes were then clustered based on their expression patterns across groups, and the resulting modules were functionally annotated according to the known biological roles of their constituent genes.

### Functional gene signature scoring

An AP-1/NF-κB response signature was defined using AP-1-associated immediate early genes (JUN, JUNB, JUND, FOS, FOSB, FOSL2, ATF3, EGR1, NR4A1, DUSP1, DUSP2, ZFP36, IER2, and BTG2) and NF-κB response genes (NFKBIA, NFKBIE, NFKBIZ, TNFAIP2, TNFAIP3, BIRC3, TRAF1, TNIP1, TNIP2, ICAM1, CFLAR, ZC3H12A, IER3, TNF, CCL3, CCL4, CCL4L2, and CD69). Signature scores were calculated for individual cells using UCell (version 2.10.1) on the RNA count matrix. For statistical comparisons, pairwise differences between study groups were assessed using two-sided Wilcoxon rank-sum tests, with *P* values adjusted using the Benjamini-Hochberg method. Adjusted *P*<0.05 was considered statistically significant.

### Rule-based classification of six infection-state expression modules

Differential expression analysis was performed separately in CD56^hi^ and CD56^dim^ NK cells for Syph versus HC, Syph+HIV versus HC, and Syph+HIV versus Syph using Seurat’s FindMarkers function with the Wilcoxon rank-sum test. DEGs were defined by a Bonferroni-adjusted *P*<0.05, |average log₂FC|≥0.25, and detection in ≥25% of cells in the higher-expression group. Genes were classified by their direction and significance across comparisons into 18 predefined patterns and grouped into six modules (Figure S4A and S4B). For visualization, log-normalized expression was averaged within donors and then across donors in each clinical group. Group means were Z-scored for each gene within each NK-cell subset and displayed as line plots and heatmaps. Black lines indicate module-average trajectories (the mean standardized expression of genes within each cluster). Gene trajectories were colored by correlation-based scores, calculated as (r+1)/2, where r is the Pearson correlation with the module-average trajectory (Figure 6).

### Functional enrichment and redundancy reduction of expression modules

Functional enrichment was performed separately for each expression module within each NK cell subset using GO biological process, cellular component, and molecular function terms, KEGG pathways, Hallmark gene sets, and Reactome pathways. Analyses used clusterProfiler and ReactomePA, with Hallmark gene sets obtained through msigdbr. For each NK cell subset, the enrichment background comprised the union of genes tested in the three differential expression comparisons (Syph versus HC, Syph+HIV versus HC, and Syph+HIV versus Syph), including both significant and nonsignificant genes. The effective background was restricted to genes annotated in the corresponding database.

Terms with a Benjamini-Hochberg adjusted *P*<0.05 and overlapping module genes ≥2 were considered significant. Significant results were combined across databases within each subset and module, then ranked by increasing adjusted *P* value, decreasing overlap gene count, and decreasing fold enrichment (Table S3 and S4).

Redundancy was reduced using a custom greedy filtering procedure. For each module, similarity between enriched terms was assessed using the Jaccard index of their overlapping module genes^49^. Jaccard similarity was calculated as the intersection divided by the union of module genes overlapping each pair of terms. Rank-based greedy filtering removed terms with similarity ≥0.70 to any previously retained term or identical normalized descriptions. All remaining terms were retained without a fixed limit per module but displayed according to their biological functions.

### Degranulation assay

For in vitro validation using samples from patients with syphilis, 1 × 10^5^ PBMCs and 1 × 10^5^ K562 cells were added per well. For all other in vitro validation experiments, 2 × 10^5^ PBMCs and 2 × 10^5^ K562 cells were added per well. In both cases, PBMCs and K562 cells were combined at a 1:1 ratio in V-bottom 96-well plates in a total volume of 200 μL per well containing Protein Transport Inhibitor Cocktail (500×) (eBioscience, Cat. No. 00-4980-03). The cells were mixed, centrifuged at 50 × g for 1 min, and incubated at 37°C in 5% CO₂ for 4 h. Surface CD107a expression on NK cells was then assessed by flow cytometry.

### Reagents and resources

The reagents and resources used in this study are listed in Table S5.

### QUANTIFICATION AND STATISTICAL ANALYSIS

All statistical analyses were performed using GraphPad Prism 10. Data are presented as mean ± SEM. Statistical tests included two-tailed Mann-Whitney U tests, paired and unpaired t-tests, Pearson correlation tests, and two-way ANOVA, as indicated in corresponding figure legends. A *P* value <0.05 was considered statistically significant. Comparisons without annotated *P* values were considered nonsignificant.

