## Supplemental Figures for "Distinct innate immune states associate with divergent serological outcomes after syphilis treatment"

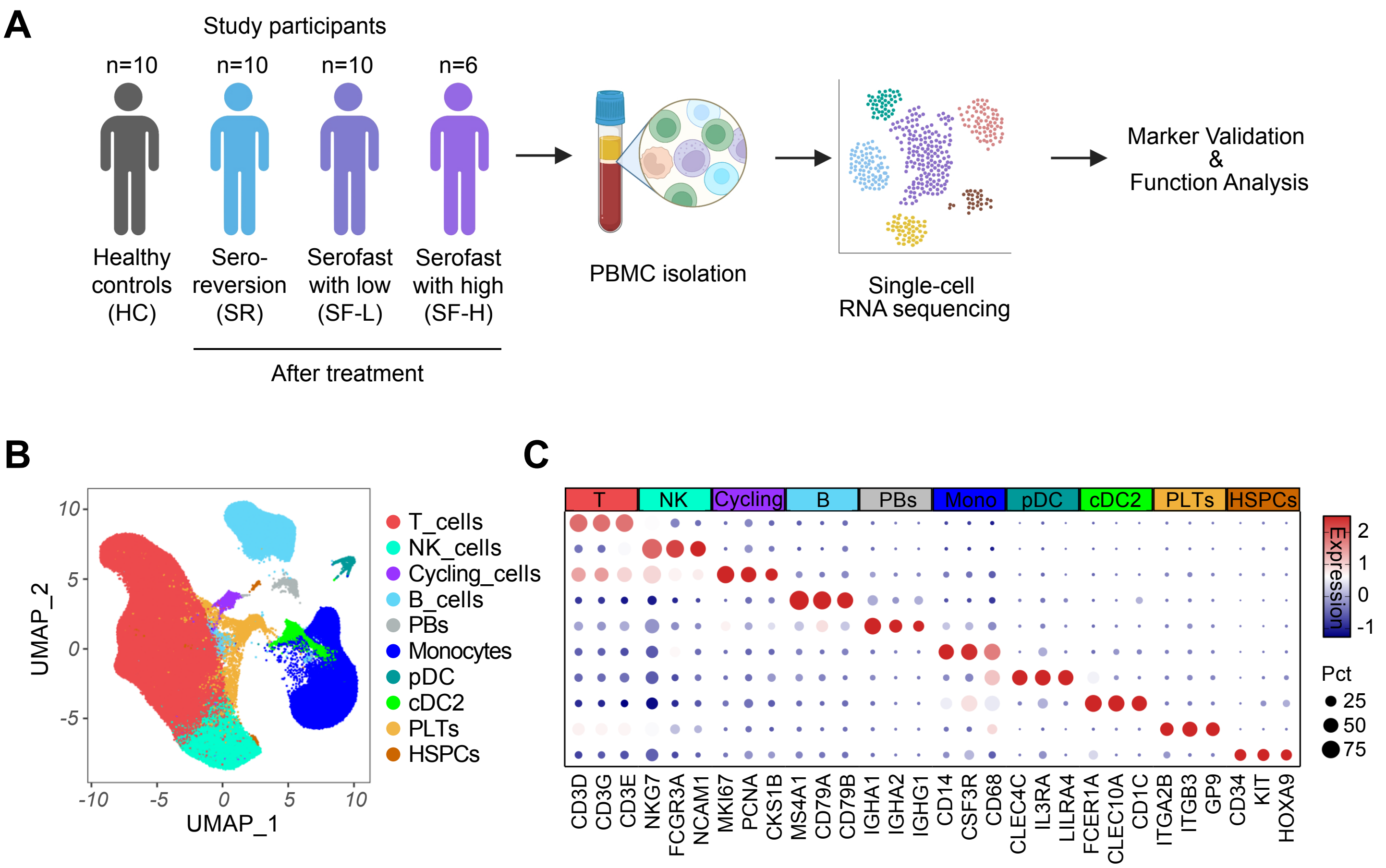

A
Study participants
n=10
n=10
n=10
n=6
Healthy controls (HC)
Sero-reversion (SR)
Serofast with low (SF-L)
Serofast with high (SF-H)
After treatment
PBMC isolation
Single-cell RNA sequencing
Marker Validation & Function Analysis
B
UMAP_2
UMAP_1
T_cells
NK_cells
Cycling_cells
B_cells
PBs
Monocytes
pDC
cDC2
PLTs
HSPCs
C
T
NK
Cycling
B
PBs
Mono
pDC
cDC2
PLTs
HSPCs
Expression
Pct
25
50
75
CD3D
CD3G
CD3E
NKG7
FCGR3A
NCAM1
MKI67
PCNA
CKS1B
MS4A1
CD79A
CD79B
IGHA1
IGHA2
IGHG1
CD14
CSF3R
CD68
CLEC4C
IL3RA
LILRA4
FCER1A
CLEC10A
CD1C
ITGA2B
ITGB3
GP9
CD34
KIT
HOXA9


**Figure S1**

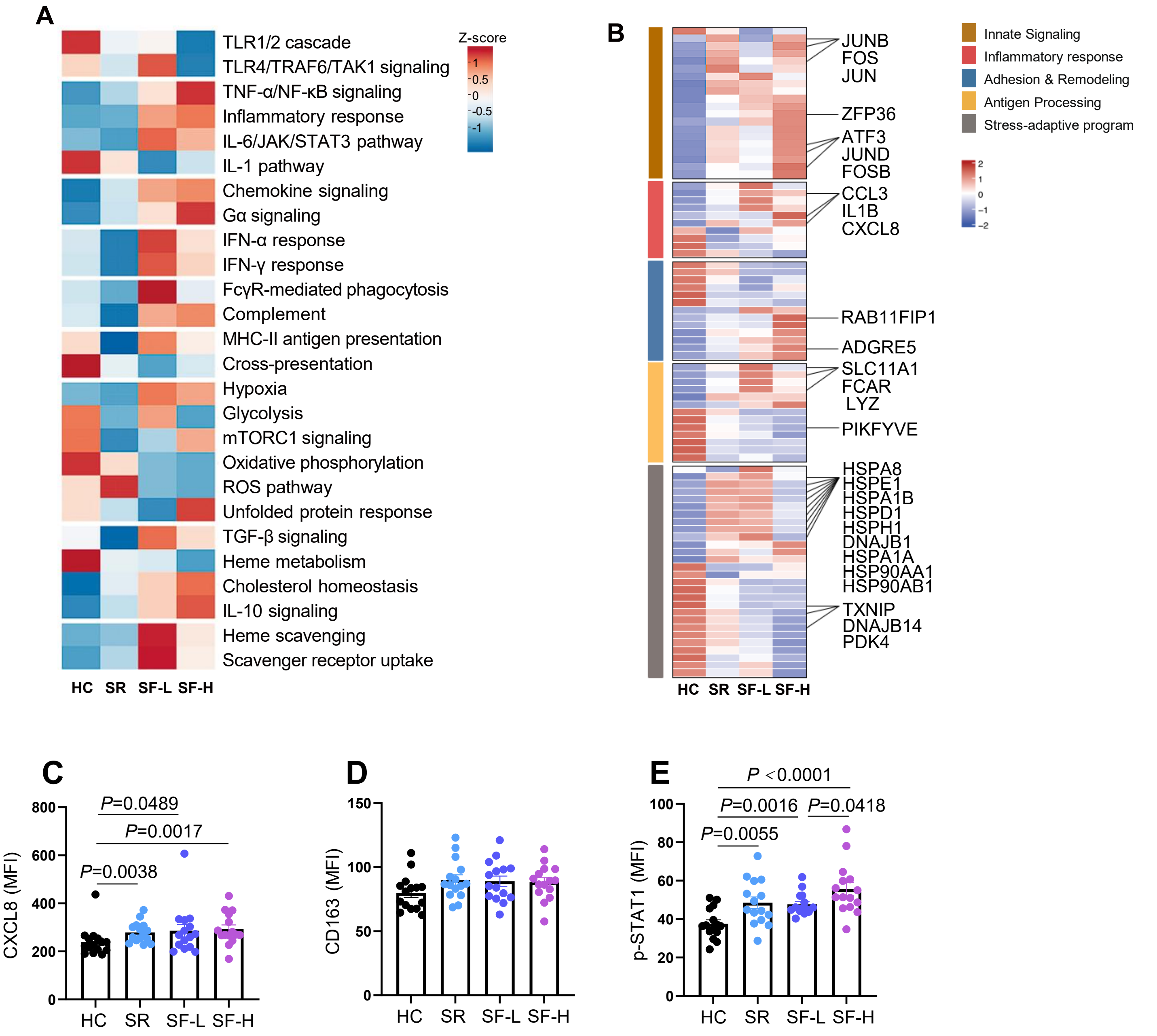
A
TLR1/2 cascade
TLR4/TRAF6/TAK1 signaling
TNF-α/NF-κB signaling
Inflammatory response
IL-6/JAK/STAT3 pathway
IL-1 pathway
Chemokine signaling
Gα signaling
IFN-α response
IFN-γ response
FcγR-mediated phagocytosis
Complement
MHC-II antigen presentation
Cross-presentation
Hypoxia
Glycolysis
mTORC1 signaling
Oxidative phosphorylation
ROS pathway
Unfolded protein response
TGF-β signaling
Heme metabolism
Cholesterol homeostasis
IL-10 signaling
Heme scavenging
Scavenger receptor uptake
HC SR SF-L SF-H
Z-score
1
0.5
0
-0.5
-1
B
JUNB
FOS
JUN
ZFP36
ATF3
JUND
FOSB
CCL3
IL1B
CXCL8
RAB11FIP1
ADGRE5
SLC11A1
FCAR
LYZ
PIKFYVE
HSPA8
HSPE1
HSPA1B
HSPD1
HSPH1
DNAJB1
HSPA1A
HSP90AA1
HSP90AB1
TXNIP
DNAJB14
PDK4
HC SR SF-L SF-H
Innate Signaling
Inflammatory response
Adhesion & Remodeling
Antigen Processing
Stress-adaptive program
2
1
0
−1
−2
C
CXCL8 (MFI)
P=0.0489
P=0.0017
P=0.0038
800
600
400
200
0
HC SR SF-L SF-H
D
CD163 (MFI)
150
100
50
0
HC SR SF-L SF-H
E
p-STAT1 (MFI)
P <0.0001
P=0.0016
P=0.0418
P=0.0055
100
80
60
40
20
0
HC SR SF-L SF-H

**Figure S2**

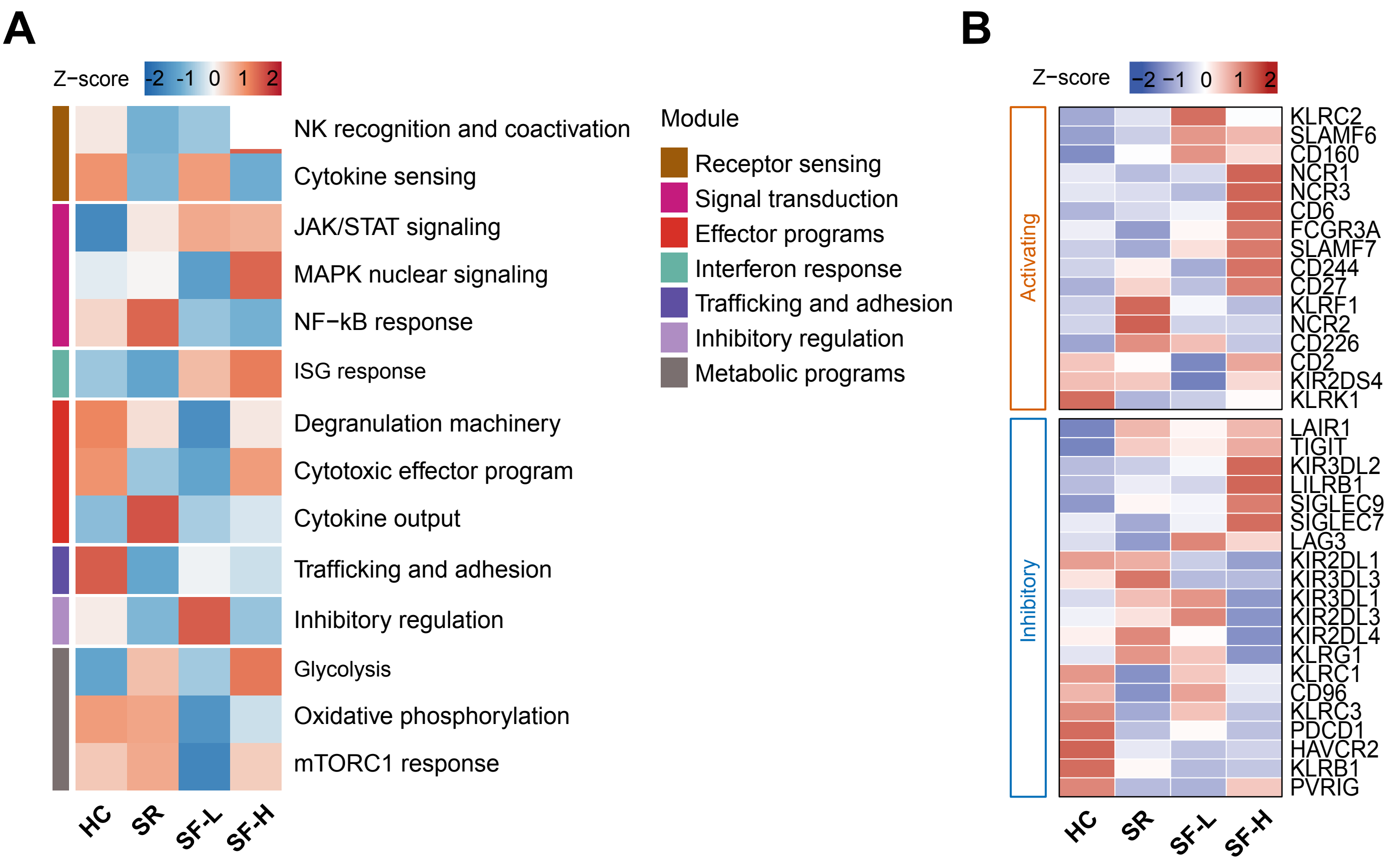
A
Z−score
-2 -1 0 1 2
NK recognition and coactivation
Cytokine sensing
JAK/STAT signaling
MAPK nuclear signaling
NF−kB response
ISG response
Degranulation machinery
Cytotoxic effector program
Cytokine output
Trafficking and adhesion
Inhibitory regulation
Glycolysis
Oxidative phosphorylation
mTORC1 response
Module
Receptor sensing
Signal transduction
Effector programs
Interferon response
Trafficking and adhesion
Inhibitory regulation
Metabolic programs
HC
SR
SF-L
SF-H
B
Z−score
−2 −1 0 1 2
Activating
Inhibitory
KLRC2
SLAMF6
CD160
NCR1
NCR3
CD6
FCGR3A
SLAMF7
CD244
CD27
KLRF1
NCR2
CD226
CD2
KIR2DS4
KLRK1
LAIR1
TIGIT
KIR3DL2
LILRB1
SIGLEC9
SIGLEC7
LAG3
KIR2DL1
KIR3DL3
KIR3DL1
KIR2DL3
KIR2DL4
KLRG1
KLRC1
CD96
KLRC3
PDCD1
HAVCR2
KLRB1
PVRIG
HC
SR
SF-L
SF-H

**Figure S3**

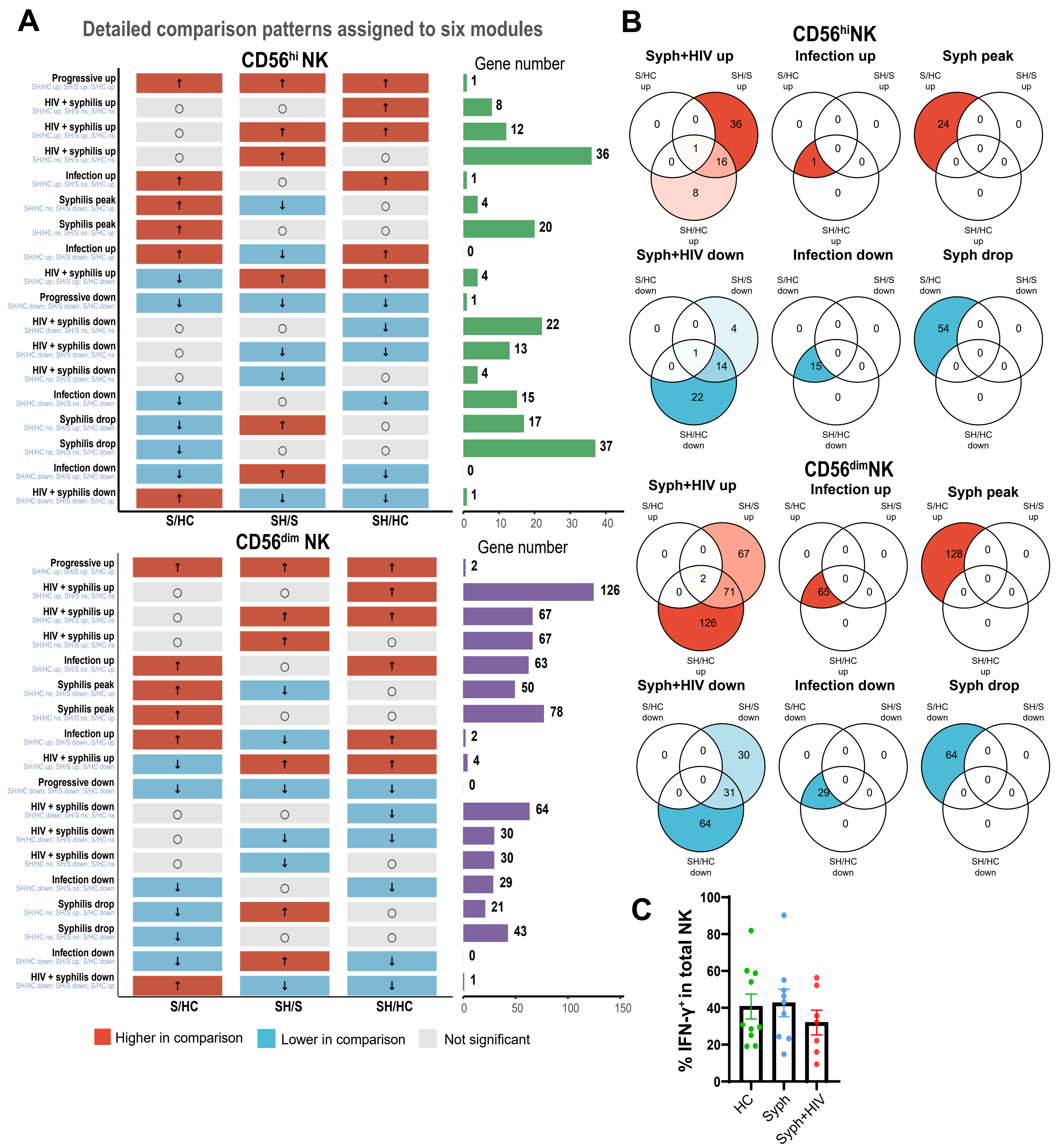
A
Detailed comparison patterns assigned to six modules
CD56hi NK
Gene number
Progressive up
HIV + syphilis up
HIV + syphilis up
HIV + syphilis up
Infection up
Syphilis peak
Syphilis peak
Infection up
HIV + syphilis up
Progressive down
HIV + syphilis down
HIV + syphilis down
HIV + syphilis down
Infection down
Syphilis drop
Syphilis drop
Infection down
HIV + syphilis down
1
8
12
36
1
4
20
0
4
1
22
13
4
15
17
37
0
1
S/HC
SH/S
SH/HC
0
10
20
30
40
CD56dim NK
Gene number
Progressive up
HIV + syphilis up
HIV + syphilis up
HIV + syphilis up
Infection up
Syphilis peak
Syphilis peak
Infection up
HIV + syphilis up
Progressive down
HIV + syphilis down
HIV + syphilis down
HIV + syphilis down
Infection down
Syphilis drop
Syphilis drop
Infection down
HIV + syphilis down
2
126
67
67
63
50
78
2
4
0
64
30
30
29
21
43
0
1
S/HC
SH/S
SH/HC
0
50
100
150
Higher in comparison
Lower in comparison
Not significant
B
CD56hiNK
Syph+HIV up
Infection up
Syph peak
Syph+HIV down
Infection down
Syph drop
CD56dimNK
Syph+HIV up
Infection up
Syph peak
Syph+HIV down
Infection down
Syph drop
S/HC up
SH/S up
SH/HC up
S/HC down
SH/S down
SH/HC down
C
% IFN-γ+ in total NK
0
20
40
60
80
100
HC
Syph
Syph+HIV

Figure S4
