## Supplemental Tables for "Distinct innate immune states associate with divergent serological outcomes after syphilis treatment"

| Table S1 │ Characteristics of PBMC donors included in scRNA-seq and flow cytometry analyses. | | | | | | | |
|---|---|---|---|---|---|---|---|
| Sample ID# | Group | Gender | Age | Technology | TRUST titer at sampling | Time since syphilis diagnosis (years) | Related Figure |
| HC_S11 | HC | female | 40 | single cell sequencing , flow cytometry | NA | NA | Figure 1, 2, 3, 4, 5, 6, 7 |
| HC_S38 | HC | female | 55 | single cell sequencing , flow cytometry | NA | NA | Figure 1, 2, 3, 4, 6, 7 |
| HC_S62 | HC | male | 31 | single cell sequencing , flow cytometry | NA | NA | Figure 1, 2, 3, 4, 6, 7 |
| HC_S68 | HC | male | 45 | single cell sequencing , flow cytometry | NA | NA | Figure 1, 2, 3, 4, 6, 7 |
| HC_S73 | HC | male | 31 | single cell sequencing , flow cytometry | NA | NA | Figure 1, 2, 3, 4, 6, 7 |
| HC_S80 | HC | male | 23 | flow cytometry | NA | NA | Figure 4, 5 |
| HC_S81 | HC | male | 23 | flow cytometry | NA | NA | Figure 2, 4, 5, 7 |
| HC_S83 | HC | male | 19 | flow cytometry | NA | NA | Figure 2, 4, 5, 7 |
| HC_S96 | HC | female | 48 | single cell sequencing , flow cytometry | NA | NA | Figure 1, 2, 3, 4, 6, 7 |
| HC_S118 | HC | male | 35 | single cell sequencing , flow cytometry | NA | NA | Figure 1, 2, 3, 4, 6, 7 |
| HC_S126 | HC | male | 36 | flow cytometry | NA | NA | Figure 2, 4, 5, 7 |
| HC_S132 | HC | male | 30 | single cell sequencing , flow cytometry | NA | NA | Figure 1, 2, 3, 4, 6, 7 |
| HC_S134 | HC | male | 32 | single cell sequencing , flow cytometry | NA | NA | Figure 1, 2, 3, 4, 6, 7 |
| HC_S137 | HC | female | 28 | flow cytometry | NA | NA | Figure 2, 4, 5, 7 |
| HC_S143 | HC | male | 24 | flow cytometry | NA | NA | Figure 2, 4, 5, 7 |
| HC_S162 | HC | male | 43 | flow cytometry | NA | NA | Figure 4, 5 |
| HC_S163 | HC | female | 49 | flow cytometry | NA | NA | Figure 4, 5 |
| HC_S170 | HC | male | 45 | flow cytometry | NA | NA | Figure 4, 5 |
| HC_S239 | HC | female | 52 | single cell sequencing , flow cytometry | NA | NA | Figure 1, 2, 3, 4, 6, 7 |
| SR_S9 | SR | male | 29 | flow cytometry | Nonreactive | > 2 | Figure 2, 4, 5, 7 |
| SR_S29 | SR | female | 41 | flow cytometry | Nonreactive | 10 | Figure 2, 4, 5, 7 |
| SR_S30 | SR | female | 56 | single cell sequencing , flow cytometry | Nonreactive | 1 | Figure 1, 2, 3, 4, 6, 7 |
| SR_S33 | SR | male | 41 | single cell sequencing , flow cytometry | Nonreactive | 1 | Figure 1, 3, 4, 5, 6, 7 |
| SR_S36 | SR | female | 24 | single cell sequencing , flow cytometry | Nonreactive | 2 | Figure 1, 3, 4, 5, 6, 7 |
| SR_S43 | SR | female | 20 | single cell sequencing , flow cytometry | Nonreactive | 0.5 | Figure 1, 3, 4, 5, 6, 7 |
| SR_S61 | SR | male | 26 | single cell sequencing , flow cytometry | Nonreactive | 2 | Figure 1, 2, 3, 4, 6, 7 |
| SR_S71 | SR | female | 47 | single cell sequencing , flow cytometry | Nonreactive | 2 | Figure 1, 2, 3, 4, 6, 7 |
| SR_S86 | SR | female | 65 | flow cytometry | Nonreactive | 0.2 | Figure 2, 4, 5, 7 |
| SR_S94 | SR | female | 55 | single cell sequencing , flow cytometry | Nonreactive | 0.25 | Figure 1, 2, 3, 4, 6, 7 |
| SR_S107 | SR | male | 63 | flow cytometry | Nonreactive | 11 | Figure 2, 4, 5, 7 |
| SR_S161 | SR | female | 24 | single cell sequencing , flow cytometry | Nonreactive | 3 | Figure 1, 2, 3, 4, 6, 7 |
| SR_S171 | SR | male | 37 | flow cytometry | Nonreactive | 10 | Figure 2, 4, 5, 7 |
| SR_S201 | SR | female | 58 | single cell sequencing , flow cytometry | Nonreactive | 1 | Figure 1, 2, 3, 4, 6, 7 |
| SR_S202 | SR | female | 63 | flow cytometry | Nonreactive | 10 | Figure 2, 4, 5, 7 |
| SR_S206 | SR | male | 62 | flow cytometry | Nonreactive | 10 | Figure 2, 4, 5, 7 |
| SR_S210 | SR | male | 41 | flow cytometry | Nonreactive | > 2 | Figure 2, 4, 5, 7 |
| SR_S223 | SR | male | 61 | flow cytometry | Nonreactive | 0.5 | Figure 2, 4, 5, 7 |
| SR_S262 | SR | male | 43 | flow cytometry | Nonreactive | 1 | Figure 4, 5 |
| SR_S265 | SR | male | 30 | flow cytometry | Nonreactive | > 3 | Figure 4, 5 |
| SR_S267 | SR | female | 42 | single cell sequencing | Nonreactive | > 2 | Figure 1, 3, 6, 7 |
| SF-L_S7 | SF-L | female | 56 | single cell sequencing , flow cytometry | 1： 4 | 5 | Figure 1, 2, 3, 4, 6, 7 |

| | | | | | | | |
|---|---|---|---|---|---|---|---|
| SF-L_S37 | SF-L | female | 25 | single cell sequencing , flow cytometry | 1: 2 | 3 | Figure 1, 2, 3, 4, 6, 7 |
| SF-L_S48 | SF-L | male | 41 | flow cytometry | 1: 2 | 2 | Figure 4, 5 |
| SF-L_S56 | SF-L | male | 32 | flow cytometry | 1: 2 | 2 | Figure 2, 4, 5, 7 |
| SF-L_S57 | SF-L | female | 35 | flow cytometry | 1: 2 | 7 | Figure 2, 4, 5, 7 |
| SF-L_S76 | SF-L | female | 23 | flow cytometry | 1: 2 | 2 | Figure 4, 5 |
| SF-L_S89 | SF-L | female | 55 | flow cytometry | 1: 2 | 3 | Figure 2, 4, 5, 7 |
| SF-L_S90 | SF-L | female | 31 | flow cytometry | 1: 2 | > 2 | Figure 2, 4, 5, 7 |
| SF-L_S104 | SF-L | female | 29 | flow cytometry | 1: 4 | 2 | Figure 2, 4, 5, 7 |
| SF-L_S108 | SF-L | female | 37 | single cell sequencing , flow cytometry | 1: 2 | 6 | Figure 1, 2, 3, 4, 6, 7 |
| SF-L_S109 | SF-L | male | 29 | single cell sequencing , flow cytometry | 1: 4 | 2 | Figure 1, 2, 3, 4, 6, 7 |
| SF-L_S115 | SF-L | male | 76 | flow cytometry | 1: 2 | 10 | Figure 2, 4, 5, 7 |
| SF-L_S125 | SF-L | male | 31 | flow cytometry | 1: 2 | 2 | Figure 2, 4, 5, 7 |
| SF-L_S131 | SF-L | male | 75 | flow cytometry | 1: 4 | 2 | Figure 4, 5 |
| SF-L_S140 | SF-L | female | 22 | flow cytometry | 1: 2 | 3 | Figure 4, 5 |
| SF-L_S141 | SF-L | male | 43 | flow cytometry | 1: 2 | 5 | Figure 4, 5 |
| SF-L_S193 | SF-L | female | 22 | single cell sequencing , flow cytometry | 1: 4 | 2 | Figure 1, 2, 3, 4, 6, 7 |
| SF-L_S229 | SF-L | female | 33 | single cell sequencing , flow cytometry | 1: 2 | 3 | Figure 1, 2, 3, 4, 6, 7 |
| SF-L_S248 | SF-L | male | 28 | single cell sequencing , flow cytometry | 1: 2 | 2 | Figure 1, 2, 3, 4, 6, 7 |
| SF-L_S278 | SF-L | female | 23 | single cell sequencing , flow cytometry | 1: 2 | 3 | Figure 1, 2, 3, 4, 6, 7 |
| SF-L_S295 | SF-L | female | 19 | single cell sequencing | 1: 4 | 2 | Figure 1, 3, 6, 7 |
| SF-L_S302 | SF-L | male | 30 | single cell sequencing | 1: 4 | 3 | Figure 1, 3, 6, 7 |
| SF-H_S2 | SF-H | female | 65 | single cell sequencing , flow cytometry | 1: 8 | 2 | Figure 1, 2, 3, 4, 5, 6, 7 |
| SF-H_S28 | SF-H | male | 21 | flow cytometry | 1: 8 | 1 | Figure 2, 4, 7 |
| SF-H_S44 | SF-H | male | 32 | flow cytometry | 1: 16 | 6 | Figure 2, 4, 5, 7 |
| SF-H_S55 | SF-H | female | 29 | single cell sequencing , flow cytometry | 1: 32 | 2 | Figure 1, 3, 4, 5, 6, 7 |
| SF-H_S84 | SF-H | male | 49 | single cell sequencing , flow cytometry | 1: 8 | > 2 | Figure 1, 2, 3, 4, 6, 7 |
| SF-H_S103 | SF-H | male | 58 | single cell sequencing , flow cytometry | 1: 8 | 2 | Figure 1, 3, 6, 7 |
| SF-H_S151 | SF-H | female | 33 | single cell sequencing , flow cytometry | 1: 16 | > 2 | Figure 1, 3, 4, 5, 6, 7 |
| SF-H_S174 | SF-H | male | 59 | single cell sequencing , flow cytometry | 1: 8 | 2 | Figure 1, 3, 4, 5, 6, 7 |
| SF-H_S179 | SF-H | female | 32 | flow cytometry | 1: 64 | 1 | Figure 2, 4, 7 |
| SF-H_S185 | SF-H | male | 69 | flow cytometry | 1: 8 | 1 | Figure 2, 4, 7 |
| SF-H_S188 | SF-H | female | 20 | flow cytometry | 1: 32 | 1 | Figure 2, 4, 7 |
| SF-H_S244 | SF-H | female | 32 | flow cytometry | 1: 8 | 1 | Figure 2, 4, 7 |
| SF-H_S246 | SF-H | female | 19 | flow cytometry | 1: 32 | 1 | Figure 2, 4, 7 |
| SF-H_S282 | SF-H | male | 27 | flow cytometry | 1: 8 | 2 | Figure 2, 4, 5, 7 |
| SF-H_S283 | SF-H | female | 62 | flow cytometry | 1: 8 | > 2 | Figure 2, 4, 7 |
| SF-H_S293 | SF-H | male | 35 | flow cytometry | 1: 16 | 10 | Figure 2, 4, 5, 7 |
| SF-H_S298 | SF-H | female | 35 | flow cytometry | 1: 16 | 2 | Figure 2, 4, 5, 7 |
| SF-H_S299 | SF-H | male | 47 | flow cytometry | 1: 8 | 2 | Figure 2, 4, 5, 7 |
| SF-H_S355 | SF-H | male | 37 | flow cytometry | 1: 64 | 1 | Figure 2, 4, 7 |
| SF-H_S372 | SF-H | male | 39 | flow cytometry | 1: 8 | 2 | Figure 4, 5 |
| SF-H_S376 | SF-H | male | 41 | flow cytometry | 1: 8 | 5 | Figure 4, 5 |
| Syph-HIV_S13 | Syph+HIV | male | 35 | flow cytometry | 1: 1 | 2 | Figure 7 |
| Syph-HIV_S46 | Syph+HIV | male | 61 | single cell sequencing , flow cytometry | 1: 4 | > 2 | Figure 6, 7 |
| Syph-HIV_S52 | Syph+HIV | male | 42 | single cell sequencing , flow cytometry | 1: 1 | > 2 | Figure 6, 7 |
| Syph-HIV_S102 | Syph+HIV | male | 29 | flow cytometry | 1: 4 | 6 | Figure 7 |
| Syph-HIV_S123 | Syph+HIV | male | 50 | flow cytometry | 1: 1 | 10 | Figure 7 |

| | | | | | | | |
|---|---|---|---|---|---|---|---|
| Syph-HIV_S142 | Syph+HIV | male | 34 | single cell sequencing , flow cytometry | 1： 1 | 5 | Figure 6, 7 |
| Syph-HIV_S150 | Syph+HIV | male | 38 | flow cytometry | 1： 1 | 6 | Figure 7 |
| HIV_S165 | HIV | male | 30 | flow cytometry | 1： 2 | > 2 | Figure 6, 7 |
| Syph-HIV_S189 | Syph+HIV | male | 44 | single cell sequencing , flow cytometry | 1： 1 | > 2 | Figure 6, 7 |
| Syph-HIV_S215 | Syph+HIV | male | 60 | flow cytometry | 1： 2 | Unknown | Figure 7 |
| Syph-HIV_S240 | Syph+HIV | male | 40 | single cell sequencing | 1： 2 | 3 | Figure 6, 7 |

| Table S2 ∣ Gene-set sources or constituent genes used in pathway enrichment analysis | | |
|---|---|---|
| **Cell type** | **Pathway name** | **Gene-set source or constituent genes** |
| Monocytes | TLR1/2 cascade | MSigDB Reactome: REACTOME_TOLL_LIKE_RECEPTOR_TLR1_TLR2_CASCADE |
| Monocytes | TLR4/TRAF6/TAK1 signaling | MSigDB Reactome: REACTOME_TRAF6_MEDIATED_INDUCTION_OF_TAK1_COMPLEX_WITHIN_TLR4_COMPLEX |
| Monocytes | TNF-α/NF-κB signaling | MSigDB Hallmark: HALLMARK_TNFA_SIGNALING_VIA_NFKB |
| Monocytes | Inflammatory response | MSigDB Hallmark: HALLMARK_INFLAMMATORY_RESPONSE |
| Monocytes | IL-6/JAK/STAT3 pathway | MSigDB Hallmark: HALLMARK_IL6_JAK_STAT3_SIGNALING |
| Monocytes | IL-1 pathway | MSigDB Reactome: REACTOME_INTERLEUKIN_1_SIGNALING |
| Monocytes | Chemokine signaling | MSigDB Reactome: REACTOME_CHEMOKINE_RECEPTORS_BIND_CHEMOKINES |
| Monocytes | Gαi signaling | MSigDB Reactome: REACTOME_G_ALPHA_I_SIGNALLING_EVENTS |
| Monocytes | IFN-α response | MSigDB Hallmark: HALLMARK_INTERFERON_ALPHA_RESPONSE |
| Monocytes | IFN-γ response | MSigDB Hallmark: HALLMARK_INTERFERON_GAMMA_RESPONSE |
| Monocytes | FcγR-mediated phagocytosis | MSigDB Reactome: REACTOME_FC_GAMMA_RECEPTOR_FCGR_DEPENDENT_PHAGOCYTOSIS |
| Monocytes | Complement | MSigDB Hallmark: HALLMARK_COMPLEMENT |
| Monocytes | MHC-II antigen presentation | MSigDB Reactome: REACTOME_MHC_CLASS_II_ANTIGEN_PRESENTATION |
| Monocytes | Cross-presentation | MSigDB Reactome: REACTOME_ANTIGEN_PROCESSING_CROSS_PRESENTATION |
| Monocytes | Hypoxia | MSigDB Hallmark: HALLMARK_HYPOXIA |
| Monocytes | Glycolysis | MSigDB Hallmark: HALLMARK_GLYCOLYSIS |
| Monocytes | mTORC1 signaling | MSigDB Hallmark: HALLMARK_MTORC1_SIGNALING |

| | | |
|---|---|---|
| Monocytes | Oxidative phosphorylation | MSigDB Hallmark: HALLMARK_OXIDATIVE_PHOSPHORYLATION |
| Monocytes | ROS pathway | MSigDB Hallmark: HALLMARK_REACTIVE_OXYGEN_SPECIES_PATHWAY |
| Monocytes | Unfolded protein response | MSigDB Hallmark: HALLMARK_UNFOLDED_PROTEIN_RESPONSE |
| Monocytes | TGF-β signaling | MSigDB Hallmark: HALLMARK_TGF_BETA_SIGNALING |
| Monocytes | Heme metabolism | MSigDB Hallmark: HALLMARK_HEME_METABOLISM |
| Monocytes | Cholesterol homeostasis | MSigDB Hallmark: HALLMARK_CHOLESTEROL_HOMEOSTASIS |
| Monocytes | IL-10 signaling | MSigDB Reactome: REACTOME_INTERLEUKIN_10_SIGNALING |
| Monocytes | Heme scavenging | MSigDB Reactome: REACTOME_SCAVENGING_OF_HEME_FROM_PLASMA |
| Monocytes | Scavenger receptor uptake | MSigDB Reactome: REACTOME_BINDING_AND_UPTAKE_OF_LIGANDS_BY_SCAVENGER_RECEPTORS |
| NK cells | NK recognition and coactivation | NCR1, NCR3, KLRK1, FCGR3A, CD226, CD244, CD2, SLAMF6 |
| NK cells | Cytokine sensing | IL12RB1, IL12RB2, IL2RB, IL2RG, IL18R1, IL18RAP |
| NK cells | JAK/STAT signaling | JAK1, JAK2, JAK3, TYK2, STAT4, STAT5A, STAT5B |
| NK cells | MAPK nuclear signaling | MSigDB Reactome: REACTOME_MAPK_TARGETS_NUCLEAR_EVENTS_MEDIATED_BY_MAP_KINASES |
| NK cells | NF-κB response | NFKBIA, NFKBIZ, NFKBIE, TNFAIP3, TNIP1, NFKB1, NFKB2, RELB, BIRC3, TRAF1, ICAM1, ZC3H12A |
| NK cells | Degranulation machinery | RAB27A, UNC13D, STX11, STXBP2, VAMP7, SNAP23, LYST, ARL8A, RAB11B, AP3B1 |
| NK cells | Cytotoxic effector program | PRF1, GNLY, GZMA, GZMB, GZMH, GZMM, NKG7, CTSW, CST7, FGFBP2 |
| NK cells | Cytokine output | IFNG, TNF, CSF2, XCL1, XCL2, CCL3, CCL4, CCL5 |
| NK cells | ISG response | ISG15, IFI6, IFI44, IFI44L, IFIT1, IFIT2, IFIT3, MX1, MX2, |

| | | |
|---|---|---|
| | | OAS1, OAS2, OAS3, OASL, RSAD2, BST2, USP18, STAT1, IRF7 |
| NK cells | Trafficking and adhesion | CXCR3, CXCR4, CX3CR1, CCR5, CCR7, S1PR1, S1PR5, SELL, ITGAL, ITGB2, ITGA4, SELPLG |
| NK cells | Inhibitory regulation | TIGIT, HAVCR2, LAG3, PDCD1, CISH, SOCS1, SOCS3, CBLB, PTPN6, INPP5D |
| NK cells | Glycolysis | MSigDB Hallmark: HALLMARK_GLYCOLYSIS |
| NK cells | Oxidative phosphorylation | MSigDB Hallmark: HALLMARK_OXIDATIVE_PHOSPHORYLATION |
| NK cells | mTORC1 response | MSigDB Hallmark: HALLMARK_MTORC1_SIGNALING |

### Table S3 | Functional enrichment analysis of differentially expressed genes in CD56$^{hi}$NK cells

| Pattern_group | ID | FoldEnrichment | p.adjust | Overlapping genes |
|---|---|---|---|---|
| HIV_UP | GO:0044194 | 127.2959184 | 4.86145E-08 | GNLY/ GZMB/ GZMH/ NKG7/ SERPINB1 |
| HIV_UP | GO:0031726 | 167.2946429 | 8.64428E-05 | CCL3/ CCL4/ CCL5 |
| HIV_UP | GO:0001906 | 11.76818182 | 9.40129E-05 | CTSC/ GAPDH/ GNLY/ GZMB/ GZMH/ HLA-C/ HLA-DRA/ IFNG/ NKG7 |
| HIV_UP | GO:0141061 | 16.70322581 | 0.00042058 | GAPDH/ GNLY/ GZMB/ GZMH/ IFNG/ NKG7 |
| HIV_UP | GO:0051702 | 16.05581395 | 0.00042058 | CCL3/ CCL4/ CCL5/ GAPDH/ PIK3C3/ RAB29 |
| HIV_UP | hsa05330 | 33.32625995 | 0.00042482 | GZMB/ HLA-C/ HLA-DRA/ IFNG |
| HIV_UP | GO:0071346 | 17.6122449 | 0.00141479 | CCL3/ CCL5/ GAPDH/ GBP4/ IFNG |
| HIV_UP | hsa04650 | 12.21545242 | 0.001515433 | GZMB/ HLA-C/ IFNG/ LAT/ RAC2 |
| HIV_UP | GO:0045910 | 26.55384615 | 0.001954184 | H1-10/ H1-3/ H1-4/ SMCHD1 |
| HIV_UP | GO:0002366 | 7.671111111 | 0.003372239 | CCL3/ HLA-DRA/ IFNG/ LAT/ LGALS1/ NKG7/ RAC2 |
| HIV_UP | GO:0061844 | 12.69117647 | 0.003853484 | CCL3/ CCL4/ CCL5/ GAPDH/ GNLY |
| HIV_UP | hsa05168 | 9.025862069 | 0.004233276 | CCL5/ HLA-C/ HLA-DRA/ IFIH1/ IFNG |
| HIV_UP | hsa05323 | 13.68130672 | 0.004233276 | CCL3/ CCL5/ HLA-DRA/ IFNG |
| HIV_UP | GO:0002443 | 5.694020619 | 0.004926827 | CCL3/ CTSC/ GZMB/ HLA-C/ HLA-DRA/ LAT/ NKG7/ RAC2 |
| HIV_UP | GO:0002274 | 8.385425101 | 0.004926827 | CCL3/ CCL5/ CTSC/ IFNG/ LAT/ RAC2 |
| HIV_UP | GO:0001909 | 11.43046358 | 0.004926827 | CTSC/ GZMB/ HLA-C/ HLA-DRA/ NKG7 |
| HIV_UP | GO:0046637 | 17.26 | 0.00516037 | HLA-DRA/ IFNG/ LGALS1/ PRDM1 |
| HIV_UP | GO:0043299 | 17.04691358 | 0.005208477 | CCL3/ LAT/ NKG7/ RAC2 |
| HIV_UP | GO:0050729 | 10.65432099 | 0.005704506 | CCL3/ CTSC/ IFNG/ LGALS1/ NKG7 |
| HIV_UP | GO:0050863 | 6.148600509 | 0.006994267 | CCL5/ HLA-DRA/ IFNG/ LAT/ LGALS1/ PRDM1/ RAC2 |
| HIV_UP | GO:0032831 | 98.62857143 | 0.008427741 | HLA-DRA/ IFNG |
| HIV_UP | GO:0002275 | 14.08979592 | 0.008859501 | CCL3/ IFNG/ LAT/ RAC2 |
| HIV_UP | hsa05163 | 7.157071244 | 0.00943361 | CCL3/ CCL4/ CCL5/ HLA-C/ RAC2 |
| HIV_UP | GO:0098553 | 30.55102041 | 0.01098338 | HLA-C/ HLA-DRA/ SPPL2A |
| HIV_UP | hsa04210 | 9.418290855 | 0.011280945 | CAPN2/ CTSC/ GZMB/ PMAIP1 |
| HIV_UP | GO:0140507 | 69.04 | 0.013578689 | GZMB/ NKG7 |
| HIV_UP | GO:1903980 | 69.04 | 0.013578689 | CCL3/ CTSC |
| HIV_UP | GO:0001774 | 21.13469388 | 0.01399342 | CCL3/ CTSC/ IFNG |
| HIV_UP | GO:0001772 | 22.27678571 | 0.014142495 | GZMB/ HLA-DRA/ LAT |
| HIV_UP | hsa05416 | 13.92561576 | 0.016113857 | HLA-C/ HLA-DRA/ RAC2 |
| HIV_UP | GO:0035710 | 10.70387597 | 0.016501777 | HLA-DRA/ IFNG/ LGALS1/ NKG7 |
| HIV_UP | HALLMARK_TNFA_SIGNALING_VIA_NFKB | 4.696071429 | 0.019007894 | BTG3/ CCL4/ CCL5/ IFIH1/ NAMPT/ PPP1R15A |
| HIV_UP | HALLMARK_INTERFERON_GAMMA_RESPONSE | 4.696071429 | 0.019007894 | CCL5/ GBP4/ IFIH1/ NAMPT/ RNF213/ SPPL2A |
| HIV_UP | GO:0031490 | 10.45591518 | 0.019037482 | H1-10/ H1-3/ H1-4/ UTY |
| HIV_UP | GO:0002696 | 5.582749326 | 0.019876908 | CCL3/ CCL5/ CTSC/ HLA-DRA/ IFNG/ LGALS1 |
| HIV_UP | GO:0005125 | 7.029186675 | 0.020110235 | CCL3/ CCL4/ CCL5/ IFNG/ NAMPT |
| HIV_UP | GO:0001913 | 16.97704918 | 0.020744987 | CTSC/ HLA-C/ HLA-DRA |

| | | | | |
|---|---|---|---|---|
| HIV_UP | GO:0045335 | 9.832512315 | 0.022599771 | GNLY/ HLA-C/ PIK3C3/ RAC2 |
| HIV_UP | GO:0044754 | 47.52380952 | 0.022599771 | HLA-DRA/ PIK3C3 |
| HIV_UP | GO:0051770 | 43.15 | 0.02488364 | IFNG/ NAMPT |
| HIV_UP | hsa04658 | 10.48164627 | 0.026020877 | HLA-DRA/ IFNG/ LAT |
| HIV_UP | R-HSA-9705683 | 6.384465575 | 0.027622638 | HLA-C/ IFIH1/ PIK3C3/ RPS4Y1/ YWHAQ |
| HIV_UP | R-HSA-5357801 | 6.084725407 | 0.027622638 | GZMB/ H1-3/ H1-4/ PMAIP1/ YWHAQ |
| HIV_UP | R-HSA-877300 | 10.8003876 | 0.027622638 | GBP4/ HLA-C/ HLA-DRA/ IFNG |
| HIV_UP | R-HSA-109704 | 17.67336152 | 0.027622638 | FRS2/ PDE3B/ PIK3C3 |
| HIV_UP | R-HSA-109606 | 14.13868922 | 0.027622638 | GZMB/ PMAIP1/ YWHAQ |
| HIV_UP | R-HSA-140342 | 39.8783542 | 0.027622638 | H1-3/ H1-4 |
| HIV_UP | hsa05203 | 6.340117746 | 0.027964929 | HLA-C/ PMAIP1/ RASA2/ YWHAQ |
| HIV_UP | GO:0005126 | 6.039517793 | 0.028839717 | CCL3/ CCL4/ CCL5/ FRS2/ IFNG |
| HIV_UP | GO:0016004 | 33.45892857 | 0.028839717 | CCL3/ CCL5 |
| HIV_UP | GO:0016032 | 4.582300885 | 0.039748549 | CCL3/ CCL4/ CCL5/ IFIH1/ LGALS1/ PIK3C3 |
| HIV_UP | GO:0042611 | 28.51428571 | 0.041247022 | HLA-C/ HLA-DRA |
| HIV_UP | GO:0006417 | 4.492841649 | 0.042717749 | CCL5/ GAPDH/ GZMB/ NIBAN1/ PPP1R15A/ XRN1 |
| HIV_UP | GO:0098586 | 11.63595506 | 0.043355323 | CCL5/ IFIH1/ IFNG |
| HIV_UP | hsa05131 | 5.137249557 | 0.045835219 | CAPN2/ CCL5/ PFN1/ PIK3C3 |
| SYPH_PEAK | HALLMARK_MITOTIC_SPINDLE | 8.009136592 | 0.015868654 | DYNC1H1/ EZR/ TRIO/ TUBA4A |
| SYPH_PEAK | R-HSA-9648025 | 19.05299145 | 0.045050001 | DYNC1H1/ PPP2R5C/ TUBA4A |
| SYPH_PEAK | R-HSA-9646399 | 34.56124031 | 0.045050001 | DYNC1H1/ TUBA4A |
| SYPH_PEAK | R-HSA-373752 | 33.77575758 | 0.045050001 | EZR/ TRIO |
| SYPH_PEAK | R-HSA-437239 | 31.61985816 | 0.045050001 | EZR/ TUBA4A |
| HIV_DOWN | R-HSA-156842 | 103.2745367 | 2.18317E-51 | EEF1A1/ EEF1B2/ RPL11/ RPL21/ RPL24/ RPL26/ RPL30/ RPL31/ RPL32/ RPL34/ RPL35A/ RPL36A/ RPL39/ RPL6/ RPL7/ RPL9/ RPLP2/ RPS15A/ RPS17/ RPS23/ RPS24/ RPS25/ RPS27/ RPS27A/ RPS3A/ RPS4X/ RPS6 |
| HIV_DOWN | GO:0022625 | 141.9630156 | 1.63514E-28 | RPL11/ RPL21/ RPL24/ RPL26/ RPL30/ RPL31/ RPL32/ RPL34/ RPL35A/ RPL36A/ RPL39/ RPL6/ RPL7/ RPL9/ RPLP2 |
| HIV_DOWN | R-HSA-9735869 | 106.8927637 | 1.34955E-19 | EEF1A1/ RPS15A/ RPS17/ RPS23/ RPS24/ RPS25/ RPS27/ RPS27A/ RPS3A/ RPS4X/ RPS6 |
| HIV_DOWN | GO:0042254 | 27.37954033 | 1.46692E-17 | EIF1AX/ RPL11/ RPL24/ RPL26/ RPL35A/ RPL7/ RPS15A/ RPS17/ RPS23/ RPS24/ RPS25/ RPS27/ RPS27A/ RPS3A/ RPS4X/ RPS6 |
| HIV_DOWN | GO:0006364 | 21.69828571 | 1.63435E-08 | RPL11/ RPL26/ RPL35A/ RPL7/ RPS17/ RPS24/ RPS25/ RPS27/ RPS6 |
| HIV_DOWN | GO:0005925 | 11.45079041 | 4.75445E-07 | RPL30/ RPL31/ RPL6/ RPL7/ RPL9/ RPLP2/ RPS17/ RPS3A/ RPS4X |
| HIV_DOWN | GO:0042273 | 42.37946429 | 6.12958E-06 | RPL11/ RPL24/ RPL26/ RPL35A/ RPL7 |
| HIV_DOWN | GO:0042255 | 37.41083744 | 0.000184067 | EIF1AX/ RPL11/ RPL24/ RPS27 |
| HIV_DOWN | HALLMARK_MYC_TARGETS_V1 | 9.13125 | 0.001605445 | EEF1B2/ EIF1AX/ RPL34/ RPL6/ RPS6 |
| HIV_DOWN | GO:0014069 | 7.513362945 | 0.003313913 | RPL30/ RPL6/ RPL7/ RPS25/ RPS27 |
| HIV_DOWN | GO:0019843 | 23.01842752 | 0.007477316 | RPL11/ RPL9/ RPS4X |
| HIV_DOWN | GO:0000049 | 18.75575576 | 0.007477316 | EEF1A1/ EIF1AX/ RPL35A |
| HIV_DOWN | GO:0008135 | 18.30380983 | 0.007477316 | EEF1A1/ EEF1B2/ EIF1AX |

| | | | | |
|---|---|---|---|---|
| HIV_DOWN | GO:0003746 | 56.26726727 | 0.007477316 | EEF1A1/ EEF1B2 |
| HIV_DOWN | GO:0048027 | 42.20045045 | 0.010868577 | RPL26/ RPS3A |
| HIV_DOWN | GO:0006414 | 22.60238095 | 0.01341932 | EEF1A1/ EEF1B2/ RPLP2 |
| HIV_DOWN | R-HSA-9629569 | 35.95483871 | 0.015674051 | RPS23/ RPS6 |
| HIV_DOWN | GO:0000027 | 63.81848739 | 0.016988474 | RPL11/ RPL24 |
| HIV_DOWN | R-HSA-9613829 | 32.68621701 | 0.018503533 | EEF1A1/ RPS27A |
| HIV_DOWN | GO:0005791 | 18.60205033 | 0.025887104 | RPL6/ RPS23 |
| HIV_DOWN | GO:1901798 | 34.99723502 | 0.047626987 | RPL11/ RPL26 |
| HIV_DOWN | GO:0006413 | 12.81394826 | 0.048454597 | EIF1AX/ RPS17/ RPS3A |
| INFECTION_DOWN | GO:0022626 | 124.0451977 | 4.34952E-21 | RPL17/ RPL22/ RPL23/ RPL23A/ RPL27/ RPL37/ RPL37A/ RPL41/ RPS21/ RPS26/ RPS29 |
| INFECTION_DOWN | GO:0098556 | 665.3333333 | 1.94306E-08 | RPL27/ RPS26/ RPS29 |
| INFECTION_DOWN | GO:0022627 | 95.04761905 | 6.58409E-06 | RPS21/ RPS26/ RPS29 |
| INFECTION_DOWN | GO:0005925 | 15.6918239 | 1.64466E-05 | RPL22/ RPL23/ RPL27/ RPL37A/ RPS29 |
| INFECTION_DOWN | GO:0019843 | 56.77878788 | 0.000304054 | RPL23/ RPL23A/ RPL37 |
| INFECTION_DOWN | GO:1990948 | 227.1151515 | 0.000360404 | RPL23/ RPL37 |
| INFECTION_DOWN | R-HSA-198933 | 11.97207304 | 0.013606352 | KLRB1/ SELL |
| INFECTION_DOWN | HALLMARK_INFLAMMATORY_RESPONSE | 14.61 | 0.030146991 | GPR183/ SELL |
| INFECTION_DOWN | GO:0008201 | 14.19469697 | 0.031169211 | RPL22/ SELL |
| INFECTION_DOWN | GO:0045296 | 7.479840319 | 0.047096138 | RPL23A/ RPS26 |
| SYPH_DROP | GO:0050851 | 17.52194422 | 1.96415E-06 | CD3E/ NFKB1/ NFKBIA/ NFKBIZ/ TRBC1/ TRDC/ HLA-DQB1/ NFKBID/ TRGC1 |
| SYPH_DROP | GO:0042106 | 216.9565217 | 3.55676E-05 | CD3E/ TRDC/ TRGC1 |
| SYPH_DROP | hsa04658 | 22.02664797 | 0.000307118 | CD3E/ NFKB1/ NFKBIA/ HLA-DQB1/ IL12RB2 |
| SYPH_DROP | hsa05171 | 10.2852465 | 0.001098143 | FAU/ NFKB1/ NFKBIA/ RPL10A/ RPL38/ RPS8 |
| SYPH_DROP | GO:2000516 | 34.39492754 | 0.00121565 | ANXA1/ CD3E/ NFKBIZ/ NFKBID |
| SYPH_DROP | R-HSA-202424 | 18.34430546 | 0.002144023 | CD3E/ NFKB1/ NFKBIA/ PSMA1/ HLA-DQB1 |
| SYPH_DROP | R-HSA-9010553 | 10.451988 | 0.004221163 | FAU/ PSMA1/ RPL10A/ RPL38/ RPS8 |
| SYPH_DROP | R-HSA-9692914 | 14.9811828 | 0.004221163 | FAU/ NFKB1/ NFKBIA/ RPS8 |
| SYPH_DROP | R-HSA-5603029 | 102.7281106 | 0.004221163 | NFKB1/ NFKBIA |
| SYPH_DROP | hsa05166 | 9.144992236 | 0.004303471 | CD3E/ NFKB1/ NFKBIA/ TSPO/ HLA-DQB1 |
| SYPH_DROP | hsa05145 | 14.63198758 | 0.004303471 | BIRC3/ NFKB1/ NFKBIA/ HLA-DQB1 |
| SYPH_DROP | GO:0044391 | 11.66432913 | 0.005222516 | FAU/ RBM3/ RPL10A/ RPL38/ RPS8 |
| SYPH_DROP | R-HSA-1169091 | 16.09918151 | 0.009749949 | NFKB1/ NFKBIA/ PSMA1 |
| SYPH_DROP | GO:0042105 | 72.31884058 | 0.010171503 | CD3E/ TRBC1 |
| SYPH_DROP | hsa05321 | 18.62252964 | 0.010955033 | NFKB1/ HLA-DQB1/ IL12RB2 |
| SYPH_DROP | R-HSA-202430 | 37.84719864 | 0.012791008 | CD3E/ HLA-DQB1 |
| SYPH_DROP | GO:0043403 | 28.14130435 | 0.015346687 | ANXA1/ EZH2/ HOPX |
| SYPH_DROP | hsa05235 | 13.65652174 | 0.016232842 | CD3E/ NFKB1/ NFKBIA |
| SYPH_DROP | GO:2000321 | 68.78985507 | 0.024588285 | NFKBIZ/ NFKBID |
| SYPH_DROP | R-HSA-9735869 | 19.43504795 | 0.029353049 | FAU/ RPS8 |
| SYPH_DROP | GO:0050727 | 5.653960691 | 0.031798435 | ANXA1/ BIRC3/ NFKB1/ NFKBIA/ NFKBIZ/ PSMA1 |
| SYPH_DROP | GO:1905953 | 18.20907928 | 0.031798435 | NFKB1/ NFKBIA/ TSPO |
| SYPH_DROP | GO:0030216 | 9.275036639 | 0.034394378 | ANXA1/ EZH2/ NFKBIZ/ FLNB |

| | | | | |
|---|---|---|---|---|
| SYPH_DROP | GO:0050853 | 15.67363786 | 0.034394378 | NFKB1/ NFKBIA/ NFKBIZ |
| SYPH_DROP | GO:0097398 | 45.85990338 | 0.034394378 | NFKB1/ NFKBIZ |
| SYPH_DROP | GO:1903708 | 8.466443701 | 0.03480002 | ANXA1/ KLF10/ NFKBIZ/ NFKBID |
| SYPH_DROP | GO:0050810 | 14.39787664 | 0.03480002 | IGFBP7/ NFKB1/ TSPO |
| SYPH_DROP | GO:0045063 | 34.39492754 | 0.038998695 | ANXA1/ NFKBIZ |
| SYPH_DROP | GO:0032496 | 5.829648735 | 0.03991197 | NFKB1/ NFKBIA/ NFKBIZ/ TSPO/ IL12RB2 |
| SYPH_DROP | R-HSA-449147 | 3.80072290799973 | 0.047249683 | ANXA1/ NFKB1/ NFKBIA/ PSMA1/ IL12RB2 |
| SYPH_DROP | GO:0035872 | 28.46476762 | 0.048169373 | BIRC3/ NFKBIA |

### Table S4 | Functional enrichment analysis of differentially expressed genes in CD56$^{dim}$NK cells

| Pattern_group | ID | FoldEnrichment | p.adjust | Overlapping genes |
|---|---|---|---|---|
| HIV_UP | HALLMARK_INTERFERON_GAMMA_RESPONSE | 5.229715909 | 7.856584093e-09 | ARID5B/ CCL5/ CD38/ CD74/ DDX60/ EIF2AK2/ EIF4E3/ EPSTI1/ FAS/ HLA-DRB1/ IRF1/ ISG15/ ITGB7/ JAK2/ LY6E/ PSMB8/ RNF213/ SAMD9L/ STAT1/ TRIM26/ UBE2L6 |
| HIV_UP | hsa05332 | 16.752 | 5.941981386e-08 | FAS/ GZMB/ HLA-C/ HLA-DPA1/ HLA-DPB1/ HLA-DRA/ HLA-DRB1/ HLA-E/ KIR2DL1/ KIR3DL1 |
| HIV_UP | hsa05168 | 6.7008 | 1.925347718e-07 | BIRC3/ CCL5/ CD74/ EIF2AK2/ FAS/ HLA-C/ HLA-DPA1/ HLA-DPB1/ HLA-DRA/ HLA-DRB1/ HLA-E/ JAK2/ PPP1CA/ PPP1CC/ STAT1/ TAB2 |
| HIV_UP | GO:0030134 | 9.589866783 | 2.707254461e-06 | CD74/ ECPAS/ HLA-C/ HLA-DPA1/ HLA-DPB1/ HLA-DRA/ HLA-DRB1/ HLA-E/ SEC24B/ SEC24D/ TMED9 |
| HIV_UP | GO:0098553 | 18.93064612 | 2.707254461e-06 | BCAP31/ CD74/ HLA-C/ HLA-DPA1/ HLA-DPB1/ HLA-DRA/ HLA-DRB1/ HLA-E |
| HIV_UP | hsa04145 | 4.928953846 | 3.009731924e-06 | ATG7/ ATP6V0B/ CD63/ CORO1A/ CTSD/ DCTN3/ HLA-C/ HLA-DPA1/ HLA-DPB1/ HLA-DRA/ HLA-DRB1/ HLA-E/ ITGB1/ M6PR/ RAB5C/ TUBA1B/ TUBB |
| HIV_UP | GO:0030666 | 6.001563534 | 3.506500022e-06 | ARRB2/ ATP6V0B/ CAMK2D/ CD74/ CORO1A/ HLA-C/ HLA-DPA1/ HLA-DPB1/ HLA-DRA/ HLA-DRB1/ HLA-E/ M6PR/ MDM2/ RAB10/ RAB11B |
| HIV_UP | GO:0030135 | 4.658713693 | 3.707706972e-06 | ATP6V0B/ BCAP31/ CD74/ DENND1B/ ECPAS/ HLA-C/ HLA-DPA1/ HLA-DPB1/ HLA-DRA/ HLA-DRB1/ HLA-E/ M6PR/ MYO6/ NECAP2/ SEC24B/ SEC24D/ SYT11/ TMED9 |
| HIV_UP | hsa04612 | 9.193170732 | 4.393156288e-06 | CD74/ HLA-C/ HLA-DPA1/ HLA-DPB1/ HLA-DRA/ HLA-DRB1/ HLA-E/ KIR2DL1/ KIR3DL1/ KLRC3 |
| HIV_UP | HALLMARK_INTERFERON_ALPHA_RESPONSE | 6.161668229 | 8.360022296e-06 | CD74/ DDX60/ EIF2AK2/ EPSTI1/ HLA-C/ IRF1/ ISG15/ LY6E/ PSMB8/ SAMD9L/ TRIM26/ UBE2L6 |
| HIV_UP | hsa05140 | 8.698153846 | 2.361173524e-05 | HLA-DPA1/ HLA-DPB1/ HLA-DRA/ HLA-DRB1/ ITGA4/ ITGB1/ JAK2/ STAT1/ TAB2 |
| HIV_UP | hsa05165 | 3.848432432 | 4.503656451e-05 | ATP6V0B/ BCAP31/ CCND2/ EIF2AK2/ FAS/ HLA-C/ HLA-E/ IRF1/ ISG15/ ITGA4/ ITGB1/ ITGB7/ MAML2/ MDM2/ PKM/ PRKACB/ STAT1 |
| HIV_UP | GO:0042613 | 24.35928728 | 5.426421097e-05 | CD74/ HLA-DPA1/ HLA-DPB1/ HLA-DRA/ HLA-DRB1 |
| HIV_UP | hsa05169 | 4.78044878 | 6.427858787e-05 | CCND2/ EIF2AK2/ FAS/ HLA-C/ HLA-DPA1/ HLA-DPB1/ HLA-DRA/ HLA-DRB1/ HLA-E/ ISG15/ MDM2/ STAT1/ TAB2 |
| HIV_UP | hsa04658 | 7.295225806 | 7.025768964e-05 | GATA3/ HLA-DPA1/ HLA-DPB1/ HLA-DRA/ HLA-DRB1/ JAK2/ MAML2/ STAT1/ TBX21 |
| HIV_UP | hsa05152 | 4.997834254 | 8.440104772e-05 | ATP6V0B/ CAMK2D/ CD74/ CORO1A/ CTSD/ HLA-DPA1/ HLA-DPB1/ HLA-DRA/ HLA-DRB1/ JAK2/ RAB5C/ STAT1 |
| HIV_UP | GO:0002478 | 15.89950801 | 0.0001007412159 | CD74/ CTSD/ HLA-DPA1/ HLA-DPB1/ HLA-DRA/ HLA-DRB1/ HLA-E/ MFSD6 |
| HIV_UP | R-HSA-877300 | 7.602058532 | 0.000161355409 | CAMK2D/ HLA-C/ HLA-DPA1/ HLA-DPB1/ HLA-DRA/ HLA-DRB1/ HLA-E/ IRF1/ JAK2/ STAT1/ TRIM26 |
| HIV_UP | GO:0046634 | 7.347006262 | 0.0002089084876 | CD300A/ GATA3/ HLA-DRA/ HLA-DRB1/ HLA-E/ IRF1/ JAK2/ LGALS1/ OPA1/ RASAL3/ TBX21 |
| HIV_UP | GO:0007159 | 3.789999002 | 0.0002091933095 | CCL5/ CD300A/ CD74/ CORO1A/ GATA3/ HLA-DPA1/ HLA-DPB1/ HLA-DRA/ HLA-DRB1/ HLA-E/ IRF1/ ITGA4/ ITGB1/ ITGB7/ JAK2/ LGALS1/ OPA1/ RASAL3/ TBX21/ UFL1 |

| | | | | |
|---|---|---|---|---|
| HIV_UP | GO:0030665 | 6.001563534 | 0.0002338684038 | ATP6V0B/ CD74/ HLA-DPA1/ HLA-DPB1/ HLA-DRA/ HLA-DRB1/ M6PR/ MYO6/ NECAP2/ SYT11 |
| HIV_UP | R-HSA-913531 | 4.192821738 | 0.0002553285519 | CAMK2D/ EIF2AK2/ EIF4E3/ HLA-C/ HLA-DPA1/ HLA-DPB1/ HLA-DRA/ HLA-DRB1/ HLA-E/ IRF1/ ISG15/ JAK2/ PSMB8/ STAT1/ TRIM26/ TUBA1B/ UBE2L6 |
| HIV_UP | hsa05164 | 4.793202312 | 0.0002957269527 | CCL5/ EIF2AK2/ FAS/ HLA-DPA1/ HLA-DPB1/ HLA-DRA/ HLA-DRB1/ JAK2/ RAB11B/ SLC25A6/ STAT1 |
| HIV_UP | GO:0044194 | 23.66330765 | 0.0005820161772 | GNLY/ GZMB/ GZMH/ NKG7 |
| HIV_UP | GO:0050792 | 5.65213724 | 0.0006090505224 | APOBEC3C/ BSG/ CCL5/ CD74/ EIF2AK2/ HLA-DRB1/ ISG15/ LGALS1/ LY6E/ MDFIC/ STAT1/ TRIM26 |
| HIV_UP | hsa04514 | 4.7115 | 0.0006902204857 | CD99/ HLA-C/ HLA-DPA1/ HLA-DPB1/ HLA-DRA/ HLA-DRB1/ HLA-E/ ITGA4/ ITGB1/ ITGB7 |
| HIV_UP | hsa04672 | 9.04608 | 0.0006902204857 | HLA-DPA1/ HLA-DPB1/ HLA-DRA/ HLA-DRB1/ ITGA4/ ITGB7 |
| HIV_UP | GO:0001909 | 5.935991814 | 0.0007957927397 | ARRB2/ CORO1A/ GZMB/ HLA-C/ HLA-DRA/ HLA-DRB1/ HLA-E/ KIR3DL1/ KLRC3/ NKG7/ TUBB |
| HIV_UP | GO:0002819 | 4.793234032 | 0.0009727427834 | DENND1B/ GATA3/ HLA-C/ HLA-DRA/ HLA-DRB1/ HLA-E/ IRF1/ JAK2/ LGALS1/ OPA1/ SAMSN1/ TBX21/ UFL1 |
| HIV_UP | GO:0030133 | 3.108094492 | 0.001118224437 | CD74/ ERP29/ HLA-C/ HLA-DPA1/ HLA-DPB1/ HLA-DRA/ HLA-DRB1/ HLA-E/ M6PR/ RAB10/ RAB11B/ RAB1B/ RAB5C/ SEC24B/ SEC24D/ SYT11/ TMED9 |
| HIV_UP | GO:0051251 | 3.637722256 | 0.002886225627 | CCL5/ CD38/ CD74/ CORO1A/ GATA3/ HLA-DPA1/ HLA-DPB1/ HLA-DRA/ HLA-DRB1/ HLA-E/ JAK2/ LGALS1/ OPA1/ RASAL3/ TBX21 |
| HIV_UP | GO:0002449 | 3.377615691 | 0.00341857402 | ARRB2/ CD74/ CORO1A/ DENND1B/ GATA3/ GZMB/ HLA-C/ HLA-DRA/ HLA-DRB1/ HLA-E/ KIR3DL1/ KLRC3/ NKG7/ TBX21/ TUBB/ UFL1 |
| HIV_UP | GO:0045622 | 10.18562232 | 0.00341857402 | GATA3/ HLA-DRA/ HLA-DRB1/ LGALS1/ OPA1/ TBX21 |
| HIV_UP | GO:0075294 | 21.72932761 | 0.00341857402 | BSG/ CD74/ HLA-DRB1/ LGALS1 |
| HIV_UP | GO:0048525 | 6.584644731 | 0.003629287785 | APOBEC3C/ CCL5/ CD74/ EIF2AK2/ ISG15/ LY6E/ STAT1/ TRIM26 |
| HIV_UP | hsa05130 | 3.750447761 | 0.003860121971 | FAS/ GAPDH/ ITGB1/ MYO5A/ MYO6/ SEC24B/ SEC24D/ TAB2/ TUBA1B/ TUBB |
| HIV_UP | GO:0002711 | 7.605264664 | 0.003954987324 | DENND1B/ GATA3/ HLA-C/ HLA-DRA/ HLA-DRB1/ HLA-E/ TBX21 |
| HIV_UP | GO:0046596 | 9.586468064 | 0.003954987324 | BSG/ CD74/ HLA-DRB1/ LGALS1/ LY6E/ TRIM26 |
| HIV_UP | GO:0019883 | 12.7320279 | 0.004067561718 | CD74/ HLA-C/ HLA-DRA/ HLA-DRB1/ HLA-E |
| HIV_UP | GO:0050670 | 4.074248927 | 0.004175763985 | CCL5/ CD300A/ CD38/ CD74/ CORO1A/ HLA-DPA1/ HLA-DPB1/ HLA-DRB1/ HLA-E/ IRF1/ JAK2/ RASAL3 |
| HIV_UP | R-HSA-2132295 | 5.393921796 | 0.004503889311 | CD74/ CTSD/ DCTN3/ HLA-DPA1/ HLA-DPB1/ HLA-DRA/ HLA-DRB1/ SEC24B/ SEC24D/ TUBA1B |
| HIV_UP | GO:0022408 | 4.330119633 | 0.004561323503 | CD300A/ CD74/ HLA-DRB1/ HLA-E/ IRF1/ JAK2/ LGALS1/ MAP2K5/ TBX21/ TMX1/ UFL1 |
| HIV_UP | GO:0030658 | 3.586761198 | 0.006160659644 | CD74/ HLA-C/ HLA-DPA1/ HLA-DPB1/ HLA-DRA/ HLA-DRB1/ HLA-E/ RAB11B/ RAB5C/ SEC24B/ SEC24D |
| HIV_UP | GO:0055037 | 3.943884608 | 0.006160659644 | HLA-C/ HLA-E/ ITGB1/ MCOLN2/ MYO5A/ RAB10/ RAB11B/ RAB11FIP2/ RELCH/ SYT11 |
| HIV_UP | GO:0019774 | 22.58770275 | 0.006160659644 | PSMB3/ PSMB4/ PSMB8 |
| HIV_UP | GO:0050866 | 4.092852803 | 0.006309580051 | CD300A/ CD74/ HLA-DRB1/ HLA-E/ IRF1/ LGALS1/ SAMSN1/ SYT11/ TBX21/ TMX1/ UFL1 |

| | | | | |
|---|---|---|---|---|
| HIV_UP | GO:0005765 | 2.779247542 | 0.006536814171 | ATP6V0B/ CD63/ CD74/ CTSD/ HLA-DPA1/ HLA-DPB1/ HLA-DRA/ HLA-DRB1/ M6PR/ MYO6/ NKG7/ RAB5C/ RNF13/ SYT11/ TAB2 |
| HIV_UP | GO:0045335 | 4.569466304 | 0.006536814171 | ATP6V0B/ CORO1A/ GNLY/ HLA-C/ HLA-E/ RAB10/ RAB11B/ SYT11 |
| HIV_UP | GO:0042470 | 5.176348548 | 0.006536814171 | BSG/ CD63/ CTSD/ ERP29/ ITGB1/ MYO5A/ RAB5C |
| HIV_UP | R-HSA-198933 | 4.988363767 | 0.006677052304 | CD300A/ CD99/ HLA-C/ HLA-E/ ITGA4/ ITGB1/ ITGB7/ KIR2DL1/ KIR3DL1/ KLRG1 |
| HIV_UP | GO:0043122 | 3.473130233 | 0.007072967741 | ARRB2/ BIRC3/ CD74/ GAPDH/ HLA-DRB1/ LGALS1/ LIME1/ MAP2K5/ RHOC/ SHISA5/ STAT1/ TAB2/ UFL1 |
| HIV_UP | GO:0002699 | 3.594925524 | 0.008456126744 | CD300A/ CD74/ DDX60/ DENND1B/ GATA3/ HLA-C/ HLA-DRA/ HLA-DRB1/ HLA-E/ KLRC3/ OPA1/ TBX21 |
| HIV_UP | GO:0048524 | 7.521690327 | 0.008456126744 | BSG/ CCL5/ CD74/ HLA-DRB1/ LGALS1/ MDFIC |
| HIV_UP | GO:0019058 | 3.27958118 | 0.01028584817 | APOBEC3C/ BSG/ CCL5/ CD74/ EIF2AK2/ HLA-DRB1/ ISG15/ ITGB1/ ITGB7/ LGALS1/ LY6E/ RAB1B/ TRIM26 |
| HIV_UP | hsa04921 | 3.916051948 | 0.01060629562 | CAMK2D/ CD38/ EEF2/ GNAI2/ MAP2K5/ PPP1CA/ PPP1CC/ PRKACB |
| HIV_UP | R-HSA-5357801 | 3.737759893 | 0.01081156124 | BCAP31/ BIRC3/ FAS/ GZMB/ H1-2/ H1-4/ IRF1/ OPA1/ PSMB3/ PSMB4/ PSMB8/ SPTAN1 |
| HIV_UP | R-HSA-380320 | 5.646403242 | 0.01081156124 | CEP290/ CEP57/ CEP78/ DCTN3/ MZT2A/ NEDD1/ TUBA1B/ TUBB |
| HIV_UP | R-HSA-983170 | 11.43883415 | 0.01081156124 | BCAP31/ HLA-C/ HLA-E/ SEC24B/ SEC24D |
| HIV_UP | R-HSA-199977 | 4.280337942 | 0.01085741285 | COG5/ DCTN3/ GOLGB1/ RAB1B/ SEC24B/ SEC24D/ SPTAN1/ TMED9/ TRAPPC1/ TUBA1B |
| HIV_UP | R-HSA-202424 | 5.415937804 | 0.01085741285 | HLA-DPA1/ HLA-DPB1/ HLA-DRA/ HLA-DRB1/ PSMB3/ PSMB4/ PSMB8/ TAB2 |
| HIV_UP | R-HSA-5663202 | 2.573737685 | 0.01207462573 | AGK/ AGO3/ ARRB2/ ATG7/ CAMK2D/ ERLEC1/ GOLGB1/ GZMB/ JAK2/ MAML2/ MDM2/ MIB1/ PPP1CC/ PSMB3/ PSMB4/ PSMB8/ RNF213/ STAT1 |
| HIV_UP | R-HSA-909733 | 6.031385281 | 0.01207462573 | EIF2AK2/ HLA-C/ HLA-E/ IRF1/ ISG15/ PSMB8/ STAT1 |
| HIV_UP | GO:0050777 | 3.936472393 | 0.01214427501 | ARRB2/ CD300A/ HLA-DRB1/ HLA-E/ ISG15/ LGALS1/ PSMB4/ SAMSN1/ TBX21/ UFL1 |
| HIV_UP | GO:0098791 | 2.753658562 | 0.01318789915 | BCAP31/ CD74/ COG5/ GOLGA8A/ GOLGB1/ HLA-DPA1/ HLA-DPB1/ HLA-DRA/ HLA-DRB1/ M6PR/ RAB10/ RELCH/ SYT11 |
| HIV_UP | GO:1905368 | 4.494194088 | 0.01318789915 | CAPNS1/ ECPAS/ PSMB3/ PSMB4/ PSMB8/ SUPT3H/ TAF10 |
| HIV_UP | GO:0042102 | 5.484565863 | 0.01388228272 | CCL5/ CORO1A/ HLA-DPA1/ HLA-DPB1/ HLA-E/ JAK2/ RASAL3 |
| HIV_UP | GO:0046635 | 6.518798283 | 0.01437375335 | HLA-DRA/ HLA-DRB1/ HLA-E/ JAK2/ OPA1/ RASAL3 |
| HIV_UP | hsa05203 | 3.309541463 | 0.01570159709 | CCND2/ EIF2AK2/ GTF2E2/ HLA-C/ HLA-E/ MDM2/ PKM/ PRKACB/ RASA2 |
| HIV_UP | hsa05323 | 4.761094737 | 0.01570159709 | ATP6V0B/ CCL5/ HLA-DPA1/ HLA-DPB1/ HLA-DRA/ HLA-DRB1 |
| HIV_UP | GO:0002703 | 3.474165752 | 0.01596188565 | ARRB2/ CD300A/ DENND1B/ GATA3/ HLA-C/ HLA-DRA/ HLA-DRB1/ HLA-E/ KLRC3/ TBX21/ UFL1 |
| HIV_UP | hsa05132 | 3.02746988 | 0.01611222043 | BIRC3/ DCTN3/ GAPDH/ M6PR/ MYO6/ RAB5C/ S100A10/ TAB2/ TUBA1B/ TUBB |
| HIV_UP | GO:0051133 | 20.37124464 | 0.01615589069 | CD300A/ JAK2/ RASAL3 |

| | | | | |
|---|---|---|---|---|
| HIV_UP | R-HSA-5683057 | 2.857948718 | 0.01676213154 | AGO3/ ARRB2/ CAMK2D/ JAK2/ PPP1CC/ PRKACB/ PSMB3/ PSMB4/ PSMB8/ PTPRA/ RASA2/ RASA3/ RASAL3/ SPTAN1 |
| HIV_UP | hsa04650 | 3.967578947 | 0.017262208 | FAS/ GZMB/ HLA-C/ HLA-E/ KIR2DL1/ KIR3DL1/ KLRC3 |
| HIV_UP | R-HSA-9820952 | 4.423015873 | 0.01732603569 | BCAP31/ EIF2AK2/ ISG15/ MED10/ PPP1CA/ PPP1CC/ RAB5C/ UBE2L6 |
| HIV_UP | GO:0051223 | 2.740526184 | 0.01770144883 | ACSL4/ BCAP31/ BSG/ CCL5/ CD38/ CEP290/ EFNA5/ ERLEC1/ ERP29/ HLA-DRB1/ JAK2/ KIF20B/ MDFIC/ MDM2/ OAZ1 |
| HIV_UP | GO:0072676 | 5.185407725 | 0.01770144883 | CCL5/ CD99/ CORO1A/ GATA3/ ITGA4/ ITGB7/ TBX21 |
| HIV_UP | GO:2001233 | 2.830743671 | 0.01882181598 | AATF/ BCAP31/ CD74/ CDKN2D/ ERP29/ FAS/ GNAI2/ JAK2/ MAP2K5/ MDM2/ OPA1/ PPP1CA/ PRELID1/ SLC25A6 |
| HIV_UP | GO:0035745 | 18.80422582 | 0.01882181598 | DENND1B/ GATA3/ TBX21 |
| HIV_UP | R-HSA-5658442 | 5.853991597 | 0.01901857586 | PSMB3/ PSMB4/ PSMB8/ RASA2/ RASA3/ RASAL3 |
| HIV_UP | GO:0032386 | 3.295348397 | 0.02119137801 | BCAP31/ CEP290/ ERLEC1/ JAK2/ KIF20B/ MDFIC/ MDM2/ OAZ1/ PPP1CC/ RAB11B/ SEC24B |
| HIV_UP | GO:0005770 | 2.864897309 | 0.02127768525 | CD63/ CD74/ ECPAS/ HLA-DPA1/ HLA-DPB1/ HLA-DRA/ HLA-DRB1/ M6PR/ MCOLN2/ MYO5A/ RNF13 |
| HIV_UP | GO:0006417 | 2.65135505 | 0.02127953688 | AGO3/ CCL5/ EEF2/ EIF2AK2/ EIF4E3/ ELP4/ ENC1/ GAPDH/ GTDC1/ GZMB/ NIBAN1/ PATL2/ PKM/ PPP1CA/ SECISBP2 |
| HIV_UP | GO:0045580 | 3.839606319 | 0.02127953688 | CD74/ GATA3/ HLA-DRA/ HLA-DRB1/ IRF1/ LGALS1/ OPA1/ PRELID1/ TBX21 |
| HIV_UP | GO:2000328 | 17.46106683 | 0.02127953688 | LGALS1/ OPA1/ TBX21 |
| HIV_UP | hsa05166 | 3.028821429 | 0.02435078505 | CCND2/ HLA-C/ HLA-DPA1/ HLA-DPB1/ HLA-DRA/ HLA-DRB1/ HLA-E/ PRKACB/ SLC25A6 |
| HIV_UP | hsa05012 | 2.802379182 | 0.02442120503 | CAMK2D/ GNAI2/ NDUFB7/ PRKACB/ PSMB3/ PSMB4/ SLC25A6/ TUBA1B/ TUBB/ UBE2L6 |
| HIV_UP | GO:0034341 | 4.714007023 | 0.02551095418 | CCL5/ CD74/ GAPDH/ HLA-DPA1/ IRF1/ JAK2/ STAT1 |
| HIV_UP | GO:0044233 | 11.29385138 | 0.0266465685 | ACSL4/ BCAP31/ TMX1 |
| HIV_UP | GO:0002468 | 15.27843348 | 0.03004419835 | CD74/ HLA-DRA/ HLA-DRB1 |
| HIV_UP | GO:0001772 | 6.901798064 | 0.03047242482 | CORO1A/ GZMB/ HLA-DRA/ HLA-DRB1 |
| HIV_UP | GO:0072357 | 23.66330765 | 0.03174822524 | PPP1CA/ PPP1CC |
| HIV_UP | GO:0032593 | 23.66330765 | 0.03174822524 | MYO5A/ RAB10 |
| HIV_UP | GO:0030073 | 3.560023334 | 0.03286652514 | ACSL4/ CCL5/ CD38/ CYB5R4/ EFNA5/ HLA-DRB1/ JAK2/ RAB11B/ RAB11FIP2 |
| HIV_UP | GO:0072175 | 4.456209764 | 0.03286652514 | CEP290/ GATA3/ KIF20B/ MIB1/ OPA1/ PRKACB/ SEC24B |
| HIV_UP | GO:0045058 | 6.467061789 | 0.03309189198 | CD74/ GATA3/ LGALS1/ OPA1/ TBX21 |
| HIV_UP | GO:0032395 | 23.42125 | 0.03431327704 | HLA-DPA1/ HLA-DRA/ HLA-DRB1 |
| HIV_UP | GO:0090316 | 5.146419697 | 0.03433072977 | BCAP31/ CEP290/ JAK2/ KIF20B/ MDM2/ OAZ1 |
| HIV_UP | GO:0034121 | 8.577366162 | 0.0350238816 | ARRB2/ BIRC3/ CD300A/ IRF1 |
| HIV_UP | GO:0046677 | 8.577366162 | 0.0350238816 | CRIP1/ CYB5R4/ JAK2/ MDM2 |
| HIV_UP | GO:0051222 | 3.170621733 | 0.03699722305 | ACSL4/ BCAP31/ BSG/ CD38/ CEP290/ HLA-DRB1/ JAK2/ KIF20B/ MDM2/ OAZ1 |
| HIV_UP | GO:0001910 | 4.988876237 | 0.03699722305 | ARRB2/ HLA-C/ HLA-DRA/ HLA-DRB1/ HLA-E/ KLRC3 |
| HIV_UP | GO:0033631 | 13.58082976 | 0.03699722305 | CCL5/ ITGA4/ ITGB1 |

| | | | | |
|---|---|---|---|---|
| HIV_UP | GO:0002830 | 13.58082976 | 0.03699722305 | CD74/ DENND1B/ GATA3 |
| HIV_UP | GO:0042608 | 21.29204545 | 0.03738798563 | HLA-DRA/ HLA-DRB1/ HLA-E |
| HIV_UP | hsa04218 | 3.361070064 | 0.03775397498 | CCND2/ HLA-C/ HLA-E/ MDM2/ PPP1CA/ PPP1CC/ SLC25A6 |
| HIV_UP | R-HSA-9679506 | 2.403812974 | 0.03874531034 | ARID4A/ CHD3/ HLA-C/ HLA-E/ ISG15/ ITGA4/ ITGB1/ JAK2/ RPS4Y1/ SEC24B/ SEC24D/ STAT1/ TAB2/ TUBB/ ZCRB1 |
| HIV_UP | GO:0070375 | 32.59399142 | 0.0393478786 | MAP2K5/ MEF2A |
| HIV_UP | GO:0071455 | 32.59399142 | 0.0393478786 | ATG7/ FAS |
| HIV_UP | hsa05202 | 3.000358209 | 0.03955031284 | BIRC3/ CCND2/ GZMB/ ITGB7/ MDM2/ MLLT3/ PBX3/ SUPT3H |
| HIV_UP | R-HSA-68886 | 2.386519356 | 0.03974277189 | CEP290/ CEP57/ CEP78/ DCTN3/ MZT2A/ NEDD1/ PPP1CC/ PSMB3/ PSMB4/ PSMB8/ RAB1B/ SMC4/ TUBA1B/ TUBB/ VRK2 |
| HIV_UP | R-HSA-6802957 | 4.738945578 | 0.03974277189 | AGK/ ARRB2/ ATG7/ CAMK2D/ JAK2/ PPP1CC |
| HIV_UP | hsa05205 | 2.956235294 | 0.04183083421 | CAMK2D/ CD63/ FAS/ ITGB1/ MDM2/ PPP1CA/ PPP1CC/ PRKACB |
| HIV_UP | GO:0032154 | 6.134931612 | 0.04312112688 | DCTN3/ ITGB1/ PPP1CC/ RHOC |
| HIV_UP | R-HSA-983169 | 2.437882765 | 0.04340369631 | ATG7/ BCAP31/ HERC4/ HLA-C/ HLA-E/ PSMB3/ PSMB4/ PSMB8/ RNF213/ SEC24B/ SEC24D/ UBA6/ UBE2L6/ UFL1 |
| HIV_UP | GO:0002704 | 5.73837877 | 0.04495982753 | ARRB2/ CD300A/ HLA-E/ TBX21/ UFL1 |
| HIV_UP | GO:0033194 | 12.22274678 | 0.04495982753 | CD38/ JAK2/ OXR1 |
| HIV_UP | GO:0035455 | 12.22274678 | 0.04495982753 | EIF2AK2/ GATA3/ RO60 |
| HIV_UP | GO:0034451 | 3.912830398 | 0.04568386424 | CCDC14/ CEP290/ CEP57/ CHD3/ GPR174/ MIB1 |
| HIV_UP | GO:0001891 | 8.873740368 | 0.04568386424 | CORO1A/ RAB11FIP2/ SYT11 |
| HIV_UP | GO:0046640 | 7.579998004 | 0.04587502059 | HLA-E/ IRF1/ JAK2/ RASAL3 |
| HIV_UP | hsa04022 | 3.178843373 | 0.04633936372 | GNAI2/ GTF2I/ MEF2A/ PDE3B/ PPP1CA/ PPP1CC/ SLC25A6 |
| HIV_UP | hsa04217 | 3.141 | 0.04656726811 | BIRC3/ CAMK2D/ EIF2AK2/ FAS/ JAK2/ SLC25A6/ STAT1 |
| HIV_UP | hsa04540 | 4.188 | 0.04656726811 | GNAI2/ MAP2K5/ PRKACB/ TUBA1B/ TUBB |
| HIV_UP | GO:0002285 | 3.230681969 | 0.04857301155 | CD74/ CORO1A/ GATA3/ HLA-DRA/ HLA-DRB1/ LGALS1/ NKG7/ OPA1/ TBX21 |
| HIV_UP | GO:0072567 | 11.64071122 | 0.04857301155 | CD74/ MAP2K5/ MCOLN2 |
| HIV_UP | GO:0039535 | 11.64071122 | 0.04857301155 | BIRC3/ DDX60/ NOP53 |
| INFECTION_UP | GO:0043368 | 33.90357143 | 0.0005170640287 | BRD4/ EP300/ NFATC3/ TOX/ ZAP70 |
| INFECTION_UP | GO:0046631 | 12.10841837 | 0.0008214007498 | BRD4/ CD81/ EP300/ HSPH1/ KMT2A/ TOX/ ZAP70 |
| INFECTION_UP | GO:0023023 | 32.67131648 | 0.001540817116 | CD81/ HLA-DRB5/ KLRC2/ TAPBP |
| INFECTION_UP | GO:0051251 | 8.072278912 | 0.001743057288 | BRD4/ CD81/ EP300/ HLA-DRB5/ HSPD1/ HSPH1/ TOX/ ZAP70 |
| INFECTION_UP | GO:0045059 | 67.80714286 | 0.002241410061 | NFATC3/ TOX/ ZAP70 |
| INFECTION_UP | GO:0001772 | 27.26775956 | 0.002708440045 | ARHGDIA/ CD37/ CD81/ ZAP70 |
| INFECTION_UP | GO:0002824 | 12.74570354 | 0.004021075276 | BRD4/ CD7/ CD81/ EP300/ HSPD1 |
| INFECTION_UP | GO:0002285 | 8.961296413 | 0.004084692464 | BRD4/ CD81/ EP300/ HSPD1/ KLRC2/ KMT2A |
| INFECTION_UP | GO:0043373 | 32.80990783 | 0.007124159771 | BRD4/ EP300/ TOX |
| INFECTION_UP | GO:0140033 | 37.77620968 | 0.007741695777 | BRD4/ EP300/ KMT2A |
| INFECTION_UP | GO:0045345 | 113.0119048 | 0.00803045702 | HSPH1/ NLRC5 |
| INFECTION_UP | GO:0044183 | 16.11784946 | 0.008604547744 | HSPD1/ HSPH1/ TAPBP/ TCP1 |

| | | | | |
|---|---|---|---|---|
| INFECTION_UP | GO:2000330 | 84.75892857 | 0.01110965429 | BRD4/ EP300 |
| INFECTION_UP | GO:0023024 | 86.34562212 | 0.0130382946 | KLRC2/ TAPBP |
| INFECTION_UP | GO:0034504 | 6.337116155 | 0.01394864872 | ADAR/ EP300/ MARK3/ NFATC3/ TCP1/ ZBTB7A |
| INFECTION_UP | GO:2000516 | 21.18973214 | 0.01462058337 | BRD4/ CD81/ EP300 |
| INFECTION_UP | GO:0002429 | 6.054209184 | 0.01643980037 | CD81/ DGKZ/ EP300/ KLRC2/ MYO1G/ ZAP70 |
| INFECTION_UP | GO:0046638 | 18.16262755 | 0.01990195123 | BRD4/ EP300/ ZAP70 |
| INFECTION_UP | GO:0002708 | 9.971638655 | 0.0214166902 | CD7/ CD81/ HSPD1/ KLRC2 |
| INFECTION_UP | GO:0002456 | 9.827122153 | 0.02150463736 | CD7/ CD81/ HSPD1/ MYO1G |
| INFECTION_UP | GO:0051293 | 16.67388759 | 0.02163749347 | DYNC1H1/ LLGL2/ MAP4 |
| INFECTION_UP | GO:0051015 | 7.161366763 | 0.02473857283 | ABLIM1/ ADD1/ MYO1G/ MYO9B/ RCSD1 |
| INFECTION_UP | GO:0031490 | 9.444052419 | 0.02473857283 | EP300/ H1-10/ H1-3/ TOX |
| INFECTION_UP | GO:0045159 | 46.49379653 | 0.02473857283 | LLGL2/ NHERF1 |
| INFECTION_UP | GO:0097157 | 43.17281106 | 0.02473857283 | EP300/ PRPF8 |
| INFECTION_UP | GO:1900034 | 42.37946429 | 0.02684941089 | CHORDC1/ EP300 |
| INFECTION_UP | GO:0048002 | 14.1264881 | 0.03139273243 | ERAP2/ HLA-DRB5/ TAPBP |
| INFECTION_UP | GO:0042113 | 5.968938632 | 0.03468566331 | CD81/ EP300/ HSPD1/ ZAP70/ ZBTB7A |
| INFECTION_UP | GO:0061077 | 13.38298872 | 0.03468566331 | CHORDC1/ HSPH1/ TCP1 |
| INFECTION_UP | GO:0060216 | 33.90357143 | 0.0355986628 | ADAR/ KMT2A |
| INFECTION_UP | GO:0002039 | 13.13955119 | 0.03605416189 | BRD4/ EP300/ HSPD1 |
| INFECTION_UP | GO:0007009 | 7.838976053 | 0.03696145409 | AHNAK/ MTSS1/ NHERF1/ VMP1 |
| INFECTION_UP | GO:0002396 | 32.28911565 | 0.03696145409 | HLA-DRB5/ TAPBP |
| INFECTION_UP | GO:0016032 | 4.500474083 | 0.03842080852 | ADAR/ BRD4/ CD81/ EP300/ SELPLG/ ST6GAL1 |
| INFECTION_UP | GO:0032943 | 5.557962529 | 0.03842080852 | CD81/ HSPD1/ SH2D2A/ ST6GAL1/ ZAP70 |
| INFECTION_UP | GO:0019885 | 29.48136646 | 0.03842080852 | ERAP2/ TAPBP |
| INFECTION_UP | GO:1990858 | 29.48136646 | 0.03842080852 | EP300/ KLRC2 |
| INFECTION_UP | GO:0060339 | 28.25297619 | 0.04005628922 | ADAR/ NLRC5 |
| INFECTION_UP | GO:0046641 | 28.25297619 | 0.04005628922 | CD81/ ZAP70 |
| INFECTION_UP | GO:0045088 | 4.328115502 | 0.04214379743 | ABHD17A/ ADAR/ EP300/ HSPD1/ KLRC2/ NLRC5 |
| INFECTION_UP | GO:0044403 | 5.264529725 | 0.04214379743 | BRD4/ CD81/ EP300/ HSPD1/ SELPLG |
| INFECTION_UP | GO:0005096 | 5.555324953 | 0.0433789474 | ARHGAP10/ ARHGAP30/ ARHGDIA/ LLGL2/ MYO9B |
| INFECTION_UP | GO:0002520 | 6.713578501 | 0.04602710157 | ADAR/ HSPD1/ KMT2A/ TOX |
| INFECTION_UP | GO:0007097 | 24.21683673 | 0.04602710157 | DYNC1H1/ NHERF1 |
| INFECTION_UP | GO:0002726 | 24.21683673 | 0.04602710157 | CD7/ CD81 |
| INFECTION_UP | GO:0060765 | 23.3817734 | 0.04834642161 | EP300/ ZBTB7A |
| SYPH_PEAK | GO:0030098 | 6.210305549 | 5.824295456e-06 | BCL11B/ BCL2/ CARD11/ EOMES/ FGL2/ FOSL2/ IKZF1/ IKZF3/ INPP5D/ JUNB/ LCK/ PTK2B/ RUNX1/ ST3GAL1/ STAT3/ TGFB1/ TMEM131L |
| SYPH_PEAK | R-HSA-3371568 | 47.38945578 | 3.002924645e-05 | CREBBP/ DNAJB1/ HSP90AA1/ HSP90AB1/ HSPA8 |
| SYPH_PEAK | R-HSA-9012999 | 4.728391134 | 6.916865958e-05 | ARHGAP25/ ARHGEF1/ ARHGEF3/ FGD3/ FMNL1/ GRB2/ HSP90AA1/ HSP90AB1/ HSPE1/ LCK/ MSI2/ PTK2B/ RAPGEF1/ SPTBN1/ STK10/ TRIO |
| SYPH_PEAK | R-HSA-9725370 | 11.66509681 | 6.96257813e-05 | EML4/ GRB2/ JUN/ JUNB/ MCL1/ MYH9/ SQSTM1/ STAT3 |

| | | | | |
|---|---|---|---|---|
| SYPH_PEAK | R-HSA-5663202 | 4.289562808 | 0.0001718687218 | CREBBP/ EML4/ GRB2/ HSP90AA1/ JUN/ JUNB/ LCK/ MCL1/ MYH9/ SND1/ SPTBN1/ SQSTM1/ STAT3/ TGFB1/ ZC3HAV1 |
| SYPH_PEAK | R-HSA-6785807 | 9.828924162 | 0.0001718687218 | BCL2/ HSP90AA1/ HSPA8/ JUNB/ MCL1/ STAT3/ TGFB1/ TNFRSF1B |
| SYPH_PEAK | R-HSA-449147 | 4.20794322 | 0.0001909317386 | BCL2/ GRB2/ HSP90AA1/ HSPA8/ INPP5D/ JUN/ JUNB/ LCK/ MCL1/ PTK2B/ RAPGEF1/ SQSTM1/ STAT3/ TGFB1/ TNFRSF1B |
| SYPH_PEAK | GO:0007264 | 4.7542653 | 0.0005907582218 | ARHGAP25/ ARHGEF1/ ARHGEF3/ AUTS2/ CCDC88C/ CYTH1/ FGD3/ GRAP2/ GRB2/ IQSEC1/ RAP1GAP2/ RAPGEF1/ RIPOR2/ SQSTM1/ TRIO |
| SYPH_PEAK | GO:0030099 | 4.830867653 | 0.0008944780707 | FLI1/ FLNA/ FOSL2/ IKZF1/ INPP5D/ JUNB/ KLF13/ MYH9/ NEDD9/ PTK2B/ RUNX1/ STAT3/ TGFB1/ ZFP36 |
| SYPH_PEAK | GO:0002429 | 5.557962529 | 0.001026606814 | BCL2/ CARD11/ CREBBP/ EZR/ FGR/ FOSL2/ GRB2/ INPP5D/ LCK/ SLA2/ SPPL3/ SQSTM1 |
| SYPH_PEAK | GO:0005938 | 5.537452711 | 0.001510425678 | EZR/ FLNA/ FMNL1/ GRB2/ MYH9/ NEDD9/ PTK2B/ SEPTIN6/ SPTBN1/ STIM1/ STK39 |
| SYPH_PEAK | R-HSA-451927 | 15.0784632 | 0.001515755352 | GRB2/ INPP5D/ LCK/ PTK2B/ STAT3 |
| SYPH_PEAK | GO:0001772 | 17.04234973 | 0.001624282047 | CARD11/ EZR/ LCK/ MYH9/ NEDD9 |
| SYPH_PEAK | GO:1903706 | 4.672282588 | 0.001683109612 | CARD11/ FGL2/ IKZF3/ INPP5D/ JUNB/ KLF13/ NEDD9/ PTK2B/ RUNX1/ STAT3/ TGFB1/ TMEM131L/ ZFP36 |
| SYPH_PEAK | GO:0031647 | 4.825517855 | 0.002425149722 | ATP1B3/ BCL2/ CCDC88C/ CREBBP/ FLNA/ HSP90AA1/ HSP90AB1/ HSPA8/ MYLIP/ RNF149/ TMC8/ USP36 |
| SYPH_PEAK | GO:0051169 | 5.034860174 | 0.002638091313 | ATXN1/ FLNA/ HSP90AA1/ HSPA8/ NEAT1/ POM121/ POM121C/ SMG6/ SQSTM1/ STAT3/ TGFB1 |
| SYPH_PEAK | GO:0070661 | 5.020095188 | 0.002638091313 | BCL2/ CARD11/ CCND3/ FOSL2/ IKZF3/ INPP5D/ JUNB/ RIPOR2/ TGFB1/ TMEM131L/ TNFRSF1B |
| SYPH_PEAK | hsa04380 | 6.635915493 | 0.003640523813 | FOSB/ FOSL2/ GRB2/ JUN/ JUNB/ LCK/ SQSTM1/ TGFB1 |
| SYPH_PEAK | hsa04659 | 7.564334862 | 0.003640523813 | HSP90AA1/ HSP90AB1/ JUN/ LCK/ RUNX1/ STAT3/ TGFB1 |
| SYPH_PEAK | R-HSA-3371497 | 12.06277056 | 0.003722895616 | DNAJB1/ HSP90AA1/ HSP90AB1/ HSPA8/ TUBA4A |
| SYPH_PEAK | GO:0046632 | 8.190681622 | 0.004147799618 | BCL11B/ BCL2/ EOMES/ FOSL2/ JUNB/ RUNX1/ STAT3 |
| SYPH_PEAK | GO:0030183 | 7.83712702 | 0.004905059979 | BCL2/ CARD11/ FOSL2/ IKZF3/ INPP5D/ PTK2B/ ST3GAL1 |
| SYPH_PEAK | GO:0008360 | 7.781147541 | 0.004905059979 | ATP10A/ EZR/ FGD3/ FGR/ FMNL1/ MYH9/ PTK2B |
| SYPH_PEAK | R-HSA-9680350 | 17.12135177 | 0.004947172417 | GRAP2/ GRB2/ INPP5D/ STAT3 |
| SYPH_PEAK | GO:0050852 | 7.311145341 | 0.006638426665 | CARD11/ EZR/ FOSL2/ INPP5D/ LCK/ SLA2/ SPPL3 |
| SYPH_PEAK | GO:0001931 | 35.05854801 | 0.00715735393 | DNM2/ EZR/ MYH9 |
| SYPH_PEAK | R-HSA-9018519 | 6.233780761 | 0.007168498634 | BCL2/ CREBBP/ FOSB/ HSP90AA1/ HSP90AB1/ JUN/ RUNX1 |
| SYPH_PEAK | GO:0048872 | 4.590647517 | 0.007774949402 | BCL2/ CARD11/ FOSL2/ IKZF1/ INPP5D/ KLF13/ PACS1/ STAT3/ TGFB1/ ZFP36 |
| SYPH_PEAK | GO:0048545 | 4.577145612 | 0.007774949402 | BCL2/ FOSB/ FOSL2/ HSPA8/ JUN/ JUNB/ STK39/ TGFB1/ TRERF1/ ZFP36 |
| SYPH_PEAK | GO:0032623 | 11.78961749 | 0.007774949402 | CARD11/ EZR/ RUNX1/ SPTBN1/ ZFP36 |
| SYPH_PEAK | GO:1903829 | 3.757495794 | 0.008481055251 | ACSL3/ CREBBP/ EZR/ FLNA/ HSP90AA1/ HSP90AB1/ HUWE1/ NEDD9/ SPTBN1/ SQSTM1/ TGFB1/ USP36 |

| | | | | |
|---|---|---|---|---|
| SYPH_PEAK | GO:0045637 | 5.684856651 | 0.008481055251 | INPP5D/ KLF13/ NEDD9/ PTK2B/ RUNX1/ STAT3/ TGFB1/ ZFP36 |
| SYPH_PEAK | GO:2000401 | 11.11592506 | 0.008481055251 | NEDD9/ PTK2B/ RIPOR2/ STK10/ STK39 |
| SYPH_PEAK | GO:0045619 | 5.582886128 | 0.009159174267 | CARD11/ FGL2/ IKZF3/ INPP5D/ JUNB/ RUNX1/ TGFB1/ TMEM131L |
| SYPH_PEAK | R-HSA-9607240 | 13.96741855 | 0.009556635195 | GRAP2/ GRB2/ LCK/ SLA2 |
| SYPH_PEAK | GO:0051250 | 6.44592104 | 0.009813247287 | FGL2/ FGR/ INPP5D/ RIPOR2/ RUNX1/ TGFB1/ TMEM131L |
| SYPH_PEAK | GO:0043374 | 31.12459016 | 0.009813247287 | BCL2/ EOMES/ RUNX1 |
| SYPH_PEAK | R-HSA-187037 | 6.922981366 | 0.01025601314 | DNM2/ FOSB/ GRB2/ JUNB/ RAPGEF1/ STAT3 |
| SYPH_PEAK | R-HSA-9820965 | 24.87946429 | 0.01025601314 | HSP90AA1/ HSP90AB1/ HSPA8 |
| SYPH_PEAK | GO:0072659 | 4.747818839 | 0.01052011907 | ACSL3/ ATP1B3/ EZR/ FLNA/ NSF/ PACS1/ SPTBN1/ SQSTM1/ TGFB1 |
| SYPH_PEAK | R-HSA-204998 | 8.729636591 | 0.01130464539 | ARHGEF1/ ARHGEF3/ FGD3/ SQSTM1/ TRIO |
| SYPH_PEAK | R-HSA-9820952 | 6.63452381 | 0.01144074282 | ARIH1/ CREBBP/ HSP90AA1/ HSP90AB1/ HSPA8/ MED15 |
| SYPH_PEAK | R-HSA-1433557 | 12.34330011 | 0.01144074282 | GRAP2/ GRB2/ LCK/ STAT3 |
| SYPH_PEAK | GO:0030695 | 3.753530767 | 0.01202584254 | ARHGAP25/ ARHGEF1/ ARHGEF3/ CCDC88C/ CYTH1/ FGD3/ GRB2/ IQSEC1/ JUN/ RAP1GAP2/ RAPGEF1/ TRIO |
| SYPH_PEAK | GO:0031625 | 4.592151844 | 0.01202584254 | ARIH1/ BCL2/ HSP90AA1/ HSP90AB1/ HSPA8/ JUN/ PTK2B/ SQSTM1/ TNFRSF1B |
| SYPH_PEAK | GO:0005085 | 5.341981468 | 0.01202584254 | ARHGEF1/ ARHGEF3/ CCDC88C/ CYTH1/ FGD3/ IQSEC1/ RAPGEF1/ TRIO |
| SYPH_PEAK | GO:0044183 | 10.23879781 | 0.01202584254 | DNAJB1/ HSP90AA1/ HSP90AB1/ HSPA8/ HSPE1 |
| SYPH_PEAK | GO:0001784 | 13.07080572 | 0.0138567909 | FGR/ GRAP2/ GRB2/ LCK |
| SYPH_PEAK | GO:0045296 | 4.138436242 | 0.01442247382 | DNAJB1/ EZR/ FLNA/ HSP90AB1/ HSPA8/ MYH9/ SND1/ SPTBN1/ ZC3HAV1 |
| SYPH_PEAK | GO:0070182 | 21.94028103 | 0.01442247382 | HSP90AA1/ HSP90AB1/ SMG6 |
| SYPH_PEAK | GO:0140313 | 11.37644202 | 0.01457253145 | SEC14L1/ SQSTM1/ STAT3/ TMC8 |
| SYPH_PEAK | GO:1905172 | 61.43278689 | 0.01457253145 | NEAT1/ SND1 |
| SYPH_PEAK | GO:0032570 | 14.14754098 | 0.01459275892 | FOSB/ HSPA8/ TGFB1/ TRERF1 |
| SYPH_PEAK | GO:0072594 | 3.705308353 | 0.01468327076 | FLNA/ HSP90AA1/ HSPA8/ HUWE1/ POM121/ POM121C/ SQSTM1/ STAT3/ TGFB1/ USP36/ VPS37B |
| SYPH_PEAK | GO:1903729 | 25.93715847 | 0.01468327076 | ATP10A/ DNM2/ MYH9 |
| SYPH_PEAK | GO:0007015 | 3.681403138 | 0.01500056045 | ARHGAP25/ BCL2/ CYRIA/ DNM2/ EZR/ FLNA/ GRB2/ MYO1F/ NEDD9/ PTK2B/ SPTBN1 |
| SYPH_PEAK | GO:0045646 | 13.53243051 | 0.01525080273 | INPP5D/ KLF13/ STAT3/ ZFP36 |
| SYPH_PEAK | R-HSA-2029480 | 7.714562569 | 0.01552642605 | FGR/ GRB2/ HSP90AA1/ HSP90AB1/ MYH9 |
| SYPH_PEAK | GO:0150034 | 4.657837932 | 0.01558595995 | AUTS2/ DNM2/ FLNA/ HSP90AA1/ HSP90AB1/ HSPA8/ PTK2B/ SEPTIN6 |
| SYPH_PEAK | GO:0032783 | 65.44262295 | 0.01558595995 | AFF1/ AFF4 |
| SYPH_PEAK | GO:0035976 | 65.44262295 | 0.01558595995 | JUN/ JUNB |
| SYPH_PEAK | GO:0034504 | 4.363260303 | 0.01574034961 | CREBBP/ FLNA/ HSP90AA1/ HSPA8/ POM121/ POM121C/ SQSTM1/ STAT3/ TGFB1 |
| SYPH_PEAK | GO:0140131 | 24.57204487 | 0.01574034961 | NEDD9/ PTK2B/ STK39 |
| SYPH_PEAK | GO:0140545 | 51.19398907 | 0.01574943628 | HSPA8/ NSF |
| SYPH_PEAK | GO:0051400 | 51.19398907 | 0.01574943628 | BCL2/ MCL1 |
| SYPH_PEAK | GO:0007492 | 8.943847748 | 0.01618525092 | EOMES/ GRB2/ MYH9/ SSBP3/ TGFB1 |

| | | | | |
|---|---|---|---|---|
| SYPH_PEAK | GO:0097718 | 16.45521077 | 0.01646643449 | EZR/ HSP90AA1/ HSP90AB1 |
| SYPH_PEAK | GO:0030235 | 43.88056206 | 0.01646643449 | HSP90AA1/ HSP90AB1 |
| SYPH_PEAK | GO:0043120 | 43.88056206 | 0.01646643449 | TMC8/ TNFRSF1B |
| SYPH_PEAK | GO:0050856 | 12.70391435 | 0.01748242188 | CARD11/ EZR/ LCK/ SLA2 |
| SYPH_PEAK | GO:0042470 | 7.303864169 | 0.01856695109 | ATP1B3/ HSP90AA1/ HSP90AB1/ HSPA8/ SND1 |
| SYPH_PEAK | R-HSA-9013418 | 17.30745342 | 0.01925556644 | HSP90AA1/ HSP90AB1/ MSI2 |
| SYPH_PEAK | GO:0044295 | 16.36065574 | 0.01991101732 | FLNA/ HSP90AA1/ HSP90AB1 |
| SYPH_PEAK | R-HSA-9662361 | 9.828924162 | 0.02011907566 | EZR/ MYH9/ RIPOR2/ SPTBN1 |
| SYPH_PEAK | GO:0001228 | 3.199624317 | 0.02017454739 | BCL11B/ FLI1/ FOSB/ FOSL2/ IKZF3/ JUN/ JUNB/ KLF13/ RUNX1/ STAT3 |
| SYPH_PEAK | hsa05211 | 8.413392857 | 0.02047792964 | CREBBP/ GRB2/ JUN/ RAPGEF1/ TGFB1 |
| SYPH_PEAK | GO:1904813 | 6.597038604 | 0.0215594847 | FGL2/ HSP90AA1/ HSP90AB1/ HSPA8/ HUWE1 |
| SYPH_PEAK | GO:0097413 | 40.90163934 | 0.0215594847 | ARIH1/ SQSTM1 |
| SYPH_PEAK | R-HSA-1169410 | 5.343240652 | 0.02177658509 | ARIH1/ FLNA/ HSPA8/ POM121/ STAT3/ TUBA4A |
| SYPH_PEAK | R-HSA-9679191 | 6.701539202 | 0.02177658509 | ATP1B3/ GATAD2B/ GRB2/ HSP90AA1/ HSP90AB1 |
| SYPH_PEAK | GO:0016234 | 8.843597696 | 0.02183128673 | ARIH1/ ATXN1/ FGR/ SQSTM1 |
| SYPH_PEAK | hsa05161 | 5.058358896 | 0.02254401876 | BCL2/ CREBBP/ GRB2/ JUN/ PTK2B/ STAT3/ TGFB1 |
| SYPH_PEAK | GO:1904685 | 62.24918033 | 0.02262258096 | MBP/ STAT3 |
| SYPH_PEAK | GO:1903525 | 62.24918033 | 0.02262258096 | ATP10A/ DNM2 |
| SYPH_PEAK | GO:0031490 | 5.999295594 | 0.02296898184 | CREBBP/ EOMES/ GATAD2B/ KDM6A/ STAT3 |
| SYPH_PEAK | GO:0008180 | 13.63387978 | 0.02406633061 | GRB2/ HSP90AB1/ MYH9 |
| SYPH_PEAK | R-HSA-9679506 | 3.205083966 | 0.02434773358 | ATP1B3/ CREBBP/ GATAD2B/ GRB2/ HSP90AA1/ HSP90AB1/ PARP14/ POM121/ RUNX1/ ST3GAL1 |
| SYPH_PEAK | GO:0045638 | 7.859744991 | 0.02446534209 | INPP5D/ KLF13/ PTK2B/ RUNX1/ ZFP36 |
| SYPH_PEAK | R-HSA-8939246 | 37.91156463 | 0.02482722328 | CREBBP/ RUNX1 |
| SYPH_PEAK | R-HSA-9612973 | 5.038878843 | 0.02529905873 | HSP90AA1/ HSP90AB1/ HSPA8/ SQSTM1/ TUBA4A/ VPS37B |
| SYPH_PEAK | GO:0032535 | 3.879796558 | 0.0263724588 | CYRIA/ DNM2/ EZR/ GRB2/ HSP90AA1/ PTK2B/ RAP1GAP2/ SPTBN1/ STK39 |
| SYPH_PEAK | GO:0002064 | 4.97424957 | 0.0263724588 | BCL11B/ CCDC88C/ EZR/ FLNA/ FOSL2/ RAPGEF1/ TGFB1 |
| SYPH_PEAK | GO:0031960 | 5.985498108 | 0.0263724588 | BCL2/ FOSB/ FOSL2/ STK39/ TGFB1/ ZFP36 |
| SYPH_PEAK | GO:0070233 | 18.6747541 | 0.0263724588 | BCL11B/ BCL2/ ST3GAL1 |
| SYPH_PEAK | R-HSA-186763 | 13.72660099 | 0.0267051518 | GRB2/ RAPGEF1/ STAT3 |
| SYPH_PEAK | R-HSA-9843745 | 6.086719091 | 0.0269811573 | CCND3/ CREBBP/ GATAD2B/ MED15/ TGFB1 |
| SYPH_PEAK | GO:0071211 | 51.87431694 | 0.02770121655 | HSPA8/ SQSTM1 |
| SYPH_PEAK | GO:2000634 | 51.87431694 | 0.02770121655 | STAT3/ TGFB1 |
| SYPH_PEAK | GO:0045639 | 7.272100506 | 0.02846674495 | INPP5D/ NEDD9/ RUNX1/ STAT3/ TGFB1 |
| SYPH_PEAK | GO:0003714 | 4.495081967 | 0.02860234635 | CREBBP/ DNAJB1/ JAZF1/ PARP14/ PHF12/ TRERF1 |
| SYPH_PEAK | GO:0051117 | 7.31342701 | 0.02860234635 | ATP1B3/ DNAJB1/ EZR/ LCK |
| SYPH_PEAK | GO:0002285 | 4.79894562 | 0.02936487305 | EOMES/ FGL2/ JUNB/ PTK2B/ ST3GAL1/ STAT3/ TGFB1 |
| SYPH_PEAK | GO:0010586 | 7.073770492 | 0.03041281854 | JUN/ NEAT1/ SND1/ STAT3/ TGFB1 |
| SYPH_PEAK | GO:0045058 | 9.880822274 | 0.03041281854 | BCL11B/ BCL2/ CARD11/ STAT3 |

| | | | | |
|---|---|---|---|---|
| SYPH_PEAK | GO:0007229 | 7.01004283 | 0.03089773213 | FGR/ FLNA/ MYH9/ NEDD9/ PTK2B |
| SYPH_PEAK | GO:0033619 | 9.726434426 | 0.03093562208 | MYH9/ SPPL3/ TGFB1/ TNFRSF1B |
| SYPH_PEAK | GO:0051015 | 4.367259731 | 0.03096112526 | EZR/ FLNA/ FMNL1/ MYH9/ MYO1F/ SPTBN1 |
| SYPH_PEAK | GO:0019903 | 5.332707195 | 0.03096112526 | BCL2/ GRB2/ HSP90AA1/ LCK/ STAT3 |
| SYPH_PEAK | GO:0030316 | 6.947453162 | 0.03118247439 | FOSL2/ INPP5D/ JUNB/ NEDD9/ TGFB1 |
| SYPH_PEAK | R-HSA-2871796 | 12.43973214 | 0.03125491395 | GRAP2/ GRB2/ JUN |
| SYPH_PEAK | GO:0043198 | 11.97121152 | 0.03144285615 | FLNA/ HSPA8/ NSF |
| SYPH_PEAK | GO:1903078 | 9.576796974 | 0.03148986981 | ACSL3/ EZR/ SPTBN1/ SQSTM1 |
| SYPH_PEAK | GO:0040014 | 9.576796974 | 0.03148986981 | BCL2/ EZR/ FOSL2/ STAT3 |
| SYPH_PEAK | R-HSA-111453 | 29.48677249 | 0.03184339775 | BCL2/ STAT3 |
| SYPH_PEAK | R-HSA-8951936 | 29.48677249 | 0.03184339775 | RUNX1/ TGFB1 |
| SYPH_PEAK | R-HSA-373760 | 5.575230092 | 0.03208900225 | DNM2/ EZR/ HSPA8/ SPTBN1/ TUBA4A |
| SYPH_PEAK | GO:0045088 | 3.311126613 | 0.03326071204 | CREBBP/ FGR/ HSP90AA1/ HSPA8/ PARP14/ SEC14L1/ SQSTM1/ TGFB1/ ZC3HAV1/ ZNFX1 |
| SYPH_PEAK | GO:0051168 | 5.397327774 | 0.03326071204 | ATXN1/ NEAT1/ POM121/ POM121C/ SMG6/ TGFB1 |
| SYPH_PEAK | GO:0030888 | 9.290922437 | 0.03326071204 | BCL2/ CARD11/ IKZF3/ INPP5D |
| SYPH_PEAK | hsa04517 | 3.533625 | 0.03423722316 | EZR/ GRB2/ INPP5D/ LCK/ MBP/ MYH9/ SPTBN1/ TRIO/ TUBA4A |
| SYPH_PEAK | hsa05132 | 3.784337349 | 0.03423722316 | BCL2/ CYTH1/ DNM2/ FLNA/ HSP90AA1/ HSP90AB1/ JUN/ TUBA4A |
| SYPH_PEAK | hsa04915 | 5.084352518 | 0.03423722316 | BCL2/ GRB2/ HSP90AA1/ HSP90AB1/ HSPA8/ JUN |
| SYPH_PEAK | GO:0050863 | 3.56388437 | 0.03480058758 | CARD11/ FGL2/ JUNB/ LCK/ RIPOR2/ RUNX1/ TGFB1/ TMEM131L/ TNFRSF1B |
| SYPH_PEAK | GO:0004715 | 10.23879781 | 0.03480543921 | FGR/ LCK/ PTK2B |
| SYPH_PEAK | GO:0001223 | 10.23879781 | 0.03480543921 | CREBBP/ RUNX1/ TRERF1 |
| SYPH_PEAK | GO:0060964 | 15.06028556 | 0.03533424911 | STAT3/ TGFB1/ ZFP36 |
| SYPH_PEAK | GO:0051085 | 14.58965164 | 0.03543305514 | DNAJB1/ HSPA8/ HSPE1 |
| SYPH_PEAK | GO:0045647 | 38.9057377 | 0.03543305514 | KLF13/ ZFP36 |
| SYPH_PEAK | R-HSA-9833482 | 7.076825397 | 0.03715942921 | ARIH1/ HSPA8/ STAT3/ TUBA4A |
| SYPH_PEAK | GO:0045862 | 3.830718789 | 0.03723112191 | HSPA8/ HSPE1/ LCK/ MBP/ MYH9/ PTK2B/ STAT3/ TNFRSF1B |
| SYPH_PEAK | GO:0048863 | 4.340082294 | 0.03723112191 | BCL2/ CCDC88C/ EOMES/ GATAD2B/ MSI2/ STAT3/ ZFP36 |
| SYPH_PEAK | hsa05130 | 4.102052239 | 0.03873972989 | ARHGEF1/ CYTH1/ EZR/ JUN/ MYH9/ MYO1F/ TUBA4A |
| SYPH_PEAK | GO:0035254 | 9.598872951 | 0.03982145476 | NSF/ PTK2B/ SQSTM1 |
| SYPH_PEAK | GO:0048771 | 4.888679083 | 0.04166990223 | FLNA/ FOSL2/ INPP5D/ PTK2B/ RUNX1/ TGFB1 |
| SYPH_PEAK | GO:0032743 | 13.33911007 | 0.04166990223 | CARD11/ RUNX1/ SPTBN1 |
| SYPH_PEAK | GO:0060613 | 34.58287796 | 0.04166990223 | ARRDC3/ FOSL2 |
| SYPH_PEAK | GO:0048382 | 34.58287796 | 0.04166990223 | EOMES/ SSBP3 |
| SYPH_PEAK | R-HSA-9634285 | 24.12554113 | 0.041690312 | GRB2/ HSP90AA1 |
| SYPH_PEAK | R-HSA-8875555 | 24.12554113 | 0.041690312 | GRB2/ RAPGEF1 |
| SYPH_PEAK | R-HSA-9818030 | 24.12554113 | 0.041690312 | BCL2/ CREBBP |
| SYPH_PEAK | R-HSA-9031628 | 10.20695971 | 0.04218047822 | DNM2/ FOSB/ JUNB |
| SYPH_PEAK | R-HSA-389359 | 22.11507937 | 0.04243897393 | GRB2/ LCK |
| SYPH_PEAK | R-HSA-9020558 | 22.11507937 | 0.04243897393 | LCK/ PTK2B |

| | | | | |
|---|---|---|---|---|
| SYPH_PEAK | R-HSA-9673770 | 22.11507937 | 0.04243897393 | GRB2/ STAT3 |
| SYPH_PEAK | R-HSA-202733 | 4.842718109 | 0.04313663135 | ATP1B3/ GRB2/ INPP5D/ LCK/ TGFB1 |
| SYPH_PEAK | hsa05417 | 3.8171875 | 0.04356428698 | ARHGEF1/ BCL2/ HSP90AA1/ HSP90AB1/ HSPA8/ JUN/ STAT3 |
| SYPH_PEAK | hsa05207 | 3.799596774 | 0.04356428698 | BCL2/ CCND3/ GRB2/ HSP90AA1/ HSP90AB1/ JUN/ STAT3 |
| SYPH_PEAK | hsa05215 | 5.556014151 | 0.04356428698 | BCL2/ CREBBP/ GRB2/ HSP90AA1/ HSP90AB1 |
| SYPH_PEAK | GO:0043209 | 10.22540984 | 0.04474742447 | BCL2/ HSP90AA1/ MBP |
| SYPH_PEAK | GO:0010038 | 3.608648135 | 0.04680491793 | BCL2/ FOSB/ HSPA8/ JUNB/ MT2A/ PTK2B/ STIM1/ STK39 |
| SYPH_PEAK | GO:0006882 | 12.61807709 | 0.04680491793 | LCK/ MT2A/ TMC8 |
| SYPH_PEAK | GO:0030853 | 31.12459016 | 0.04712135705 | INPP5D/ RUNX1 |
| SYPH_PEAK | GO:0051028 | 5.763812993 | 0.04715325094 | NEAT1/ POM121/ POM121C/ SMG6/ ZFP36 |
| SYPH_PEAK | GO:0007159 | 3.257224552 | 0.04770906479 | CARD11/ EZR/ FGL2/ LCK/ RIPOR2/ RUNX1/ STK10/ TGFB1/ TMEM131L |
| SYPH_PEAK | GO:0002028 | 7.591363455 | 0.04922112649 | ATP1B3/ MLLT6/ STK39/ TGFB1 |
| SYPH_PEAK | hsa04630 | 4.206696429 | 0.04976731418 | BCL2/ CCND3/ CREBBP/ GRB2/ MCL1/ STAT3 |
| HIV_DOWN | R-HSA-156827 | 15.11663653 | 6.623409905e-09 | EIF1AX/ EIF2S3/ RPL23A/ RPL26/ RPL35A/ RPL7/ RPL9/ RPS24/ RPS27/ RPS27A/ RPS4X/ RPSA |
| HIV_DOWN | R-HSA-72695 | 18.99269718 | 2.491858576e-06 | EIF1AX/ EIF2S3/ RPS24/ RPS27/ RPS27A/ RPS4X/ RPSA |
| HIV_DOWN | hsa05171 | 7.118723849 | 5.024718444e-06 | FOS/ IFNAR2/ MAPK1/ RPL23A/ RPL26/ RPL35A/ RPL7/ RPL9/ RPS24/ RPS27/ RPS27A/ RPS4X/ RPSA |
| HIV_DOWN | GO:0022627 | 25.68854569 | 6.70057241e-06 | DDX3X/ RPS24/ RPS27/ RPS27A/ RPS4X/ RPSA |
| HIV_DOWN | R-HSA-9692914 | 11.75738397 | 9.541135896e-06 | ITCH/ PPIA/ RPS24/ RPS27/ RPS27A/ RPS4X/ RPSA/ SMAD3 |
| HIV_DOWN | GO:0046631 | 9.958325386 | 1.790904707e-05 | CD3E/ CD69/ CD83/ FOXP1/ IL18R1/ ITCH/ ITK/ KLRF1/ RORA/ RUNX3/ ZBTB16 |
| HIV_DOWN | GO:0030098 | 5.831336931 | 0.0001029219768 | B2M/ CD3E/ CD69/ CD83/ FOXP1/ ID2/ IL18R1/ ITK/ KLF6/ PPP2R3C/ RORA/ RUNX3/ ZBTB16/ ZFP36L2 |
| HIV_DOWN | hsa04659 | 9.605504587 | 0.0001100828909 | CD3E/ FOS/ HIF1A/ IFNGR1/ MAPK1/ NFATC2/ RORA/ SMAD3 |
| HIV_DOWN | GO:0022625 | 15.77366841 | 0.0006273570249 | RPL23A/ RPL26/ RPL35A/ RPL7/ RPL9 |
| HIV_DOWN | HALLMARK_TNFA_SIGNALING_VIA_NFKB | 4.548396226 | 0.0007080957878 | CD69/ CD83/ DUSP1/ FOS/ GADD45B/ ID2/ IER5/ KLF6/ PTPRE/ SMAD3/ ZBTB10 |
| HIV_DOWN | GO:0005925 | 4.665136835 | 0.0008187146924 | ARHGAP26/ B2M/ MAPK1/ PEAK1/ PHLDB2/ PPIA/ PXN/ RPL7/ RPL9/ RPS4X/ SSH2 |
| HIV_DOWN | hsa04658 | 8.443548387 | 0.002994908626 | CD3E/ FOS/ IFNGR1/ MAPK1/ NFATC2/ RUNX3 |
| HIV_DOWN | GO:0042254 | 5.597452755 | 0.003969316231 | DDX3X/ EIF1AX/ RPL26/ RPL35A/ RPL7/ RPS24/ RPS27/ RPS27A/ RPS4X/ RPSA |
| HIV_DOWN | GO:0045444 | 6.36236363 | 0.003969316231 | ID2/ LRRC8C/ PLCB1/ RGS2/ RORA/ SMAD3/ SYAP1/ ZBTB16/ ZFP36L2 |
| HIV_DOWN | hsa05321 | 9.914772727 | 0.004834448647 | IFNGR1/ IL18R1/ IL18RAP/ RORA/ SMAD3 |
| HIV_DOWN | GO:0043367 | 10.43760308 | 0.006865594546 | CD69/ CD83/ FOXP1/ IL18R1/ RORA/ RUNX3 |
| HIV_DOWN | GO:0035976 | 71.92792793 | 0.007063530714 | FOS/ NFATC2 |
| HIV_DOWN | GO:1903706 | 4.507694632 | 0.008491322714 | B2M/ CD69/ CD83/ FOS/ FOXP1/ HIF1A/ ID2/ PPP2R3C/ RUNX3/ ZBTB16/ ZFP36L2 |
| HIV_DOWN | R-HSA-449147 | 3.281130409 | 0.009256065107 | FOS/ HIF1A/ IL18R1/ IL18RAP/ IL32/ MAPK1/ PELI2/ PPIA/ RORA/ RPS27A/ SMAD3 |
| HIV_DOWN | R-HSA-8952158 | 56.43544304 | 0.009282086955 | RUNX3/ SMAD3 |

| | | | | |
|---|---|---|---|---|
| HIV_DOWN | GO:0001221 | 8.980110232 | 0.009329314708 | FOS/ HIF1A/ RORA/ RUNX3/ SMAD3/ ZBTB16 |
| HIV_DOWN | hsa04660 | 6.43647541 | 0.009532601891 | CD3E/ FOS/ ITK/ MAPK1/ NFATC2/ PPP2R3C |
| HIV_DOWN | R-HSA-9705683 | 4.8651244 | 0.01006018428 | B2M/ IFNAR2/ RPS24/ RPS27/ RPS27A/ RPS4X/ RPSA |
| HIV_DOWN | R-HSA-9006927 | 10.45100797 | 0.01006018428 | ELMO1/ HIF1A/ PXN/ RPS27A |
| HIV_DOWN | HALLMARK_ALLOGRAFT_REJECTION | 3.721415094 | 0.01021844875 | B2M/ CD3E/ GZMA/ HIF1A/ IFNAR2/ IFNGR1/ IL18RAP/ ITK/ RPL9 |
| HIV_DOWN | hsa04650 | 5.904135338 | 0.01324616236 | CD48/ IFNAR2/ IFNGR1/ KIR2DL3/ MAPK1/ NFATC2 |
| HIV_DOWN | hsa04062 | 4.746761658 | 0.01464547663 | CCL4L2/ CXCR4/ ELMO1/ ITK/ MAPK1/ PLCB1/ PXN |
| HIV_DOWN | hsa05167 | 4.674107143 | 0.01464547663 | FOS/ HIF1A/ IFNAR2/ IFNGR1/ MAPK1/ NFATC2/ RPS27A |
| HIV_DOWN | R-HSA-9635486 | 15.67651195 | 0.01518331271 | B2M/ MAPK1/ RPS27A |
| HIV_DOWN | GO:0046634 | 8.726520607 | 0.01539700752 | CD3E/ CD69/ CD83/ ITCH/ RUNX3/ ZBTB16 |
| HIV_DOWN | GO:0032040 | 9.719990261 | 0.01636488969 | RPS24/ RPS27/ RPS27A/ RPS4X |
| HIV_DOWN | R-HSA-9637628 | 40.31103074 | 0.01661862624 | B2M/ RPS27A |
| HIV_DOWN | GO:0062197 | 4.974932308 | 0.01666478237 | DDX3X/ FOS/ FOXP1/ HIF1A/ IL18RAP/ LRRC8C/ PPIA/ PRR5L/ PXN |
| HIV_DOWN | GO:0140375 | 6.82255128 | 0.0168960713 | CXCR4/ IFNAR2/ IFNGR1/ IL18R1/ IL18RAP/ KLRF1 |
| HIV_DOWN | GO:0001222 | 14.29486935 | 0.0168960713 | RORA/ RUNX3/ SMAD3/ ZBTB16 |
| HIV_DOWN | GO:0004904 | 70.04485981 | 0.01762440396 | IFNAR2/ IFNGR1 |
| HIV_DOWN | HALLMARK_IL2_STAT5_SIGNALING | 3.324547265 | 0.01822873413 | CD48/ CD83/ GADD45B/ IFNGR1/ IL18R1/ KLF6/ LRRC8C/ RORA |
| HIV_DOWN | HALLMARK_INFLAMMATORY_RESPONSE | 3.307924528 | 0.01822873413 | CD48/ CD69/ HIF1A/ IL18R1/ IL18RAP/ KLF6/ PTPRE/ RGS1 |
| HIV_DOWN | HALLMARK_HYPOXIA | 3.307924528 | 0.01822873413 | CITED2/ CXCR4/ DUSP1/ FOS/ GLRX/ KLF6/ PAM/ RORA |
| HIV_DOWN | R-HSA-2173795 | 14.10886076 | 0.01934026982 | MAPK1/ RPS27A/ SMAD3 |
| HIV_DOWN | hsa05170 | 4.280957944 | 0.02043709386 | B2M/ CD3E/ CXCR4/ FOS/ MAPK1/ NFATC2/ PXN |
| HIV_DOWN | hsa05142 | 6.35315534 | 0.02043709386 | CD3E/ FOS/ IFNGR1/ MAPK1/ PLCB1 |
| HIV_DOWN | R-HSA-9732724 | 35.2721519 | 0.02061790302 | IFNGR1/ MAPK1 |
| HIV_DOWN | R-HSA-9012546 | 35.2721519 | 0.02061790302 | IL18R1/ IL18RAP |
| HIV_DOWN | GO:0098586 | 9.968497322 | 0.02177671835 | DDX3X/ HIF1A/ IFNAR2/ IFNGR1/ SMAD3 |
| HIV_DOWN | GO:0070849 | 14.78660436 | 0.02177671835 | FOS/ MAPK1/ SYAP1/ ZFP36L2 |
| HIV_DOWN | R-HSA-400253 | 8.062206148 | 0.02245793894 | CREM/ HIF1A/ RORA/ RPS27A |
| HIV_DOWN | R-HSA-8949613 | 12.82623705 | 0.02346349255 | APOO/ ATP5MK/ ATP5PD |
| HIV_DOWN | hsa05208 | 4.035792952 | 0.0246134072 | ATP5PD/ COX6C/ FOS/ HIF1A/ MAPK1/ NDUFS5/ UQCRB |
| HIV_DOWN | hsa05163 | 4.035792952 | 0.0246134072 | B2M/ CCL4L2/ CXCR4/ MAPK1/ NFATC2/ PLCB1/ PXN |
| HIV_DOWN | R-HSA-1253288 | 31.35302391 | 0.02477146889 | ITCH/ RPS27A |
| HIV_DOWN | GO:0045598 | 7.242418463 | 0.02488172117 | ID2/ RORA/ SMAD3/ SYAP1/ ZBTB16/ ZFP36L2 |
| HIV_DOWN | R-HSA-450341 | 28.21772152 | 0.02961350136 | FOS/ MAPK1 |
| HIV_DOWN | R-HSA-198933 | 5.304082992 | 0.03340800497 | B2M/ CD3E/ KIR2DL3/ KLRF1/ SLAMF7 |
| HIV_DOWN | R-HSA-2454202 | 5.264500283 | 0.03340800497 | FOS/ ITK/ MAPK1/ NFATC2/ RPS27A |
| HIV_DOWN | R-HSA-2025928 | 25.65247411 | 0.03340800497 | NFATC2/ PPIA |
| HIV_DOWN | R-HSA-1234158 | 25.65247411 | 0.03340800497 | CITED2/ HIF1A |

| | | | | |
|---|---|---|---|---|
| HIV_DOWN | GO:0045619 | 5.569842002 | 0.03461287855 | CD69/ CD83/ ID2/ PPP2R3C/ RUNX3/ ZBTB16/ ZFP36L2 |
| HIV_DOWN | R-HSA-9692916 | 10.58164557 | 0.03514646769 | ITCH/ PPIA/ RPS27A |
| HIV_DOWN | GO:0042255 | 12.23718982 | 0.03558924657 | DDX3X/ EIF1AX/ RPS27/ RPSA |
| HIV_DOWN | GO:0002669 | 70.97570093 | 0.03558924657 | CD3E/ ITCH |
| HIV_DOWN | GO:0045926 | 5.353770545 | 0.03664666785 | CITED2/ DDX3X/ FOXP1/ HIF1A/ IFRD1/ RGS2/ SMAD3 |
| HIV_DOWN | GO:0019902 | 5.500905221 | 0.03779208142 | MAPK1/ NFATC2/ PXN/ SBF2/ SMAD3/ SYTL2 |
| HIV_DOWN | R-HSA-5675221 | 9.843391228 | 0.04247935262 | DUSP1/ MAPK1/ RPS27A |
| HIV_DOWN | GO:0001217 | 4.409298729 | 0.04479780625 | CREM/ FOXP1/ HIF1A/ NFATC2/ SMAD3/ ZBTB10/ ZBTB16 |
| HIV_DOWN | GO:0055022 | 19.01134846 | 0.04872819707 | CITED2/ FOXP1/ RGS2 |
| HIV_DOWN | GO:2000630 | 10.5933882 | 0.04905068222 | FOS/ HIF1A/ SMAD3/ XIST |
| INFECTION_DOWN | GO:0022626 | 56.38418079 | 4.782524326e-11 | RPL22/ RPL23/ RPL36A/ RPL37/ RPL37A/ RPL39/ RPS26/ RPS29 |
| INFECTION_DOWN | GO:0098556 | 277.2222222 | 0.0002072306767 | RPS26/ RPS29 |
| INFECTION_DOWN | GO:1990948 | 154.8512397 | 0.003730805215 | RPL23/ RPL37 |
| INFECTION_DOWN | GO:0016846 | 113.5575758 | 0.003730805215 | ALOX5AP/ GLO1 |
| INFECTION_DOWN | GO:0005925 | 7.84591195 | 0.009047863535 | RPL22/ RPL23/ RPL37A/ RPS29 |
| INFECTION_DOWN | R-HSA-8978868 | 9.499431818 | 0.01901048739 | AKR1C3/ ALOX5AP/ DBI |
| INFECTION_DOWN | R-HSA-2142753 | 18.89152542 | 0.02222733915 | AKR1C3/ ALOX5AP |
| INFECTION_DOWN | GO:0098803 | 15.12121212 | 0.03289069674 | ATP5ME/ NDUFA1 |
| INFECTION_DOWN | GO:0016655 | 29.36833856 | 0.037977129 | AKR1C3/ NDUFA1 |
| INFECTION_DOWN | R-HSA-1428517 | 6.430384615 | 0.04437057478 | ATP5ME/ GLO1/ NDUFA1 |
| SYPH_DROP | R-HSA-376176 | 8.482496195 | 0.001392992976 | CDC42/ FAU/ PSME1/ RPL27/ RPL38/ RPL41/ RPS21 |
| SYPH_DROP | R-HSA-1799339 | 14.09102402 | 0.001392992976 | FAU/ RPL27/ RPL38/ RPL41/ RPS21/ SEC11C |
| SYPH_DROP | hsa05208 | 8.588485493 | 0.002146735893 | ATP5F1E/ ATP5MC3/ NDUFA12/ NDUFC2/ NFKBIA/ UQCR11 |
| SYPH_DROP | HALLMARK_OXIDATIVE_PHOSPHORYLATION | 6.8484375 | 0.009797784554 | ACAA2/ ATP5F1E/ ATP5MC3/ NDUFC2/ UQCR11 |
| SYPH_DROP | GO:0005681 | 8.171620404 | 0.01216556726 | FRG1/ IVNS1ABP/ RBM17/ RBM3/ SNRPF |
| SYPH_DROP | GO:0098803 | 12.30200308 | 0.01216556726 | ATP5F1E/ ATP5MC3/ NDUFA12/ NDUFC2 |
| SYPH_DROP | GO:0015934 | 11.87035385 | 0.01216556726 | RBM3/ RPL27/ RPL38/ RPL41 |
| SYPH_DROP | R-HSA-9754678 | 15.61064426 | 0.0133664391 | FAU/ RPS21/ SNRPF |
| SYPH_DROP | hsa04932 | 8.278497694 | 0.01817331732 | CDC42/ NDUFA12/ NDUFC2/ UQCR11 |
| SYPH_DROP | GO:0031492 | 23.07512315 | 0.0215020454 | H3-3A/ HMGN1/ SMARCE1 |
| SYPH_DROP | R-HSA-163210 | 26.53809524 | 0.03436334524 | ATP5F1E/ ATP5MC3 |
| SYPH_DROP | R-HSA-611105 | 6.718505124 | 0.03767779527 | NDUFA12/ NDUFC2/ PET100/ UQCR11 |
| SYPH_DROP | hsa05417 | 6.017241379 | 0.04362867444 | CCL3/ CDC42/ HSP90B1/ NFKBIA |
| SYPH_DROP | GO:0022613 | 5.145257453 | 0.04988662897 | FAU/ FRG1/ NSA2/ RBIS/ RPL27/ RPL38/ RPS21/ SNRPF |

| Table S5 ｜ Reagent and resource | | |
|---|---|---|
| **Reagent or resource** | **Source** | **Identifier** |
| Antibodies | | |
| Anti-Human CD3 | Biolegend | Cat# 317306 |
| Anti-Human CD4 | Biolegend | Cat# 317408 |
| Anti-Human TCRα/β | Biolegend | Cat# 306706 |
| Anti-Human TCRγ/δ | Biolegend | Cat# 331208 |
| Anti-Human CD19 | Biolegend | Cat# 302206 |
| Anti-Human CD20 | Biolegend | Cat# 302304 |
| Anti-Human CD22 | Biolegend | Cat# 363508 |
| Anti-Human CD14 | Biolegend | Cat# 325604 |
| Anti-Human CD34 | eBioscience | Cat# 11-0349-42 |
| Anti-Human FcεR1α | eBioscience | Cat# 11-5899-42 |
| Anti-Human CD1a | Biolegend | Cat# 300104 |
| Anti-Human CD11c | Biolegend | Cat# 301604 |
| Anti-Human CD123 | Biolegend | Cat# 306014 |
| Anti-Human IFN-γ | eBioscience | Cat# 17-7319-82 |
| Anti-Human IFN-γ | eBioscience | Cat# 45-7319-42 |
| Anti-Human AREG | ebioscience | Cat# 17-5370-42 |
| Anti-Human CD56 | Biolegend | Cat# 318310 |
| Anti-Human CD56 | BD Biosciences | Cat# 564057 |
| Anti-Human TBX21 | eBioscience | Cat# 25-5825-82 |
| Anti-Human CD14 | Biolegend | Cat# 367123 |
| Anti-Human CD16 | Biolegend | Cat# 302011 |
| Anti-Human CD2 | Biolegend | Cat# 300221 |
| Anti-Human CD96 | Biolegend | Cat# 338415 |
| Anti-Human c-Jun | Cell SignalingTechnology | Cat# 15683S |
| Anti-Human CD107a | eBioscience | Cat# 12-1079-42 |
| Anti-Human CD107a | Biolegend | Cat# 328634 |
| Anti-Human CXCL8 | Biolegend | Cat# 511415 |
| Anti-Human NKp30 | Biolegend | Cat# 325233 |
| Anti-Human CD163 | Biolegend | Cat# 333621 |
| Anti-Human Phospho-STAT1 | eBioscience | Cat# 12-9008-42 |
| Biological Samples | | |
| PBMCs | The Second Hospital of Nanjing | https://njsech.com.cn/ |
| K562 | ATCC | Cat# CCL-243 |
| Chemicals, Peptides, and Recombinant Proteins | | |
| IL-2 | Novoprotein | Cat# C013 |
| IL-12 | Novoprotein | Cat# CI58 |
| IL-15 | Novoprotein | Cat# C016 |
| IL-18 | Novoprotein | Cat# CH29 |
| Dexamethasone Sodium Phosphate Injection | Shiyao YINHU Pharmaceutical Co., Ltd. | LOT# 1022122362 |
| Protein transport inhibitor | ebioscience | Cat# 00-4980-03 |
| Fixable Viability Dye | ebioscience | Cat# 65-0865-14 |
| JNK-IN-8 | MedChemExpress | Cat # HY-13319 |
| EasySep Human NK Cell Isolation Kit | STEMCELL | Cat#17955 |
| NK MACS Medium human | MACS | Cat# 130-114-429 |
| Software and Algorithms | | |
| FlowJo | FlowJo, LLC | https://www.flowjo.com/ |

| | | |
|---|---|---|
| GraphPad Prism | GraphPad Software, Inc | https://www.graphpad.com/scientific-software/prism/ |
